# In vivo genome editing of human hematopoietic stem cells for treatment of blood disorders by mRNA delivery

**DOI:** 10.1101/2024.10.28.620445

**Authors:** Saijuan Xu, Dan Liang, Qiudao Wang, Yan Cheng, Da Xie, Yang Gui, Haokun Zhang, Changrui Feng, Feiyan Zhao, Wendan Ren, Gongrui Sun, Yang Yang, Lin Li, Yongrong Lai, Bin Fu, Yuming Lu, Zi Jun Wang, Yuxuan Wu

## Abstract

*Ex vivo* autologous hematopoietic stem cells (HSCs) gene therapy provides promising new treatments for hematological disorders. However, current methods involve complex processes and chemotherapeutic conditioning, leading to limited accessibility for treatment and significant side effects. Here, we developed an antibody-free targeted lipid nanoparticles (LNPs) for mRNA delivery to HSCs *in vivo*, enabling efficient base editing of the HBG target in human HSCs to reactivate fetal hemoglobin in derived erythroid cells. Delivery of ABE8e/sgRNA mRNA with optimized structure LNPs achieves efficient *in vivo* base editing of HBG in transfusion-dependent β-thalassemia (TDT) patients derived HSCs engrafted in immunodeficient *NCG-X* mice, showing restored globin chain balance in erythroid cells. Our research indicated that utilizing LNPs for delivery of genome editor achieves efficient editing of endogenous genes of human HSCs. Notably, this non-viral delivery system eliminates the need for harvesting or mobilizing HSCs, providing a potent and one-time treatment potential for blood disorders like sickle cell disease (SCD) and TDT.

**Highlights:**

- Antibody-free targeted LNPs system achieves efficient mRNA delivery and base editing in human bone marrow cells including HSCs via one-time intravenous injection.
- Delivery of ABE8e/sgRNA mRNA with optimized LNPs achieves efficient *in vivo* base editing of HBG in TDT patients derived HSCs engrafted in immunodeficient *NCG-X* mice, showing restored globin chain balance in erythroid cells.
- LNPs efficiently deliver gene editing system to the BM for *in vivo* editing of human HSCs, providing preclinical evidence for the next generation non-viral gene therapy for blood disorders.

## Introduction

Autologous transplantation of gene-modified hematopoietic stem cells (HSCs) has been demonstrated as a curative treatment for hematopoietic disorders including β-thalassemia and sickle cell disease (SCD)^1–3^. This approach, while successful in numerous clinical trials and not requiring donor matching, potentially preventing graft-versus-host disease (GVHD), is still hindered by the challenge of acquiring sufficient high-quality HSCs for *ex vivo* manipulation^4^. Importantly, a pre-conditioning process is needed, typically involving chemotherapy or radiation, to eliminate the patient’s own HSCs and create space in the bone marrow (BM) niches for the gene-modified autologous HSCs^5,6^. However, this process comes with significant acute and chronic systemic toxicity, including infertility and secondary malignancies due to accumulated DNA damage^7^. In addition, due to the personalized patient-specific nature of autologous gene therapy, which requires specialized manufacturing centers, the cost of *ex vivo* gene therapy is often quite prohibitive for patients^8^. *In vivo* HSCs gene therapy approaches aim to simplify the gene therapy process by directly delivering RNA to HSCs within the body.

Current reported *in vivo* delivery strategies of HSCs utilize helper-dependent adenoviruses (HDAd) which require mobilization for efficient transduction followed by selection of edited cells through low doses of chemotherapy^9–10^. HDAd vectors have several notable disadvantages, including the induction of proinflammatory cytokine release upon administration, challenges posed by pre-existing immune responses against capsid proteins, high seroprevalence limiting their widespread use, limited tissue transduction capability for certain serotypes^11^. Using non-viral LNPs delivery systems for HSCs could overcome these limitations posed by adenovirus, enabling repeated dosing, and streamlining production. LNPs delivery system enables transient expression of gene-editing agents with low immunogenicity, comprised of biodegradable and non-toxic synthetic components, and potential for repeated use^12–15^. In previous studies, efficient gene editing using LNPs mRNA has been demonstrated in mice and non-human primates *in vivo*, and specific cell targeting through antibody modification on the LNPs surface^15–22^. Recent advancements have both successfully utilized antibodies against CD117 conjugated to LNP (CD117/LNP-mRNA) that, following a single intravenous injection, can deliver RNA (both siRNA and mRNA) to HSCs in mice *in vivo* ^19,22^. However, there are few studies that utilize established humanized mouse models to evaluate the therapeutic effect of *in vivo* gene-edited HSCs using LNPs.

Here, we developed an antibody-free conjugated LNPs for *in vivo* delivery of adenine base editor (ABE8e) mRNA and single-guide RNA (sgRNA) to human hematopoietic stem and progenitor cells (HSPCs). By designing and screening library of ionizable lipids featuring distinct head, linker and tail structures, we pinpointed LNP-028-ABE8e as the most effective for BM RNA delivery. In this study, we pioneered the delivery of ABE8e mRNA and sgRNA to human cells using the LNP-028-ABE8e, achieving efficient base editing at the HBG target to reactivate fetal hemoglobin (HbF, α_2_γ_2_) levels in red blood cells (RBCs). By optimizing the structural combination of head and tail moieties based on Library A, we identified Lipid-168 possess significantly enhanced delivery efficiency *in vivo*, which, following administration of LNP-168-ABE8e in a humanized β-thalassemia model, effectively base edited and significantly induced γ-globin (HBG1/2) expression in HSCs derived from TDT patients. Our findings highlight the significant therapeutic promise of this antibody-free conjugated LNPs mediated *in vivo* editing strategy for HSCs, effectively delivering gene-editing elements to HSCs in the BM. This robust preclinical evidence paves the way for an innovative approach to genetic therapies in blood disorders.

## RESULTS

### Identify novel point mutations inducing HbF expression in CD34^+^ HSPCs

For successful LNPs-mediated *in vivo* genome editing to treat β-hemoglobinopathy, optimizing the genome editing system with the most active base editor and sgRNA is essential due to the limited mRNA delivery efficiency of the LNPs system. ABE8e is currently the most efficient base editor, but screening for the optimal sgRNA to induce HbF activation in combination with it is pending. To identifying novel effective targets for ABE8e in the γ-globin region, we created HUDEP (human umbilical cord blood-derived erythroid progenitor)-2 cell lines stably expressing ABE8e harboring Cas9 nickase (nCas9) to mutate target sequences or dead Cas9 (dCas9) to bind target sequences without introduce DNA breaks^23,24^. HUDEP-2 is an immortalized human erythroblast cell line that can be induced to undergo terminal erythroid maturation and produce high levels of adult-type (β-globin) and low levels of fetal-type hemoglobin (γ-globin) ^25^. Whole-transcriptome RNA sequencing (RNA-seq) showed that stable expression of ABE8e altered neither gene expression in HUDEP-2 cells nor the capacity of the cells to undergo terminal erythroid maturation (Extended Data Figs. 1A, B and Supplementary Data Table 1). We realized that a tiling set of sgRNAs could reveal critical regions by disrupting almost all the sequences within the cis-modulating element. According to the presence of the *Streptococcus pyogenes* Cas9 (SpCas9) *NGG protospacer adjacent motif* (PAM) sequence^23^, we designed all possible sgRNAs (n=92) within 500 bp upstream of the human HBG promoter regions, which could approach saturation mutagenesis *in situ* at each genomic position (Supplementary Data Table 2). After intracellular HbF staining, cells were fractionated using fluorescence-activated cell sorting (FACS), followed by high-throughput DNA sequencing to compare sgRNA representation between HbF-high and HbF-low cells, resulting in the initial identification of 40 sgRNAs (Fig. 1a and Supplementary Data Table 3). As expected, known target sites were represented by sgRNA-22^26^, -35^27^, and -41^27^. Moreover, our screening process also revealed novel targeting sites, especially sgRNA-25 and sgRNA-26, which induced significant HbF expression. To validate the results, four sgRNAs (sgRNA-25, -26, -35 and -41) were selected to further confirm their efficacy in primary erythroblasts (Fig. 1b and Supplementary Data Table 4).

**Fig.1.**
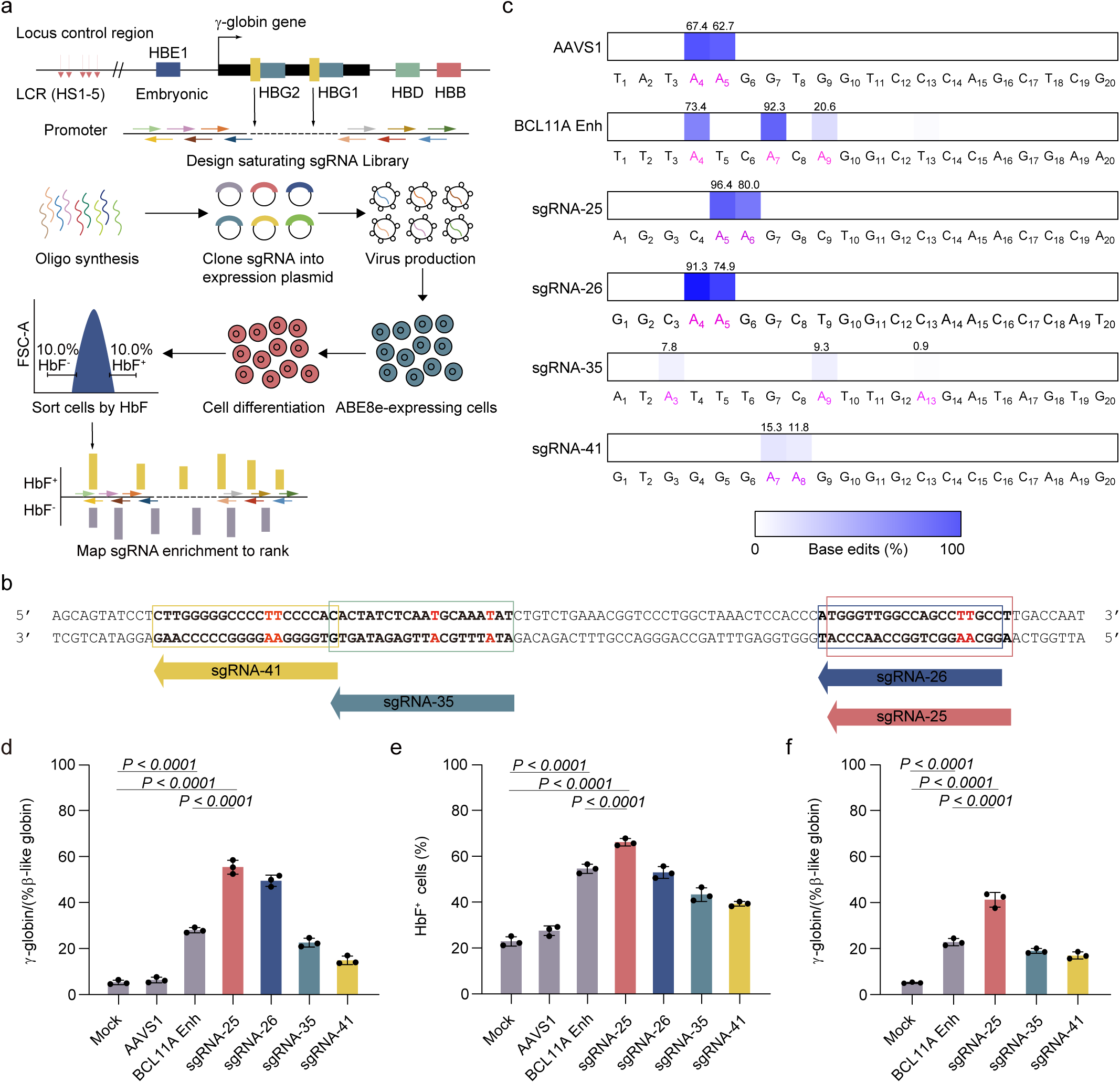
| High-throughput screening of the HBG promoter using ABE8e to identify efficient sgRNA in CD34^+^ HSPCs that elevate HbF levels. **a**, Workflow of sgRNAs screening showing library synthesis, delivery, and analysis. Designing sgRNAs saturation libraries with NGG PAM motifs within 500 bp upstream of the HBG promoter, to screen for efficient induction of HbF expression in HUDEP-2 cell lines expressing ABE8e. The representative HbF FACS sorting strategy is used to separate high-HbF and low-HbF cells for sgRNA screening. **b,** Schematic sgRNAs of the human γ-globin proximal promoter within β-like globin gene locus. The position of the sgRNA is indicated by an arrow. **c,** Overall base editing frequencies were determined for various sgRNAs in CD34^+^ HSPCs 5 days after ABE8e RNP electroporation. Positions exhibiting measurable base editing were indicated in purple. Base editing quantification was performed using *EditR*. **d,** Percentage of γ-globin mRNA (γ/γ + β) expression by RT-qPCR analysis in erythroid cells *in vitro* differentiated from RNP edited CD34^+^ HSPCs. RT-qPCR, quantitative reverse transcription polymerase chain reaction. **e,** Percentage of HbF^+^ in erythroid cells *in vitro* differentiated from ABE8e RNP edited CD34^+^ HSPCs with various gRNAs, measured by flow cytometry analysis. **f,** γ-globin protein levels after *in vitro* erythroid maturation from ABE8e RNP edited CD34^+^ HSPCs, measured by HPLC analysis. Data are representative of three biologically independent replicates. All statistical significances in the figures were analyzed using one-way ANOVA with Dunnett’s multiple comparisons test, and data represent mean ± SD, with *p*-values noted where appropriate.

Electroporation was employed to deliver ABE8e proteins complexed with chemically modified synthetic sgRNAs (ABE8e/sgRNA ribonucleoprotein complex, RNP) specifically targeting the HBG promoter in human CD34^+^ HSPCs obtained from a healthy donor^28^. The +58 BCL11A erythroid enhancers, previously validated for their ability to reactivate HbF expression, were utilized as a positive control for RNP editing, while the *AAVS1* locus served as the functionally neutral control^29^. sgRNA-25 target exhibited significant base editing activity, resulting in an increase in γ-globin mRNA levels and an elevated proportion of HbF^+^ cells in erythroid cells (Fig. 1c-e). Therefore, due to its stronger impact on γ-globin expression and effective base editing, sgRNA-25 was prioritized for subsequent experiments. Flow cytometry analysis revealed that a substantial proportion of the edited cells treated with ABE8e RNP underwent enucleation during terminal differentiation without affecting erythroid cell differentiation (Extended Data Fig. 1C,D). Bulk-edited cells from the sgRNA-25 group were FACS sorted based on HbF intracellular levels at the terminal differentiation stage, enabling a comparison of T-C conversion between HbF-high and HbF-low cells, which revealed a significantly higher base editing T-C conversion in HbF-high cells compared to HbF-low cells (90.3% versus 4.8%, Extended Data Fig. 1E). Further confirmation of the reactivation of Aγ and Gγ-globin chains at the sgRNA-25 target was provided in erythroid cells through reversed-phase high-performance liquid chromatography (RP-HPLC) analysis (Fig. 1f and Extended Data Fig. 1F).

Through comprehensive sequence analysis, our study investigated the impact of base editing on epigenetic regulation, identifying a new Sp1 (Sp1 transcription factor) binding motif at the target site (Extended Data Fig. 2A). Our analysis revealed increased chromatin accessibility, significant enrichment of Sp1 binding sites, heightened peaks, and curve analysis in the HBG gene region of edited CD34^+^ HSPCs (Extended Data Fig. 2B-F). A recent study suggested that introducing mutations similar to HPFH (−123T > C and -124T > C) might induce the expression of γ-globin by creating a new binding site for KLF1^30^. Transcription factors from the Sp1-like and Krüppel-like factors family (Sp1/KLFs) play essential roles in transcriptional regulation, recognizing CACCC/GC boxes in gene promoters^31^. It has been reported that *Sp1* can activate the expression of certain genes by recruiting histone deacetylase 1 (HDAC1) while simultaneously repressing the expression of other genes^32^. Studies indicate that *Sp1* acts as an inhibitor of β-globin gene transcription during erythroid terminal differentiation. Its phosphorylation and release allow erythroid-specific FKLF2 or KLF to interact with other erythroid-specific transcription factors, thereby initiating β-globin gene transcription^33^. Our research results confirm that *Sp1* can bind to the HBG promoter region after mutations -123 T > C and -124 T > C, with KLF1 potentially acting as the main activating factor, and other Sp/KLF transcription factors may also contribute to the activation of the HBG gene. Future studies need to further clarify the specific mechanisms underlying the upregulation of γ-globin expression induced by these mutations.

### Base editing of -123 T > C and -124 T > C induces therapeutic HbF expression in β**-thalassemia patient-derived CD34^+^ HSPCs**

To further examine the effectiveness of sgRNA-25 base editing in the context of disease pathobiology, we evaluate the therapeutic HbF induction through ABE8e RNP editing in primary HSPCs obtained from individuals with β-thalassemia (Fig. 2a). The base editing efficiency of RNP with sgRNA-25 was high in β-thalassemia HSPCs, with editing rates of 86.9% at A5 and 71.0% at A6 (Fig. 2b). Additionally, we observed potent induction of γ-globin and show restored globin chain balance in base edited erythroid cells derived from β-thalassemia HSPCs (Fig. 2c). The sgRNA-25 bulk base edited erythroid progeny demonstrated 47.0% HbF as compared to a baseline of 21.9% in unedited β-thalassemia cells (Fig. 2d). We hypothesized that improving the imbalance of globin chains, which underlies the pathophysiology of β-thalassemia, would enhance terminal erythroid maturation. As anticipated, the sgRNA-25-edited samples demonstrated a higher frequency of enucleation in terminal erythroid cells compared to the unedited cells (Fig. 2e). The results provide evidence for the successful introduction of therapeutically relevant base edits in HSPCs derived from β-thalassemia patients.

**Fig.2.**
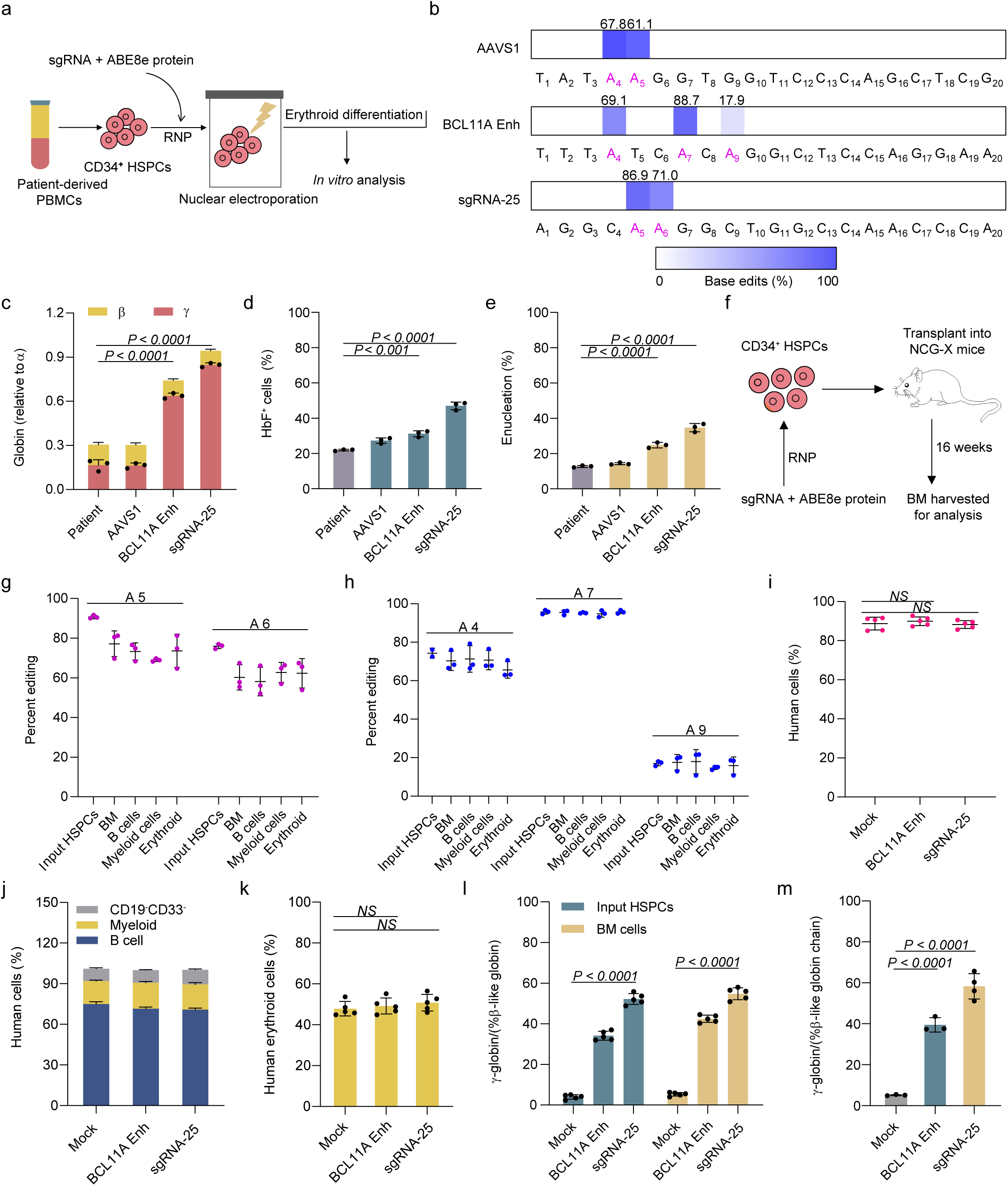
| Efficient base editing to induce HbF expression in CD34^+^ HSPCs from β-thalassemia patients and in immunodeficient mice. **a,** Schematic representation of steps involved in base editing of β-thalassemia CD34^+^ HSPCs. CD34^+^ HSPCs from β-thalassemia were nucleofected using Lonza system with ABE8e protein and respective sgRNA on day 2 of expansion. The β^0^/β^0^ thalassemia cells originate from a patient with CD41-42 (-TCTT) and IVS-II-654 (C-T) mutations. **b,** Deep sequencing analysis was used to measure base editing efficiency in β^0^β^0^ CD34^+^ HSPCs electroporated with ABE8e RNP after 5 days. **c,** β-like globin expression by RT–qPCR normalized by α-globin, measured in ABE8e RNP edited erythroid progeny after *in vitro* erythroid differentiation. *n*=3 replicates from independent electroporations. **d,** Flow cytometry analysis showed the percentage of HbF^+^ cells derived from edited β^0^β^0^ CD34^+^ HSPCs after *in vitro* erythroid differentiation. *n*=3 replicates from independent electroporations. **e,** Flow cytometry analysis showed the level of enucleation of RBCs derived from β^0^β^0^ CD34^+^ HSPCs edited after *in vitro* erythroid differentiation. *n*=3 replicates from independent electroporations. **f,** The RNP composed of ABE8e protein complexed with sgRNA was electroporated into human CD34^+^ HSPCs. After 24 h post-electroporation, 0.8 million edited cells were obtained, and unedited cells were transplanted into *NCG-X* mice via intra-orbital injection. After 16 weeks transplantation, BM cells were harvested and analyzed using flow cytometry to assess human HSC reconstitution, editing efficiency, and γ-globin expression levels. Non-electroporated cells were transplanted as controls. **g,** Mouse BM cells was analyzed for human cell chimerism by flow cytometry of 16 weeks after transplantation, defined as %hCD45^+^/(%hCD45^+^ + %mCD45^+^) cells. Each symbol represents a mouse, and there were a total of n=3 mice. **h,** BM cells collected after transplantation was analyzed by flow cytometry for multilineage reconstitution (calculated as percentage of hCD45^+^ cells). **i,** BM cells collected after transplantation was analyzed by flow cytometry for CD235a^+^ erythroid cells (calculated as percentage of mCD45^−^hCD45^−^ cells). **j,** Base editing frequencies at the HBG promoter were determined in the input HSPCs before transplantation and in engrafted B cells, myeloid cells, erythroid cells, and BM cells after transplantation, using deep sequence analysis. Each dot represents one mouse that underwent engraftment, and the mean value for each group is presented. **k,** Base editing frequencies at the *BCL11A* enhancer were determined in the input HSPCs before transplantation and in engrafted B cells, myeloid cells, erythroid cells, and BM cells after transplantation, using deep sequence analysis. Each dot represents one mouse that underwent engraftment, and the mean value for each group is presented. **l,** β-like globin expression (γ) by RT-qPCR in the input HSPCs before transplant and in human cells from BM of engrafted mice. **m,** Quantification of γ-globin protein levels (γ/γ+β), as assessed by HPLC. All statistical significances in the figures were analyzed using one-way ANOVA with Dunnett’s multiple comparisons test, and data represent mean ± SD, with *p*-values noted where appropriate. *NS*, not significant.

### Highly efficient base editing of HSCs induces HbF expression *in vivo*

To assess the effects of HBG promoter editing on HSCs, we transplanted modified human CD34^+^ HSPCs into immunodeficient *NCG-X* mice, since they support not only myeloid and lymphoid, but also erythroid engraftment. We further conducted *in vivo* studies by infusing an equal number of viable unedited or edited CD34^+^ HSPCs from healthy donors into highly immunodeficient NCG-X mice, which support human hematopoietic engraftment without conditioning therapy (Fig. 2f) ^34–36^. After 16 weeks, we observed that recipients of both edited and unedited CD34^+^ HSPCs showed comparable levels of human chimerism in BM cells (Fig. 2g). Additionally, the engraftment levels of human lymphoid, myeloid, and erythroid cells within the BM were similar for both groups (Fig. 2h,i). Flow cytometry was utilized to isolate donor-derived mononuclear cells, B cells, myeloid cells, and erythroblasts from mouse BM, allowing for the examination of editing frequency in each cell group (Extended Data Fig. 3A,B). Editing efficiency at *BCL11A* target loci in the engrafted BM cells was virtually equal to the input cell populations whereas editing efficiency at sgRNA-25 target sequence was modestly decreased in BM compared with the input cells (Fig. 2j,k). In human erythroid cells derived from BM of engrafted mice, a significant induction of γ-globin expression was observed, with levels increasing from 5.3% to 54.8% after editing, consistent with input HSPCs (Fig. 2l). RP-HPLC analysis confirmed reactivation of γ-globin chains in edited erythroid cells sorted from BM using immunomagnetic beads (Fig. 2m). Thus, these results indicate that the engraftment and differentiation potential of transplanted HSPCs is not altered by ABE8e-mediated efficient base editing.

### Rational design of ionizable LNPs for efficient HbF induction by base editing

Therapeutic *in vivo* delivery of gene editing agents by LNPs as a versatile and promising solution for blood disorders, surpassing the limitations of *ex vivo* methods^13,14,16,17,19,22,37–40^. Currently, most of the work focuses on adjusting the amino head, linker, and hydrophobic tail structure of ionizable lipids to improve escape efficiency, transfection efficiency, and safety profile^13,21,41–43^. To improve the *in vivo* delivery efficiency of LNPs for BM cells, we systematically designed and synthesis three distinct ionizable lipid libraries each showcasing a unique linker structure. Library A employed an α-acylamino acyl amide linker structure, Library B utilized (2-(aminomethyl) propane-1,3-diol) as the linker structure, and Library C utilized (3-hydroxy-2-(hydroxymethyl) propyl) carbamothioic S-acid as the linker structure (Fig. 3a). Within Library A, 35 unique ionizable lipids were synthesized by maintaining the linker structure (purple section) while simultaneously modifying head structures and three independently adjustable tail structures. Similarly, Libraries B and C generated 32 and 25 unique ionizable lipid structures, respectively, by modifying the head and tail structures. To screen for ionizable lipids with high-efficiency delivery to BM from three libraries *in vivo*, we opted for the use of PCSK9, which has been previously reported or validated, as the target gene for LNP-mRNA delivery system^16,17^. The LNPs encapsulating ABE8e mRNA and sgRNA-PCSK9 (LNP-ABE8e-PCSK9) were injected via the tail vein into wild-type C57BL/6 mice at a dose of 2.0 mg/kg, and the base editing frequency in BM cells was measured after second-dose injection (Supplementary Data Table 7). Following the administration of LNP-ABE8e, we present the chemical structures of ionizable lipids from various libraries along with their editing efficiency of BM cells (Extended Data Fig 4A,B, 5A,B and 6A,B). Specifically, the editing efficiency of LNP-028-ABE8e-PCSK9 from Library A was significantly higher *in vivo* compared to the highest performers from Library B (LNP-217-ABE8e-PCSK9) and Library C (LNP-306-ABE8e-PCSK9). Furthermore, the editing efficiency significantly increased after re-administration, consistent with *in vitro* validation results (Fig. 3b,c). Together, these findings prompted us to continue further *in vivo* experiments in this study with Lipid-028.

**Fig.3.**
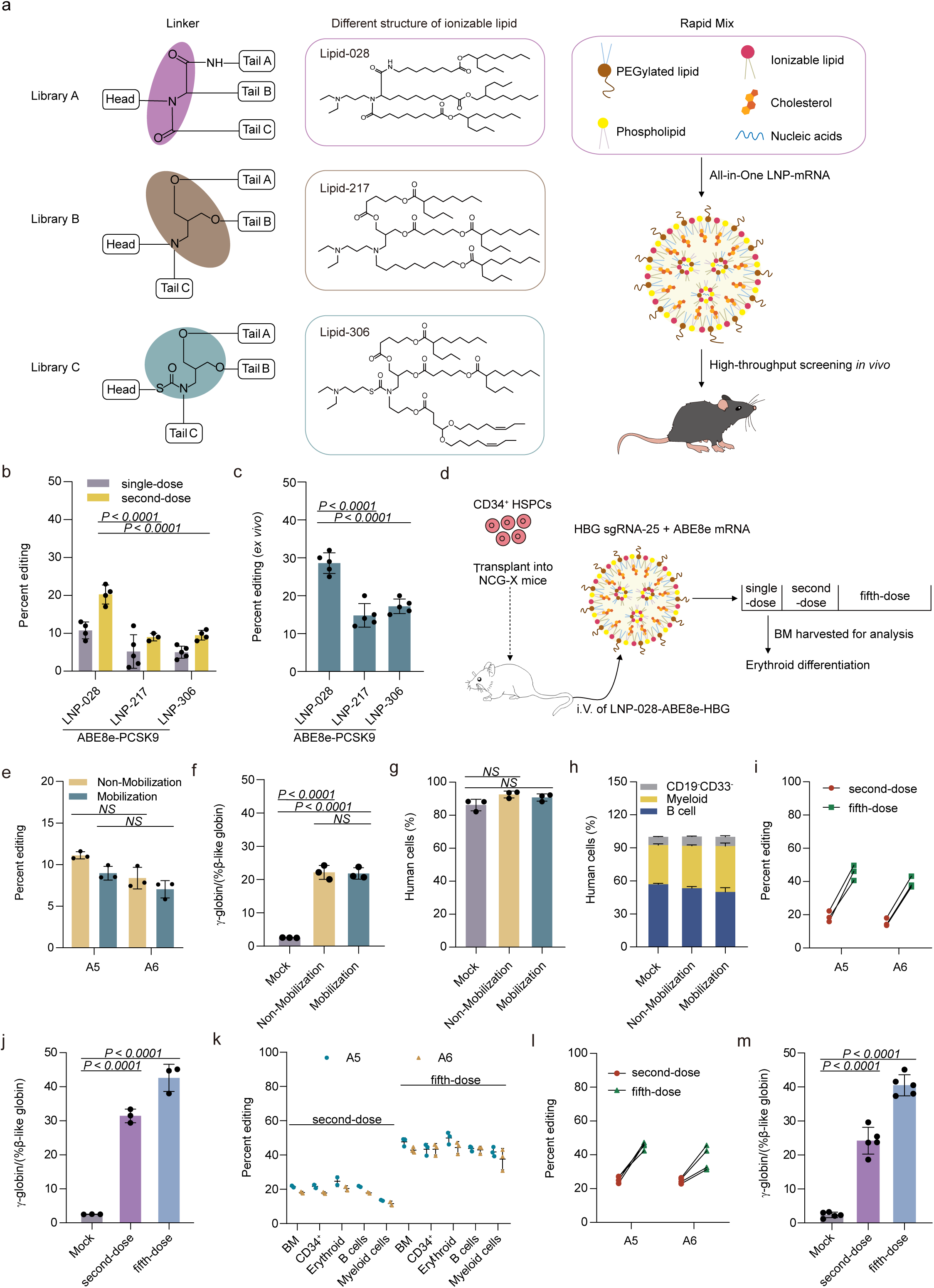
| Efficient *in vivo* base editing by LNP-mediated mRNA/sgRNA delivery in humanized mice. **a,** The illustration depicts the integration of four components (PEGylated lipids, phospholipids, ionizable lipids, and cholesterol) using a microfluidic mixing device to prepare LNPs mRNA. Libraries A, B, and C were designed by modifying the structure of ionizable lipid heads group, linkers, and tails to enhance their efficiency in targeting BM through *in vivo* animal experiments. **b,** LNPs were formulated with distinct ionizable lipid candidates, encapsulated ABE8e mRNA and sgRNA-PCSK9, and intravenously injected into C57 mice at a dose of 2 mg/kg. Fourteen days after the LNP-ABE8e-PCSK9 initial injection, a second dose was administered, and BM cells was collected for analyzing the editing efficiency. **c,** Percent editing at the target base (A6) of sgRNA-PCSK9 was assessed after treating *in vitro* transfected BM cells with different candidate LNPs encapsulating ABE8e and sgRNA-PCSK9 for 3 days. **d,** Intravenous (i.v.) administration workflow employing co-encapsulation of LNP-028-ABE8e-HBG. After 16 weeks of transplanting CD34^+^ HSPCs from healthy donors into *NCG-X* mice, LNP-028-ABE8e-HBG was intravenously injected into the humanized mice at a dose of 2 mg/kg every two weeks for a total of 5 injections. After two weeks following the single-dose, second-dose, and fifth-dose of LNP-028-ABE8e-HBG injection, BM cells were collected for analysis. **e,** After LNP-028-ABE8e-HBG single-dose injection, the average base editing efficiency in BM cells of mobilized treatment and non-mobilized treatment groups of engrafted mice. **f,** β-like globin expression (γ) in the BM cells of engrafted mice after LNP-028-ABE8e-HBG treated measured by RT-qPCR. **g-h,** After administering LNP-028-ABE8e-HBG for base editing, the BM of engrafted mice was analyzed using flow cytometry to assess human cell chimerism **(g)** and multilineage reconstitution **(h)**. **i,** Percentage of base editing after second-dose and the fifth-dose of LNP-028-ABE8e-HBG administered intravenously via tail vein injection was assessed in the BM cells of engrafted mice. **j,** The β-like globin expression (γ) in the BM cells of engrafted mice after LNP-028-ABE8e-HBG base editing was measured by RT-qPCR. **k,** Overall editing frequency engrafted HSPCs, B cells, myeloid cells, CD34^+^, erythroid cells, and BM cells of engrafted mice after LNP-028-ABE8e-HBG base editing, measured by deep sequence analysis. **l,** Base editing by deep sequence analysis at A5 and A6 position from LNP-028-ABE8e-HBG edited BM cells after 16 weeks of secondary transplantation. Edited BM cells were collected from humanized mice following two and five administrations of LNP-028-ABE8e-HBG, and subsequently transplanted into *NCG-X* mice at a dose of 1 million cells per recipient. **m,** γ-globin expression in LNP-028-ABE8e-HBG edited BM cells after 16 weeks of secondary transplantation. All statistical significances in the figures were analyzed using one-way ANOVA with Dunnett’s multiple comparisons test, and data represent mean ± SD, with *p*-values noted where appropriate. *NS*, not significant.

To assess the feasibility of using this platform for therapeutic human HSPCs editing, we used LNPs encapsulating ABE8e mRNA and sgRNA-25 targeted to the HBG promoter (LNP-028-ABE8e-HBG). We first transplanted CD34^+^ HSPCs from healthy donors into immunodeficient NCG-X mice after 16 weeks, followed by second intravenous injections of LNP-028-ABE8e-HBG at a dose of 2.0 mg/kg (Fig. 3d). Our method involves subcutaneous injection of G-CSF/plerixafor to effectively mobilize HSCs from the BM to the peripheral blood stream, followed by intravenous injection of a BM-targeted LNPs delivery system, potentially enhancing the editing efficiency of HSC cells in the BM^44^. After LNP-028-ABE8e-HBG infusion, we observed editing efficiencies of 11.0% (A5) and 8.4% (A6) in the non-mobilized group, versus 9.0% (A5) and 7.0% (A6) in the mobilized group (Fig. 3e and Extended Data Fig. 7A). We observed a significant induction of γ-globin, with mean levels reaching 22.2% in the non-mobilized group and 21.8% in the mobilized group of edited human cells from BM (Fig. 3f). We found that mice of non-mobilized edited group and mobilized edited group exhibited similar levels of human chimerism in the BM cells after LNP-028-ABE8e-HBG injections compared to the mock group (Fig. 3g,h and Extended Data Fig. 3). These results suggest similar editing efficiency and levels of γ-globin induction in human cells from BM, regardless of mobilization status, following LNP-028-ABE8e-HBG injections. Consequently, we opted for the non-mobilized approach for further *in vivo* treatment.

Next, we aimed to enhance both the editing efficiency of HBG promoter and the expression levels of γ-globin by increasing the number of injections. After 16 weeks post-transplantation of CD34^+^ HSPCs from healthy donors into NCG-X mice, LNP-028-ABE8e/sgRNA-25 was intravenously administered at a dose of 2.0 mg/kg every two weeks, totaling five injections. We observed that the second (2nd dose) and the fifth doses significantly increased both the editing rates and the γ-globin mRNA levels in human bone marrow cells of the engrafted mice following LNP-028-ABE8e-HBG injections (Fig. 3i,j and Extended Data Fig. 7B). Our findings revealed that mice transplanted with CD34^+^ HSPCs from healthy donors and subsequently injected with LNP-028-ABE8e-HBG to edit the cells exhibited similar levels of human chimerism in the BM cells compared to the mock group (Extended Data Fig. 8A). Furthermore, it was noted that recipients of second-dose edited, fifth-dose edited, and unedited CD34^+^ HSPCs showed similar levels of engraftment for human lymphoid, myeloid, and erythroid cells in the BM cells from engrafted mice (Extended Data Fig. 8B,C), suggesting that administration of LNP-028-ABE8e-HBG into humanized mice did not damage human HSPCs or perturb their reconstitution. FACS analysis in human cells from BM showed similar base editing frequencies in each hematopoietic lineage after LNP-028-ABE8e-HBG injections (Fig. 3k). In human erythroid cells from the BM of engrafted mice, we observed effective editing rates and robust induction of γ-globin at similar levels as human BM cells after LNP-028-ABE8e injections (Extended Data Fig. 8D,E). After *in vitro* differentiation edited BM of engrafted mice into human erythroid cells after LNP-028-ABE8e-HBG injections, we observed efficient enucleation (Extended Data Fig. 8F). The hallmark features of HSCs are the capacity for multi-lineage hematopoietic repopulation and self-renewal. Secondary transplantation of edited BM cells from engrafted mice showed a comparable base editing frequency to primary transplantation (Fig. 3l). In secondary BM recipients through LNP-028-ABE8e-HBG base editing, the activation level of γ-globin in BM cells remained comparable to that observed after the primary transplantation (Fig. 3m and Extended Data Fig. 9A). The edited HSCs demonstrated the ability to support secondary transplantation at a comparable level of multilineage human hematopoietic engraftment, in line with the activity of unedited hematopoietic stem cells (Extended Data Fig. 9B,C). FACS analysis of BM cells revealed similar and effective base editing frequency in each hematopoietic lineage, mirroring the outcomes of the primary transplantation (Extended Data Fig. 9D). These data suggest the feasibility to produce base edits by LNP-028-ABE8e-HBG injection in long-term hematopoietic stem cells (LT-HSCs). This suggests the possibility of generating base edits in LT-HSCs without compromising their engraftment and reconstitution capabilities after injections of LNP-028-ABE8e-HBG.

### *In vivo* screening of the ionizable lipids for efficient mRNA delivery to BM

To further explore more efficient and safer LNPs for mRNA delivery to the BM, we conducted a secondary screening of Library A and proceeded to systematically design and synthesize 50 different ionizable lipids by altering the head and tail of the lipids (Extended Data Fig. 10 and Supplementary Data Table 8). Following the administration of LNP-ABE8e-PCSK9, we identified noteworthy discrepancies in the editing efficiency among various ionizable lipids (Fig. 4a). As a representative, LNP-168-ABE8e-PCSK9 were presented chemical structure of Lipid-168 and imaged with cryo–transmission electron microscopy (Fig. 4b). Among these, Lipid-168 stood out for its superior efficiency in BM delivery, as evidenced by the editing efficiency of LNP-168-ABE8e-PCSK9 increasing to 48.5% with second-dose injections, compared to 19.7% for LNP-028-ABE8e-PCSK9 (Fig. 4a,c). Further analysis of chemical structures highlighted ionizable lipids sharing consistent linker and diethylamino head groups but varying tail structures, emphasizing the critical role of tail modifications in enhancing BM delivery.

**Fig.4.**
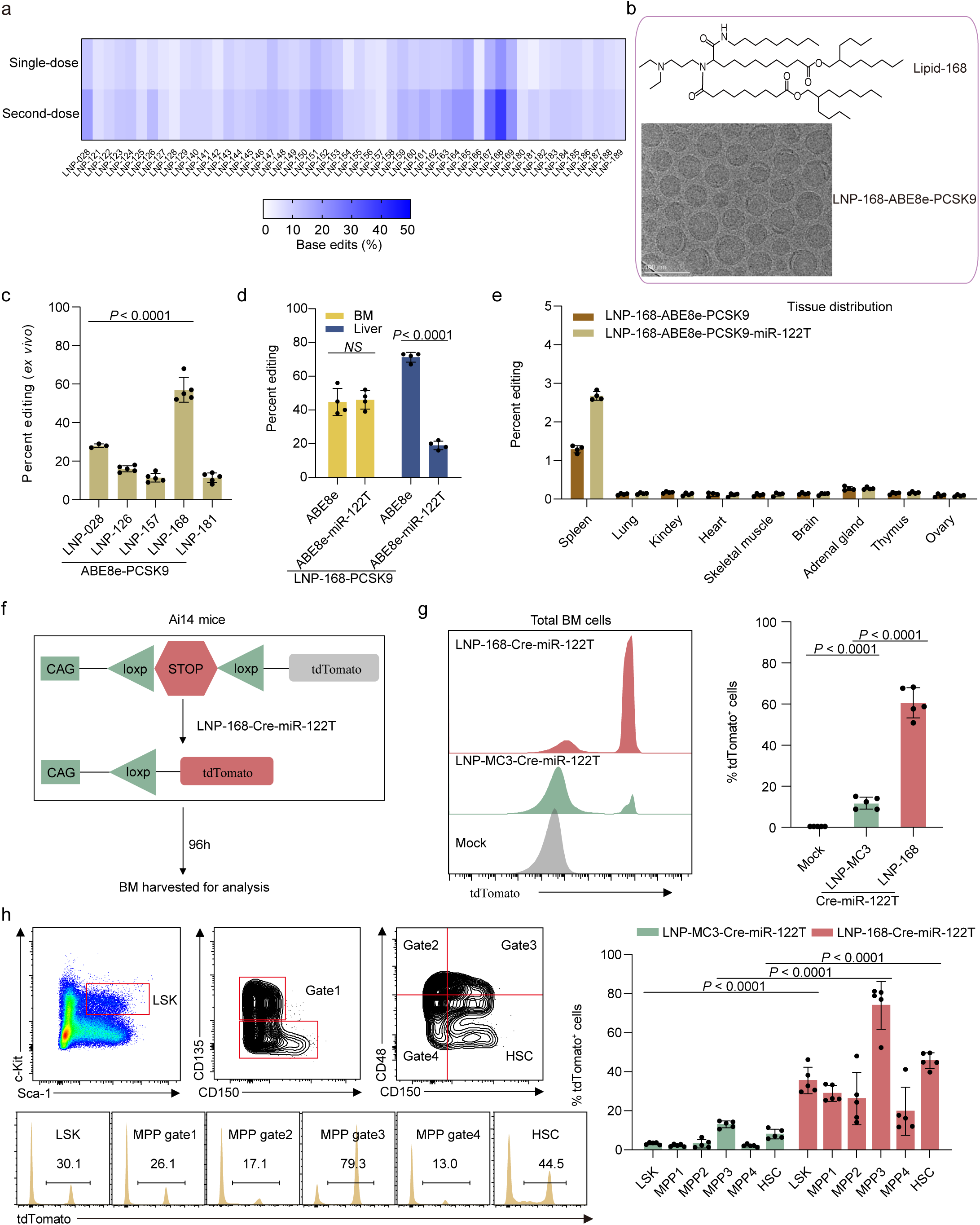
| *In vivo* LNP-168-Cre editing of BM HSPC in Ai14 mice leads to efficient tdTomato^+^ expression. **a,** Heat map image showing *in vivo* editing efficiency in Bm cells. C57 mice were treated with 50 different LNP packages containing ABE8e mRNA and sgRNA-PCSK9 at a dose of 2 mg/kg, and editing efficiency was measured 14 days post-treatment. A second injection at the same dosage was administered after another 14 days, followed by the collection of BM cells 7 days later for analysis of editing efficiency. **b,** Representative ionizable lipids with a chemical structure similar to that of Lipid-168 were used to prepare LNP-168-ABE8e-PCSK9, which was then analyzed for size using cryo-transmission electron microscopy. **c,** Percent editing at the target base (A6) of sgRNA-PCSK9 was assessed after treating *in vitro* transfected BM cells with different candidate LNP-ABE8e-PCSK9 for 3 days. Each symbol represents a mouse, and there were a total of n=4 mice. **d,** C57 mice were administered LNP-168 encapsulated ABE8e or ABE8e-miR-122T mRNA and sgRNA-PCSK9, via tail vein injection at a dose of 2 mg/kg. After an initial injection, LNP-168-ABE8e-PCSK9 or LNP-168-ABE8e-miR-122T-PCSK9 was second-dosed with the same dose after 14 days, and BM along with liver cells were collected to assess editing efficiency. **e,** The tissue distribution of base editing was observed in C57 mice in tissues other than the liver and BM cells two weeks after LNP-168-ABE8e-PCSK9 or LNP-168-ABE8e-miR-122T-PCSK9 treatment. **f,** Schematic of the Ai14 transgenic mouse LoxP-flanked stop cassette preventing the transcription of tdTomato. LNP-168-Cre-miR-122T and LNP-MC3-Cre-miR-122T were intravenous injected to mice at dose of 1 mg/kg, leading to tdTomato expression in the cell. Four days after injection, tdTomato fluorescence was analyzed. **g,** Flow cytometric analysis of tdTomato^+^ fluorescence induction in total BM cells following intravenous injection of LNP-Cre-miR-122T. **h,** Representative flow cytometry schematics show the percentage of tdTomato^+^ cells among BM HSPCs subsets after single LNP-Cre-miR-122T injections. Histograms illustrate tdTomato^+^ fluorescence, and summary bar graphs present the percentage of tdTomato^+^ cells among HSPCs subsets. All statistical significances in the figures were analyzed using one-way ANOVA with Dunnett’s multiple comparisons test, and data represent mean ± SD, with *p*-values noted where appropriate. *NS*, not significant.

### Liver editing was further reduced by adding miR-122T component to the editor mRNA

Despite LNP-168-ABE8e-PCSK9 showing effective editing efficiency in BM cells, there is still considerable editing observed in the liver because of the liver-tropic property of LNPs (Fig. 4d and Extended Data Fig. 11 A, B)^22,45,46^. Previous research demonstrated that microRNA-122 (miR-122) expression in vertebrate liver cells reduces target gene expression in hepatocytes^47^. We integrated miR-122 target sequences (miR-122T) into the 3’-untranslated region (UTR) of luciferase (Luc) mRNA and found significantly decreased expression of luciferase in the liver (Extended Data Fig. 11 A and Supplementary Data Table 9, 10)^47^. The above results suggested that miR-122T may also be used to decrease gene editing caused by LNP-168-ABE8e-PCSK9 in the liver. The results shown that LNP-168-ABE8e-PCSK9-miR-122T produced BM editing rate was comparable to that of the LNP-168-ABE8e-PCSK9 group, while the editing rate in the liver decreased significantly from 71.0% to 19.0% (Fig. 4d). Analysis across various tissues revealed lower levels of editing in the spleen and adrenal glands, with minimal detection of editing in other tissues (Fig. 4e).

Furthermore, a genetically engineered Cre/LoxP Ai14 mouse was utilized to investigate mRNA delivery to BM cell subtypes, employing LNPs-mediated Cre recombinase delivery to activate tdTomato fluorescence expression in edited cells upon stop cassette removal^48^. LNPs encapsulating mRNA encoding Cre-miR-122T recombinase (LNP-Cre-miR-122T) was administered to Ai14 mice via tail vein injection at a dose of 1 mg/kg, and liver-targeted LNPs (MC3) were injected at the same dose as the control group (Fig. 4f)^16,49^. The results showed that LNP-168-Cre-miR-122T achieved significantly higher tdTomato^+^ cells expression compared to LNP-MC3-Cre-miR-122T (Fig. 4g). To comprehensively assess tdTomato^+^ expression in HSPC subsets, we categorized LSK cells into multipotent progenitors (MPP) and HSCs^50^. Comparative analyses across various cell populations in the BM revealed higher tdTomato^+^ expression in LSK cells, MPPs, and HSCs treated with LNP-168-Cre-miR-122T compared to LNP-MC3-Cre-miR-122T at equivalent doses (Fig. 4h). Examination of distinct BM cell populations (T cells, Gr1, CD11b^+^, and B cells) demonstrated that LNP-168-Cre-miR-122T exhibited greater TdTomato fluorescence than LNP-MC3-Cre-miR-122T, indicating that LNP-168 has a stronger BM cell delivery capability compared to traditional LNPs (Extended Data Fig. 12A). Moreover, we performed polymerase chain reaction (PCR) to verify LNP-168-Cre-miR-122T mediated genomic deletion in BM DNAs (Extended Data Fig. 12B). The results show that a single dose of LNP-168-Cre can efficiently deliver functional mRNA to HSPCs *in vivo*, emphasizing its promising therapeutic potential for blood disorders through the using of optimized LNPs.

### LNP-168-ABE8e-HBG mediated *in vivo* therapeutic base editing of HBG promoter in CD34^+^ HSPCs derived from **β**-thalassemia patients

We then evaluated LNP-168-ABE8e-HBG mediated *in vivo* base editing of HSPCs on the pathobiology of the disease within a clinically significant model. We initially transplanted CD34^+^ HSPCs from β-thalassemia patients into immunodeficient *NCG-X* mice after 16 weeks, followed by second intravenous injections of LNP-168 encapsulating ABE8e mRNA and sgRNA-25 at a dose of 1.0 mg/kg (Fig. 5a and Supplementary Data Table 9). We observed that the average base editing frequency at the target HBG promoter achieved 42.6% with two doses of intravenous infusions of LNP-168-ABE8e-HBG, accompanied by potent induction of γ-globin in erythroid cells (Fig. 5b,c). After treatment with LNP-168-ABE8e-HBG, there was no significant difference in the proportion of human CD45 cells in the BM compared to the control group, indicating that LNP-168-ABE8e-HBG does not adversely affect β-thalassemia HSPCs (Fig. 5d). In addition, the BM proportions of lymphoid, myeloid, and erythroid cells in the LNP-168-ABE8e-HBG group were also comparable to those in the mock group, suggesting that treatment with LNP-168-ABE8e-HBG does not affect the lineage reconstitution of β-thalassemia HSPCs (Fig. 5e,f). In human erythroid cells derived from the BM, we observed robust induction of γ-globin after editing (mean increase from 0.14 to 0.60 relative to α-globin) and improved alpha-to-non-alpha globin chain balance (Fig. 5g). After LNP-168-ABE8e-HBG base editing, higher enucleation efficiency, increased cell size, and a more circular shape in enucleated erythroid cells were observed, indicating that *in vivo* editing by LNP-168-ABE8e-HBG led to the restoration of erythroid cell morphology of β-thalassemia samples (Fig. 5h-j). In conclusion, our results demonstrated that LNP-168-ABE8e-HBG allows for efficient *in vivo* base editing, thereby inducing sufficient γ-globin expression to rescue the hematological phenotype of β-thalassemia cells.

**Fig.5.**
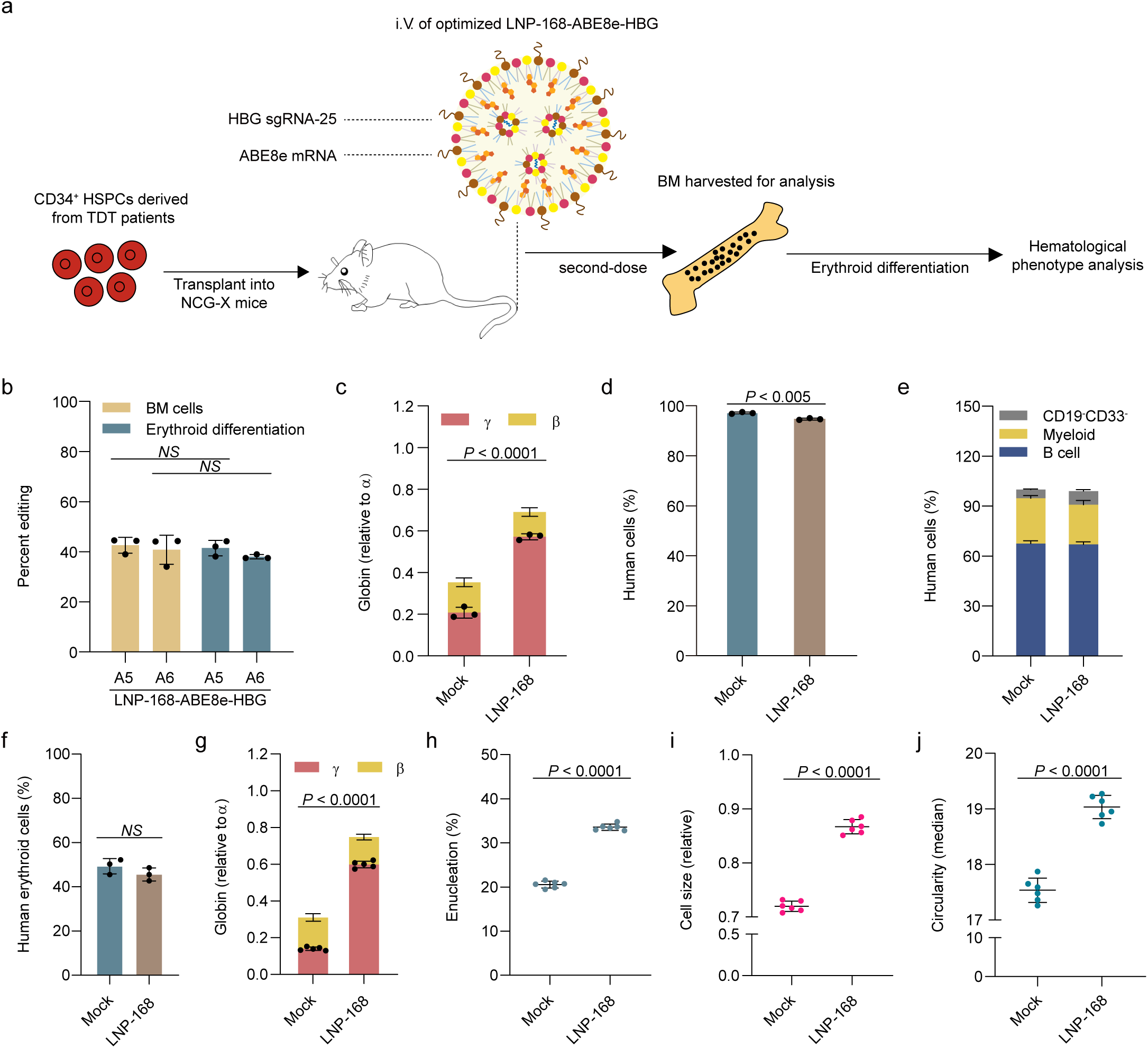
| LNPs-mediated ABE8e mRNA delivery leads to therapeutic base editing in β-thalassemia patient CD34^+^ HSPCs. **a**, After transplanting CD34^+^ HSPCs from β-thalassemia patients into *NCG-X* mice for 16 weeks, followed by intravenous injection of LNP-168 encapsulated ABE8e mRNA and sgRNA-25 at a dose of 1 mg/kg, with a second injection at the same dose after 14 days. BM cells from mice were collected 14 days after the second-dose injection for analysis. The β^+^/β^0^ patient cells we used to be derived from a single patient with CD41-42 (-TCTT) and β-28(A-T)) mutations. **b,** Evaluation of base editing efficiency in patient cells and *in vitro* differentiated erythroid cells from BM of engrafted mice after second-dose LNP-168-ABE8e-HBG injection. **c,** Quantitative assessment of β-like globin expression in patient cells from BM of engrafted mice after second-dose LNP-168-ABE8e-HBG injection. **d,** Patient cells engraftment in recipient BM measured by percentage of human CD45^+^cells (hCD45^+^) after LNP-168-ABE8e-HBG injection. **e,** Percentages of myeloid cells (hCD33^+^) and B cells (hCD19^+^) in the hCD45^+^ cell population in recipient BM after LNP-168-ABE8e-HBG injection using RT-qPCR. **f,** Percentages of human erythroid cells in recipient BM after LNP-168-ABE8e-HBG injection. **g,** Quantification of β-like globin expression *in vitro* differentiated erythroid cells from BM of engrafted mice after second-dose LNP-168-ABE8e-HBG injection using RT-qPCR. **h,** Assessment of enucleation in *in vitro* differentiated erythroid cells after LNP-168-ABE8e-HBG mediated base editing of patient cells. **i,** Cell size measured *in vitro* differentiated erythroid cells from BM of engrafted mice after second-dose LNP-168-ABE8e-HBG injection by relative forward scatter intensity. **j,** Circularity of *in vitro* differentiated enucleated erythroid cells from BM of engrafted mice after LNP-168-ABE8e-HBG injection by imaging flow cytometry. Each symbol represents a mouse, and there were a total of n=3 mice. All statistical significances in the figures were analyzed with the unpaired two-tailed Student’s *t*-test, and data represent mean ± SD, with *p*-values noted where appropriate. *NS*, not significant.

### Characterization of proteins constituting the protein corona in LNP-168-Luc

It was believed that following intravenous injection, LNPs swiftly develop a protein corona interface layer due to serum protein adhesion, fundamentally altering their surface properties and intricately determining them *in vivo* fate^18,51–53^. The controlled adsorption of specific plasma proteins on the LNPs surface may selectively direct LNPs to specific organs. Previous studies have identified apolipoprotein E (ApoE) as a crucial protein for LNPs, mediating their delivery to the liver by binding to the highly expressed low-density lipoprotein receptor (LDL-R) on hepatocytes^22,52,54^. Given the significantly BM-targeting abilities of LNP-168-mRNA, we postulate that the serum protein profile binding to LNP-168-mRNA would be distinctive, resulting in its increased targeting of BM cells. To examine this hypothesis, we exposed LNP-168-Luc to mouse plasma, isolated and retrieved protein-coated LNPs via centrifugation, and carried out a comprehensive PBS rinse (Extended Data Fig. 13A). The unbiased mass spectrometry proteomics technique allows us to identify and quantify proteins binding to LNP-168-Luc in plasma.

We detected 700 proteins adsorbed to LNP-168-Luc, with emphasis on those constituting over 0.1% of the total, anticipating their crucial roles in functions like immune response, coagulation, acute phase response, and lipid metabolism (Extended Data Fig. 13B and Supplementary Data Table 11,12). In contrast to liver-targeted LNPs, the lower levels of APOE adsorbed to LNP-168-Luc may result in reduced biological accumulation in the liver (Supplementary Data Table 13). The protein corona of BM-targeted LNP-168 is primarily composed of albumin, fibronectin, and fibrinogen, which rank as the top three proteins (Extended Data Fig. 13C). Albumin serves as the predominant protein in the corona of LNP-168-Luc, with studies highlighting the crucial role of serum proteins in the cellular uptake, protein expression, and endosomal release processes of LNPs^54^. Reports indicate that fibronectin, as an extracellular matrix protein in the BM, functions by binding to receptors on adjacent HSCs and stromal cells^55^. Additionally, studies suggest that fibrinogen coating can enhance endothelial cell adhesion and endothelialization^56,57^. Therefore, the potential mechanism for mRNA delivery, mediated by the protein corona on the surface of BM-targeted LNP-168-Luc, depends on the synergistic effects of specific proteins or multiple proteins, warranting further investigation^58^.

### Safety study following LNP-168-ABE8e treatment

To ensure the safety of utilizing LNP-ABE8e for clinical applications, we first conducted a systematic assessment of immune response or inflammation. Mice were intravenously injected with LNP-168 coencapsulated ABE8e mRNA and sgRNA-PCSK9 at 1.0 mg/kg, with a second dose after 14 days (Fig. 6a). To assess the pharmacokinetics of LNP-168-ABE8e-PCSK9, we quantified the amount of ABE8e mRNA and sgRNA-PCSK9 remaining in the plasma after injection. We found that both molecules peaked at 2 hours (h) post-administration, were rapidly cleared from the animals, and became undetectable after 72 h (Fig. 6b). Using plasma samples collected up to day 12, we measured the levels of Lipid-168 components and found that LNP-168-ABE8e-PCSK9 was cleared from the plasma with a half-life (T1/2) of approximately 2.5 h (Fig. 6c). In addition, we investigated systemic anti-Cas9 and anti-PEG specific antibody response through enzyme-linked immunosorbent assay (ELISA) using sera from the LNP-168-ABE8e-PCSK9 injected mouse. Compared with the mock group, anti-Cas9 and anti-PEG immunoglobulin G (IgG) was not evoked by second LNP-168-ABE8e-PCSK9 injection (Fig. 6d).

**Fig.6.**
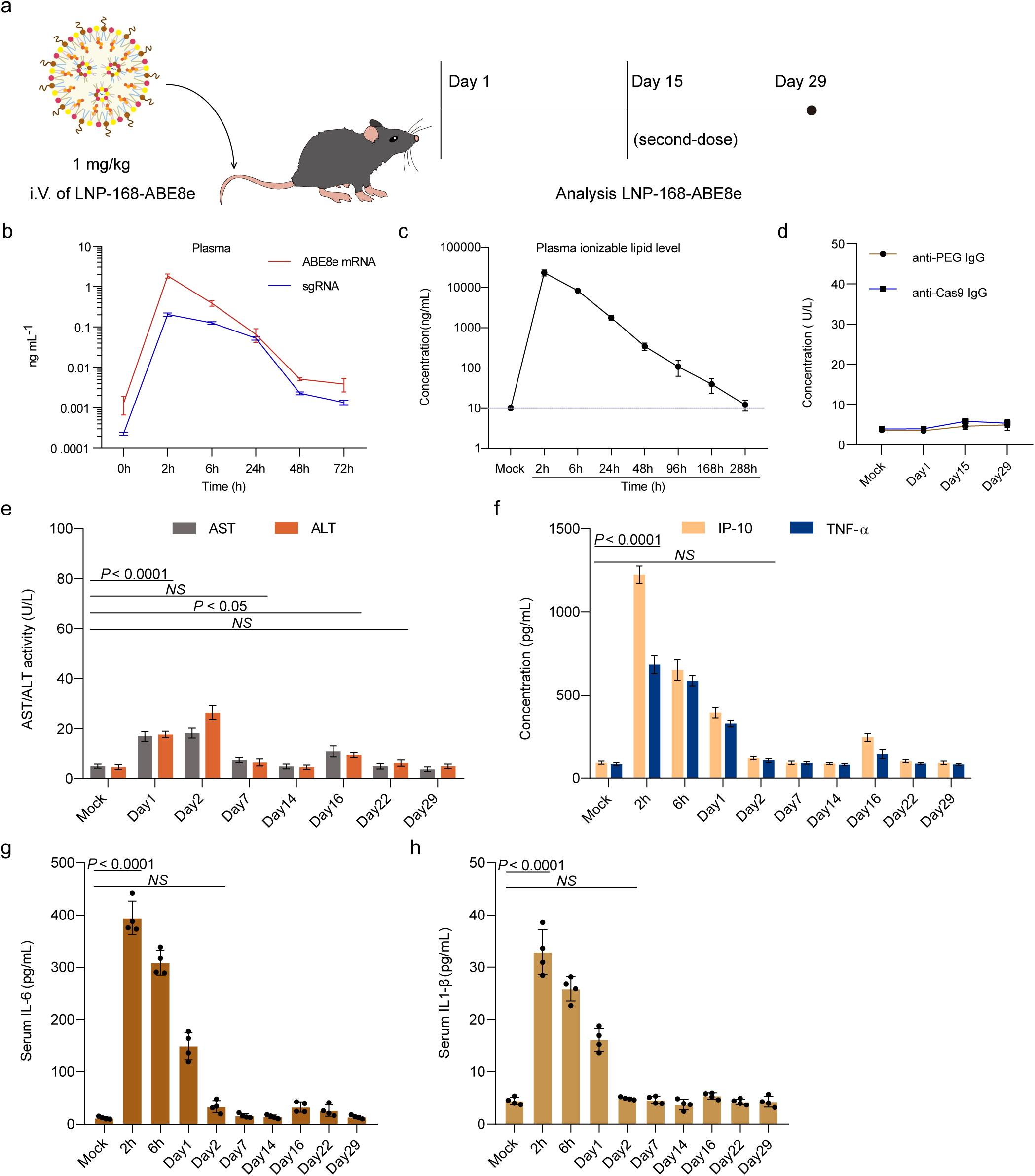
| Safety-related assessments of the LNP-168-ABE8e-PCSK9 treatment. **a,** C57 mice were administered LNP-168 encapsulated ABE8e and sgRNA-PCSK9, via tail vein injection at a dose of 1 mg/kg. Mice were initially injected once and then second-dosed with the same dose after 14 days, and were euthanized on day 29 for organ tissue collection. Collect serum or plasma at various time points for subsequent analysis. **b,** ABE8e mRNA and sgRNA levels in mouse plasma at different time points within 72 h after treatments with LNP-168-ABE8e-PCSK9. **c,** In a separate study, the clearance of Lipid-168 was measured in the plasma (dashed line=limit of quantification 10). **d,** Background-subtracted absorbance (A450-A540) representing the relative quantity of anti-Cas9 and anti-PEG IgG antibodies after LNP-168-ABE8e-PCSK9 injection. **e,** After administration of LNP-168-ABE8e-PCSK9 via tail vein injection as a single-dose and second-doses, serum was collected at various time points to assess the expression levels of ALT and AST, with the mock group used as a control. **f,** After administration of LNP-168-ABE8e-PCSK9 via tail vein injection as a single dose and second doses, serum was collected at various time points to assess the expression levels of IP-10and TNF-α, with the mock group used as a control. **g,** After administration of LNP-168-ABE8e-PCSK9 via tail vein injection as a single-dose and second doses, serum was collected at various time points to assess the expression levels of IL-6, with the mock group used as a control. **h,** After administration of LNP-168-ABE8e-PCSK9 via tail vein injection as a single-dose and second-doses, serum was collected at various time points to assess the expression levels of IL1-β, with the mock group used as a control. Each symbol represents a mouse, and there were a total of n=4 mice. All statistical significances in the figures were analyzed using one-way ANOVA with Dunnett’s multiple comparisons test, and data represent mean ± SD, with *p*-values noted where appropriate. *NS*, not significant.

Another substantial safety issue by LNP-168-ABE8e-PCSK9 is an immune response or inflammation. Following the administration of LNP-168-ABE8e-PCSK9, there were transient and moderate rises in AST and ALT that had entirely resolved by two weeks after LNP infusion, without inducing cumulative or lasting liver injury upon second dosing (Fig. 6e). Next, inflammatory response to LNP-168-ABE8e was evaluated by measuring serum immune-modulating chemokines (IP-10), tumor necrosis factor-α (TNF-α), IL-6 and interleukin-1β (IL-1β) concentrations. Although inflammatory cytokines were transiently increased after dosing, their elevated levels had resolved within 48 h, and no cumulative effects were observed with second dosing (Fig. 6f-g). Together, these data indicate that LNP-168-ABE8e gene editing strategy did not induce hepatic injury and immunogenic responses, thereby demonstrating a high level of safety.

Subsequently, our inquiry was extended to encompass an assessment of the DNA off-target impacts of ABE8e RNP edited cells *in vitro* and LNP-028-ABE8e-HBG edited cells *in vivo*, respectively. We employed next-generation sequencing (NGS) of the first 38 selected off-target sites analysis, focusing specifically on RNP-edited CD34^+^ cells (Supplementary Data Table 15)^59,60^. Among these sites, 38 exhibited nearly undetectable off-target base editing in edited cells compared to the mock group (Extended Data Fig. 14A). Furthermore, we extended our analysis to investigate off-target editing *in vivo* following LNP-028-ABE8e-HBG injection with efficient second-dose and fifth-dose administration in humanized *NCG-X* mice. Among these sites, we observed no off-target in edited cells compared to mock group at ten sites (Extended Data Fig. 14B). Consequently, these findings indicated that the LNP-028-ABE8e-HBG *in vivo* genome editing system exhibits minimal off-target editing effect. Furthermore, owing to the substantial homology between the duplicated HBG1 and HBG2 genes, concurrent editing of both HBG promoters using programmable nucleases that induce double-stranded breaks (DSBs) occasionally leads to a ∼4.9 kb deletion encompassing the HBG intergenic region, the consequences of which remain uncertain (Extended Data Fig. 14C)^26,61^. Thus, we assessed the frequency of these deletions via qRT-PCR in all edited samples. As anticipated, the 4.9 kb deletion targeted by sgRNA-25 was observed at a high frequency (62%) in Cas9-edited cells, with some deletions also detected in base-edited samples (9.1%, ABE8e), albeit significantly fewer than with Cas9 (Extended Data Fig. 14D). Additionally, we noted an increase in the frequency of large deletions with higher editing efficiency (Extended Data Fig. 14D-F). Based on these findings, we hypothesize that the simultaneous introduction of single-strand breaks (SSBs) by base editors at the editing sites of the HBG promoters could potentially lead to the generation of some large 4.9 kb deletions. Upon differentiation of cells into erythroid progenitors, the percentage of γ-globin induction was higher for ABE8e compared to Cas9 (Extended Data Fig. 14G), possibly attributable to a previously reported 4.9 kb deletion encompassing the HBG2 gene and the HBG1-HBG2 intergenic region. These findings indicate that ABE8e RNP exhibits high efficiency in editing the highly homologous regions of the γ-globin promoter, with minimal or no generation of large deletions, resulting in enhanced γ-globin induction compared to Cas9 RNP.

## DISCUSSION

Current HSCs gene therapy, involving isolating these cells from patients and complex *ex vivo* gene modifications using specialized equipment, is costly and technically challenging, limiting its accessibility. In contrast, pre-transplantation involves eliminating the patient’s stem cells through chemotherapy or radiation, causing significant toxicities^3^. Recognizing these challenges, use of LNPs to deliver mRNA encoding gene editing tools to HSCs *in vivo* would avoid the need for HSCs collection, *ex vivo* culture, and toxic patient conditioning^46^. Here, we have developed an *in vivo* gene editing technique using a non-antibody conjugated LNPs delivery system for efficient base editing through intravenous ABE8e/sgRNA mRNA injections, reactivating fetal hemoglobin expression in RBCs for cure of blood disorders.

In this study, we employed the ABE8e base editor to screen sgRNA sites associated with high HbF expression, which have confirmed therapeutic significance. Although previous studies suggest that the -123 T > C and -124 T > C substitutions may create a de novo *KLF1* binding site to upregulate HBG gene expression^30^, our data indicate that these mutations may also directly or indirectly affect *KLF1* binding through *SP1*. Further work is needed to evaluate these possibilities. We further encapsulated gene editors encoded by mRNA in LNP-028, achieving direct editing of BM cells in mice successfully. Utilizing this strategy, second injections of LNP-028 containing ABE8e mRNA and sgRNA-25 into humanized *NCG-X* mice yielded efficient base editing and significantly induced γ-globin expression in human cells. Secondary transplant of *in vivo* edited BM cells exhibited consistent base editing, γ-globin activation, and support reconstruction of multilineage human hematopoietic system, indicating the feasibility of generating efficient genome edits in LT HSCs by *in vivo* LNP-028-ABE8e-HBG delivery. Through further optimizing head group and tail structures based on library A, identifying LNP-168-ABE8e-PCSK9 exhibiting significantly enhanced editing efficiency (48.5% compared to 19.7% for LNP-028-ABE8e-PCSK9). Injection of LNP-168-ABE8e-HBG in a humanized β-thalassemia model resulted in 42.6% base editing at candidate targets and effective induction of γ-globin expression. We integrated miR-122 into the 3’-UTR of ABE8e mRNA, accomplishing efficient BM editing while reducing liver editing, providing a solution to improve targeting specificity. In a gene-edited mouse model, LNP-168 containing Cre recombinase mRNA could edit over 80.0% of MPP3 in all BM cells. Our research reveals that LNPs, carrying various mRNA cargoes, can effectively target HSPCs in the BM through a single systemic injection, with optimized ionizable lipid structures and miR-122 introduction enhancing delivery efficiency and specificity to HSPCs.

Furthermore, when using *in vivo* gene editing strategies to treat hemoglobinopathies, the selected targets for activating fetal hemoglobin, such as the BCL11A enhancer and the HBG1 promoter used in this study, only function in RBCs and not in liver cells. Therefore, even if editing occurs in liver cells during treatment, there is no safety risk, which is much safer compared to the toxicity generated by myeloablative conditioning used in HSCs transplantation.

Previous studies have demonstrated efficient genome editing in mice and non-human primates (NHPs) liver using LNPs, without inducing off-target mutations or significant safety concerns in animals^18–21^. We evaluated the clinical safety of LNP-168-ABE8e through hepatotoxicity, immunogenicity and off-target assessments of LNP-028-ABE8e. Mice treated with LNP-168-ABE8e-PCSK9 exhibited transient elevation in serum transaminases (ALT, AST) and some inflammation-related factors (IP-10, TNF-α, IL-6 and IL-1β), which normalized within 7 days. Second administration showed no cumulative effects, and no anti-Cas9 or anti-PEG antibodies were detected at 2- or 4-weeks post-treatment. The LNPs delivery system minimizes off-target editing risks by providing transient expression of genome editing agents, using biodegradable and non-toxic synthetic components, thus allowing potential repeated administration^12,20,49,62^. Differences in composition and preferentially enriched proteins between the protein corona formed on LNP-168-Luc and liver-targeted LNP-MC3 imply a crucial role for the protein corona in determining the *in vivo* tissue targeting of LNP-168-Luc, providing further design criteria for optimizing clinically relevant LNPs to enhance mRNA delivery^18,45,52,54^.

This study underscores the potential of LNPs delivery systems for genome editing in HSPCs as a promising and transformative treatment for SCD, β-thalassemia and other hematologic disorders, supported by robust evidence of safety and efficacy from our preclinical research. Besides applications for gene therapy, this LNP-ABE8e based editing system allows direct editing of BM cells including HSCs, immune cells, and other blood lineages, providing a fast and efficient way to explore gene functions without relying on genetically engineered animals.

In this study, we demonstrated that LNP-168 with ABE8e and sgRNA cargo can achieve efficient editing of HSCs without the need for coupled antibodies, which can significantly simplify the manufacturing process of LNPs drugs during clinical translation. Additionally, we used a mouse model transplanted with HSCs from TDT patients to prove that this LNP-168-ABE8e-HBG mediated *in vivo* gene editing strategy can effectively edit human HSCs and activate a sufficiently high level of fetal hemoglobin to restore the disease phenotype of patient RBCs. This reveals the tremendous potential application prospects of this therapeutic strategy in clinical settings.

## METHODS

### CD34^+^ HSPC cultures and *in vitro* differentiation

Human CD34^+^ HSPCs from mobilized peripheral blood of anonymized healthy donors and β-thalassemia patients were obtained from Xiangya Hospital Central South University and the First Affiliated Hospital of Guangxi Medical University. CD34^+^ HSPCs were enriched using the Miltenyi CD34 Microbead kit (Miltenyi Biotec). CD34^+^ HSPCs were thawed on day 0 in X-VIVO 15 (Lonza, 04-418Q) supplemented with 100 ng ml^−1^ stem cell factor (PeproTech, 300-07), 100 ng ml^−1^ human thrombopoietin (PeproTech, 300-18), and 100 ng ml^−1^ recombinant human Flt3-ligand (PeproTech, 300-19). HSPCs were electroporated with ABE8e protein 24 h after thawing and maintained in X-VIVO media with cytokines. For *in vitro* erythroid maturation experiments, 24 h after electroporation, HSPCs were transferred into erythroid differentiation medium (EDM) consisting of IMDM (Corning, 15-016-CV) supplemented with 330 μg ml^−1^ holo-human transferrin (Sigma-Aldrich, T4132), 10 μg ml^−1^ recombinant human insulin (Sigma-Aldrich, T4132), 2 IU ml^−1^ heparin (Sigma-Aldrich, H3149), 5% human solvent detergent pooled plasma AB (Gemini, 100-318), 3 IU ml^−1^ erythropoietin (Amgen, 55513-144-10), 1% L-glutamine (Life Technologies, 25030-164), and 1% penicillin/streptomycin (Gibco, 15140122). During days 0-7 of culture, EDM was further supplemented with 10^−6^ M hydrocortisone (Sigma, H0135), 100 ng ml^−1^ human SCF, and 5 ng ml^−1^ human IL-3 (PeproTech, 200-03) as EDM-1. During days 7-11 of culture, EDM was supplemented with 100 ng ml^−1^ human SCF only as EDM-2. During days 11-18 of culture, EDM had no additional supplements (EDM-3). Enucleation frequency and γ-globin induction were assessed on days 18 of erythroid culture.

### HUDEP cell culture

HUDEP-2 cells were grown in culture as described previously^25^. HUDEP-2 cells were maintained in an expansion medium: StemSpan SFEM (STEMCELL Technologies, 09650) with 50 ng ml^−1^ human stem cell factor (PeproTech, 300-07), 3 IU ml^−1^ erythropoietin, 10^-6^ M dexamethasone (Sigma-Aldrich, D4902), and 1 µg ml^−1^ doxycycline (Sigma-Aldrich, D9891) and passaged every three days. Erythroid differentiation was carried out by replacement the medium with EDM2 (IMDM) supplemented with 330 μg ml^−1^ holo-human transferrin, 10 μg ml^−1^ recombinant human insulin, 2 IU ml^−1^ heparin, 5% inactivated plasma, 3 IU ml^−1^ erythropoietin, 2 mM L-glutamine, 100 ng ml^−1^ SCF, 1 μg ml^−1^ doxycycline.

### Design and synthesis of lentiviral sgRNA libraries

We identified every 20-mer sequence upstream of the SpCas9 *NGG* PAM sequence at the HBG locus from hg38. We designed 92 sgRNAs sequences (including 9 negative control genome-targeting sgRNAs with 4 non-targeting and five AAVS1 targeting). Sequences information of sgRNAs are listed in Supplementary Data Table 2. sgRNAs library was constructed according to a published protocol^63^. The sgRNAs oligos (synthesized by Agilent) were cloned using Gibson Assembly master mix (New England Biolabs) into lentiGuide-Puro (addgene plasmid no. #52963) that had been digested with BsmBI, gel purified, and dephosphorylated. The assembly products were transformed into NEB-stable competent cells, and then the plasmids were extracted. Sufficient colonies were isolated to ensure > 1,000 colonies per spacer sequence. Plasmid libraries were deep-sequenced to confirm representation.

To produce lentivirus, HEK293T cells were cultured with DMEM (Gibco, C11995500CP) supplemented with 10% fetal bovine serum (Gibco, 10270-106) and 2% penicillin/streptomycin (Gibco, 15140122) in 15 cm tissue culture-treated Petri dishes. HEK293T cells were transfected at 80% confluence in 12 mL of medium with 13.3 μg of psPAX2 (addgene plasmid no.#12260), 6.7 μg of VSV-G (addgene plasmid no. #14888) and 20 μg of the lentiviral construct plasmid of interest using 180 μg of linear polyethylenimine (Polysciences). The medium was changed 8-12 h after transfection. Lentiviral supernatant was collected at 48 and 72 h post-transfection.

### Transduction of HUDEP-2 cells with a lentiviral library and analysis

We procured lentiCas9-Blast (addgene plasmid no.#52962) and ABE8e (addgene plasmid no.#138489) vectors from the addgene website. To construct the novel lentiABE8e-nCas9-Blast plasmid, distinct DNA fragments were amplified using different DNA polymerases (TOYOBO, KOD-Plus-Neo, KOD-401), and a complete plasmid was successfully assembled using the Gibson Assembly method (NEB, E2611L). To achieve inactivation of the nCas9 (D10A) sequence, a mutation was introduced at the 840th amino acid position, replacing the histidine residue with alanine, thereby converting it into dCas9 (D10A, H840A). This process resulted in the generation of the new lentiABE8e-dCas9-Blast plasmid, followed by lentivirus packaging with the protocols described above.

HUDEP-2-ABE8e cells were transduced with the library at a low multiplicity of infection of 0.3 to minimize the transduction of cells with more than one vector particle and to achieve an approximately 1,000-fold coverage of the sgRNAs library. After 24 h, transduced cells were selected in puromycin (1 µg ml^−1^) for two additional days and then maintained in the expansion medium for 5 days. Erythroid maturation was induced in differentiation medium for 5 days. Cells were then stained for HbF as indicated above and sorted into HbF-high and HbF-low populations. Genomic DNA was extracted with the PureLink Genomic DNA Mini Kit (Thermo Fisher Scientific, K182001). sgRNAs were amplified for 27 cycles using barcoded lentiGuide-Puro-specific primers with 400 ng genomic DNA per reaction. The PCR products were then pooled for each sample. The barcoded libraries were pooled at an equimolar ratio, gel purified and subjected to massively parallel sequencing with a MiSeq instrument (Illumina), using 75-bp paired-end sequencing.

For data analysis, to identify significantly enriched sgRNAs in the top 10% of between HbF-high and HbF-low cells, we employed the robust rank aggregation (RRA) method of MAGeCK with default parameters. Candidate upregulated sgRNAs were determined using filters of fold change (*FDR*) < 0.05 and log2 *FDR* > log2 (1.5). This ranking reflects the relative HbF enrichment value of the sgRNAs, with rankings surpassing a specific threshold indicating a pronounced effect on activating HbF expression. Our experimental design incorporated biological replicates, which yielded highly reproducible results (*R > 0.98* and *R > 0.94*) for both nCas9 and dCas9 screens, demonstrating a strong correlation between the two screening methods. To enhance the accuracy of our findings, we combined the two screening methods of nCas9 and dCas9, identifying same sgRNAs present in the intersection. Out of the 45 top-ranked sgRNAs selected from the two libraries, we filtered out those that did not appear in the intersection, resulting in a final selection of 40 sgRNAs (Supplementary Data Table 3).

### Flow cytometry analysis

Intracellular staining was performed as described previously^29^. Cells were fixed with 0.05% glutaraldehyde (Sigma, 111-30-8) for 10 min at room temperature and then permeabilized with 0.1% Triton X-100 (Thermo Fisher Scientific, HFH10) for 5 min at room temperature. Cells were stained with anti-human antibodies for HbF (Human Fetal Hemoglobin FITC Conjugate; Life Technologies, MHFH01) for 30 min in the dark. Cells were washed to remove unbound antibodies before FACS analysis was performed. Control cells without staining were used as negative control. For erythroid cells analysis, cells were also stained with FITC anti-human CD235a Antibody (BioLegend, 349104) and with PE anti-human CD71 Antibody (BioLegend, 334106) to determine the maturation of erythroid progenitor cells. For enucleation analysis, cells were stained with 2 μg ml^−1^ of the cell-permeable DNA dye Hoechst 33342 (Thermo Fisher Scientific, 62249) for 10 min at 37°C. The Hoechst 33342-negative cells were further gated for cell size analysis with Forward Scatter (FSC) A parameter. Median value of forward scatter intensity normalized by data from healthy donors collected at the same time was used to characterize the cell size.

### Imaging flow cytometry analysis

*In vitro* differentiated day18 erythroid cells stained with Hoechst 33342 were resuspended with 150 μL DPBS for analysis with Imagestream X Mark II (Merck Millipore). Well-focused Hoechst-negative single cells were gated for circularity analysis with IDEAS software. Cells with circularity score above 15 were further gated to exclude cell debris and aggregates. No fewer than 2000 gated cells were analyzed to obtain a median circularity score.

### RNA-seq and analysis

For RNA-seq, RNA was extracted from 1 million HUDEP-2 cells (at least three biological replicates each) using TRIzol reagent (Thermo Fisher Scientific,15596026) following the manufacturer’s procedure. Poly (a) RNA was purified from 1 μg total RNA using Dynabeads Oligo (dT) 25-61005 (Thermo Fisher Scientific, 61005) using two rounds of purification. Then, the poly (a) RNA was fragmented into small pieces using a Magnesium RNA Fragmentation Module (NEB, e6150) at 94℃ for 5-7 min. Cleaved RNA fragments were reverse transcribed to create cDNA by SuperScript II Reverse Transcriptase (Invitrogen, 1896649), which were then used to synthesize U-labeled second-stranded DNAs with E. coli DNA polymerase I (NEB, m0209), RNase H (NEB, m0297) and dUTP Solution (Thermo Fisher Scientific, R0133). An A-base is then added to the blunt ends of each strand, preparing them for ligation to the indexed adapters. Each adapter contains a T-base overhang for ligating the adapter to the A-tailed fragmented DNA. Single- or dual-index adapters were ligated to the fragments, and size selection was performed with AMPureXP beads. The average insert size for the final cDNA library was 300 ± 50 bp. Finally, 150 bp paired-end sequencing (PE150) was performed on the Illumina NovaSeq 6000 platform (LC-Bio Technology, Hangzhou, China) following the vendor’s recommended protocol. After removing the low-quality and undetermined bases, we used HISAT2 software (https://daehwankimlab.github.io/hisat2/) to map reads to the genome (Homo sapiens Ensemble v96). The differentially expressed mRNAs were selected with *FDR* > 2.0 or *FDR* < 0.5 and *p* value < 0.05 by DESeq2^64^.

### Base editor protein expression and purification

For ABE8e protein expression, were expressed in E. coli BL21 (DE3) (Thermo Fisher Scientific, EC0114), which were grown in LB media at 37°C and 4-5 h later induced by 1 mM isopropyl β-d-1-thiogalactopyranoside (IPTG) (Sigma-Aldrich, 367-93-1) for 18-20 h at 16°C. Cell paste was lysed in 20 mM Tris, 10 mM imidazole, 500 mM NaCl, and 1.5 M urea pH 8 by ultrasonic crushing. The lysate was centrifuged at 10000 × g for 45 min. Proteins were purified using nickel-NTA resin, G25 desalting column desalting, and cation exchange chromatography sequentially. Endotoxin was controlled by anion exchange chromatography, and the flow through from Q-HP column was concentrated in Ultra-15 centrifugal filters (Ultracel-30K, Merck Millipore UFC903024) to concentration of ∼10 mg ml^-1^ (UV280). The purified protein was concentrated in 30 mM HEPES buffer of pH 7.4 containing 150 mM NaCl and 10% glycerol. Protein samples and fractions were separated using SDS-PAGE and stained using Gel Code Blue Stain Reagent (Thermo Fisher Scientific, 24590).

### RNP electroporation

Electroporation was performed using a Lonza 4D Nucleofector (V4XP-3032 for 20 μL Nucleocuvette Strips or V4XP-3024 for 100 μL Nucleocuvette Strips) according to the manufacturer’s instructions. Modified synthetic sgRNAs (2’-O-methyl-3’phosphorothioate linkage modifications in the first and last three nucleotides) were purchased from Genscript. sgRNA concentration was calculated using the full-length product reporting method, which is threefold lower than the OD reporting method. CD34^+^ HSPCs were thawed 24 h before electroporation. For 20 μL Nucleocuvette Strips. RNP complexes were prepared by mixing ABE8e protein (200 pmol) and sgRNA (600 pmol) and incubating for 15 min at room temperature immediately before electroporation. 50000 HSPCs re-suspended in 20 μL of P3 solution were mixed with RNP and transferred to a cuvette for electroporation with the EO-100 program. For 100-μL cuvette electroporation, the RNP complex was made by mixing 1000 pmol ABE8e protein and 3000 pmol sgRNA. Next, 5 M HSPCs were re-suspended in 100 μL of P3 solution for RNP electroporation as described above. The electroporated cells were resuspended in X-VIVO media with cytokines and changed to EDM 24 h later for *in vitro* differentiation. For mouse transplantation experiments, cells were maintained in X-VIVO 15 with SCF, TPO, and Flt3-L for 0-2 days as indicated before infusion.

### Measurement of base editing

Editing frequencies were measured in EDM cultured cells 5 days after electroporation. Briefly, genomic DNA was extracted using the DNeasy Blood & Tissue Kit (Qiagen, 69504). Using *Benchling* (https://www.benchling.com) and *CasOFFinder* tool^60^, 38 potential off-target sites with four or fewer genomic mismatches were identified. The HBG promoter region or off-target sites were amplified with KOD Hot Start DNA Polymerase and corresponding primers (Supplementary Data Table 5,12). The resulting PCR products underwent Sanger sequencing or Illumina deep sequencing. For Sanger sequencing, traces were imported into *EditR* software for base editing measurement^65^. Frequencies of editing outcomes were quantified using *CRISPResso2* software (version 2.0.31) and collapsed on the basis of mutations in the quantification window^66^. Equal amounts of the PCR product were sent for NGS (https://doi.org/10.1007/s11427-018-9402-9) at the State Key Laboratory of Rice Biology using the *Hi-TOM* platform (China National Rice Research Institute, Chinese Academy of Agricultural Sciences, Hangzhou).

### RT-qPCR quantification of γ-globin induction

RNA isolation with RNeasy columns (Qiagen, 74106), reverse transcription with Primescript RT Master Mix (TAKARA, RR036A-1), and RT-qPCR with universal SYBR green master (Roche, 04913850001) were performed to determine γ-globin induction using primers amplifying HBG1/2, *HBB*, or *HBA1/2* cDNA (Supplementary Data Table 6).

### qPCR detection of 5.2-kb deletion alleles

A primer and probe set were designed to detect the amplification of an HBG1 promoter-specific sequence (5.2-kb-Forward: ACGGATAAGTAGATATTGAGGTAAGC; 5.2-kb-Reverse: GTCTCTTTCAGTTAGCAGTGG; TaqMan probe (FAM): ACTGCGCTGAAACTGTGGTCTTTATGA). TaqMan qPCR was performed on genomic DNA samples from CD34^+^ cells using Universal TaqMan Mix (Thermo Fisher Scientific, 4304437) for the quantification of triplicates for each sample. ΔΔCt values were calculated based on the amplification of the sequence encoding TERT (Thermo Fisher Scientific, 4403316) for copy number reference.

### Hemoglobin HPLC

Hemolysates were prepared from either erythroid cell after 18 days of differentiation for *in vitro* differentiation experiments or human erythroid cells directly purified from xenotransplanted mouse BM by CD235a microbeads (130-050-501, Miltenyi Biotec). For single-chain globin variants, 2 × 10^6^ erythroid cells were lysed using 20 μL of ddH_2_O. Reverse HPLC was performed on an Agilent 1260 Infinity II using a 4.6-nm Aeris 3.6 mM Widepore C4 LC column.

### Human CD34^+^ HSPC transplantation and flow cytometry analysis

All animal experiments were approved by ECNU Animal Care. The protocol (protocol ID: m20210238) was approved by the ECNU Animal Care and Use Committee. NOD-*Prkdc^em26Cd52^Il2rg^em26Cd22^kit^em1Cin^*(V831M)/Gpt (NCG-X) mice were obtained from GemPharmatech (Shanghai, China). No irradiated NCG-X female mice (4-5 weeks of age) were infused by retro-orbital injection with 0.8 million CD34^+^ HSPCs (live cells counted immediately before infusion, re-suspended in 200 μL of DPBS) derived from healthy donors. BM was isolated for human xenograft analysis after 16 weeks of transplantation. In the *in vivo* experiments involving NCG-X mice, LNP-ABE8e-HBG was administered via the lateral tail vein to deliver to the BM cells of the mice, 16 weeks post CD34^+^ HSPCs transplantation. Each animal received a fixed injection volume of 0.2 mL. The mobilization regimen of 4 days of G-CSF (filgrastim [Neupogen]; Amgen Thousand Oaks, CA) injection followed by an injection of plerixafor (Mozobil®, Sanofi, Cambridge, MA, USA) on day 5 is widely used for robustly mobilizing HSCs from the BM.

For flow cytometry analysis of BM cells, they were first incubated with V450 Mouse Anti-Human CD45 Clone HI30 (BD Biosciences, 560367), PE-eFluor 610 mCD45 Monoclonal Antibody (Thermo Fisher Scientific, 61-0451-82), FITC anti-human CD235a Antibody (BioLegend, 349104), PE anti-human CD33 Antibody (BioLegend, 366608), FITC anti-human CD34 Antibody (BioLegend, 343504), APC anti-human CD19 Antibody (BioLegend, 302212), and Fixable Viability Dye eFluor 780 for live/dead staining (Thermo Fisher Scientific, 65-0865-14). The percentage of human engraftment was calculated as hCD45^+^ cells/ (hCD45^+^ cells + mCD45^+^ cells) × 100. B-cell (CD19^+^) and myeloid (CD33^+^) lineages were gated on the hCD45^+^ population. Human erythroid cells (CD235a^+^) were gated on the mCD45^-^hCD45^-^ population. After washing with PBS, flow cytometric analysis or cell sorting was performed. Cell sorting was performed on a FACSAria II machine (BD Biosciences).

### ATAC-seq

ATAC-seq was performed as previously described^67^. A total of 50000 cells of differentiation phase CD34^+^ cells were collected and permeabilized with 50 mL ice cold lysis buffer (10 mM Tris-HCl pH 7.4, 10 mM NaCl, 3 mM MgCl_2_, 0.1% IGEPAL CA-360) for 10 minutes. The unfixed nuclei of these cells were tagged using tn5 transposase (TruePrep DNA Library Prep Kit V2 for Illumina) for 30 min at 37°C, and the resulting library fragments were generated by 10-12 PCR cycles. PCR amplification was carried out as follows: 72℃ for 3 min, 98℃ for 30 s, 98℃ for 15 s, 60℃ for 30 s, 72℃ for 30 s, and 72℃ for 5 min. Steps 3-5 were repeated for another 13 cycles and held at 4℃. The resulting libraries were purified using a QIAGEN MinElute PCR purification kit, quantified with a Qubit fluorometer and bioanalyzer, and sequenced on an Illumina HiSeq Xten-PE150.

ATAC-seq data processing was performed as previously described^68^, with slight modifications. Briefly, Trimmomatic was used to filter raw NGS data with default parameters, trimming adaptor sequences, and low-quality bases. Clean reads were aligned to the human genome (hg38) using Bowtie2 with settings--very-sensitive-k 10-X 2000. Then, low mapping quality and mitochondrial reads were removed with SAMtools and custom script. To call peaks, the Genrich package was used for all test samples, while the Blacklist regions bed file for humans (hg38) was downloaded from Github. Finally, for visualization of specific regions, compute Matrix, plotHeatmap, and plotProfile functions from deepTools were performed on bigwig format data generated by bamCoverage from shifted aligned reads.

### CUT&Tag

The CUT&Tag assay was performed as described previously with modifications^69^. A total of 1.0[×[10^5^ CD34^+^ cells were washed with 1 mL of wash buffer (20 mM HEPES pH 7.5), 150 mM NaCl, 0.5 mM spermidine (Sigma-Aldrich, 85558), 1.0 × protease inhibitor cocktail (Roche) and centrifuged at 1600[r.p.m. for 5[min at room temperature. Cell pellets were resuspended in 1 mL of wash buffer. Concanavalin A-coated magnetic beads were washed twice with binding buffer (20 mM HEPES pH 7.5, 10 mM KCl, 1 mM MnCl_2_, 1 mM CaCl_2_). Next, 10 μL of activated beads was added and incubated at room temperature for 15 min. Bead-bound cells were resuspended in 50 μL of antibody buffer (20 mM HEPES pH 7.5, 150 mM NaCl, 0.5 mM spermidine, 0.05% digitonin (Sigma-Aldrich, 11024-24-1), 2 mM EDTA, 0.1% BSA, 1.0 × protease inhibitor cocktail. Then, 1[μg of primary antibody rabbit monoclonal anti-SP1 antibody (Abcam, EPR22648-50) and goat anti-rabbit IgG H&L (Abcam, ab6702) was added and incubated overnight at 4°C with slow rotation. The primary antibody was removed using a magnet stand. A secondary antibody (1 μg) was diluted in 50 μL of dig-wash buffer (20 mM HEPES pH 7.5, 150 mM NaCl, 0.5 mM spermidine, 0.05% digitonin, 1.0 × protease inhibitor cocktail), and the cells were incubated at room temperature for 1 hour. Cells were washed three times with dig-wash buffer to remove unbound antibodies. The Hyperactive pA-Tn5 Transposase adapter complex (Vazyme, TTE mix, 4 μM) was diluted 1:100 in 100 μL of Dig-300 buffer (20[mM HEPES pH 7.5, 300 mM NaCl, 0.5 mM spermidine, 0.01% digitonin, 1.0 × protease inhibitor cocktail). Cells were incubated with 0.04 μM TTE mix at room temperature for 1 hour. Cells were washed three times with Dig-300 buffer to remove unbound TTE mix. Cells were then resuspended in 300 μL of tagmentation buffer (10 mM MgCl_2_ in Dig-300 buffer) and incubated at 37°C for 1 hour. To terminate tagmentation, 10 μL of 0.5 M EDTA, 3 μL of 10% SDS and 5 μL of 10[mg[ml^−1^ proteinase K were added to 300 μL of sample and incubated at 50°C for 1[hour. DNA was purified using phenol-chloroform-isoamyl alcohol extraction and ethanol precipitation as well as RNase A treatment.

For library amplification, 24 μL of DNA was mixed with 1 μL of TruePrep Amplify Enzyme (Vazyme, TAE), 10 μL of 5 × TruePrep Amplify Enzyme buffer, and 5 μL of ddH_2_O, as well as 5 μL of uniquely barcoded i5 and i7 primers from TruePrep Index Kit V2 for Illumina (Vazyme, TD202). A total volume of 50 μL of the sample was placed in a Thermocycler using the following program: 72°C for 3 min; 98°C for 30 s; 17 cycles of 98°C for 15 s, 60°C for 30 s and 72°C for 30s; 72°C for 5[min and hold at 4°C. To purify the PCR products, 1.2 × volumes of VAHTS DNA Clean Beads (Vazyme, N411-02) were added and incubated at room temperature for 10[min. Libraries were washed twice with 80% ethanol and eluted in 22 μL of ddH_2_O. Libraries were sequenced on an Illumina NovaSeq platform, and 150 bp paired-end reads were generated.

CUT&Tag data processing was performed as previously described with slight modifications^69^. Briefly, trimmomatic was used to filter raw NGS data with default parameters, trimming adaptor sequences and low-quality bases^70^. Clean reads were aligned to the human genome (hg38) using Bowtie2 with settings--local--very-sensitive--no-mixed--no-discordant--phred33-I10-X 1000. Then, low mapping quality and unmapped reads were removed with SAMtools^71^. Format conversion to bedgraph and normalization were performed using the bedtools genomecov function (v2.30.0). To call peaks, the SEACR package was used for test samples against the wild type with default parameters. For visualization of specific regions, computeMatrix, plotHeatmap, and plotProfile functions from deepTools were performed on bigwig format data generated by bamCoverage^64^.

### RNA synthesis

An mRNA encoding ABE8e protein was synthesized *in vitro* using a T7 RNA polymerase-mediated transcription with complete replacement of uridine by N1-methylpseudouridine^72^. The DNA template used in the *in vitro* reaction contained the ABE8e open-reading frame flanked by 5’-UTR and 3’-UTR sequences and was terminated by a polyA tail. Capping of the *in vitro* transcribed mRNAs was performed co-transcriptionally using the trinucleotide cap1 analog, CleanCap (TriLink, N-7413). mRNA was purified by RNeasy Mini Kit (QIAGEN, 74104) purification as described. All mRNAs were analyzed by an automatic nucleic acid analysis system (Bioptic, Qsep-100) and were stored frozen at -80°C.

### Preparation of LNPs

LNPs used in this study are proprietary to Yoltech Therapeutics (Shanghai, China). The ionizable cationic lipid and LNPs composition are described in CN114989182A, CN116162071A, PCT/CN2023/100791, PCT/CN2023/100823 and PCT/CN2023/106421. Ionizable lipids (DLin[MC3[DMA, MC3) were purchased from Cayman Chemicals. DSPC, DMG-PEG2000 were purchased from Avanti Polar Lipids. Cholesterol was purchased from Sigma-Aldrich. LNPs were formulated with an amine-to-RNA-phosphate (N:P) ratio of 4.5. The lipid nanoparticle components were dissolved in 100% ethanol with the following molar ratios: 50 mol% ionizable lipid, 38.5 mol% cholesterol, 10 mol% DSPC, and 1.5 mol% DMG-PEG2000. The ABE8e mRNA and sgRNA cargo in a 1:1 ratio was encapsulated in a LNPs through an ethanol-drop nanoprecipitation process^16,17^. Briefly, the LNPs were rapidly mixed using Precision NanoSystems NanoAssemblr Benchtop Instrument with mRNA in citrate buffer of pH 5. The RNA-loaded dispersions were neutralized by buffer exchanged into phosphate-buffered saline (PBS), pH 7.4, and filtered using a 0.2 µm sterile filter. Formulated LNPs mRNA was then vialed and stored refrigerated at 4°C until further use. Encapsulation efficiencies were determined by Quant-iT Ribogreen assay (Thermo Fisher Scientific, R11490). Particle size and polydispersity were measured by dynamic light scattering using a Malvern Zetasizer DLS instrument. All ionizable lipids were synthesized using standard organic procedures (supporting information). The diameters of all LNPs encapsulating mRNA ranged between 69 to 90 nm, displaying a uniform nanosize distribution (Supplementary Data Table 7, 8 and 9). The encapsulation efficiency of all LNPs mRNA formulations was assessed to be > 90% (Supplementary Data Table 7, 8 and 9).

### TNS assay

pKa of each LNPs was determined by 2-(p-toluidino)- 6-napthalene sulfonic acid (TNS) assay^73^. Briefly, pH buffers from 2.5 to 11.0 in 0.5 increments were prepared by adjusting a solution of 20 mM sodium phosphate, 25 mM citrate, 20 mM ammonium acetate, and 150 mM NaCl with 0.1 N NaOH and 0.1 N HCl. Each pH buffer was added to a black 96-well plate in 100 μL. In addition, 300 μM stock solution of TNS was added to the above pH buffer in the final concentration of 6 μM. Last, formulated LNPs were added to the mixture solution in the final concentration of 25 μM. Fluorescence intensity was obtained by Tecan at an excitation of 325 nm and an emission of 435 nm. The measured fluorescence was normalized by (fluorescence−minimum fluorescence) divided by the (maximum fluorescence−minimum fluorescence). The normalized fluorescence was evaluated by a curve fit analysis, resulting in a fluorescence titration S curve. pKa of each LNPs was calculated as the pH value where half of the maximum fluorescence is reached.

### Cryogenic Transmission Electron Microscopy of LNPs

A volume of 3 μL of the sample was placed on a Cu grid (300 mesh, 1.2/1.3, Quantifoil), which was glow-discharged using a PELCO easiglow under the conditions of 15 mA for 100 s. The grid was then blotted with filter paper to remove excess liquid, forming a thin film (blot time: 4 s, blot force: 0, wait time: 30 s). The grid was rapidly vitrified in liquid ethane at -180°C to prevent the crystallization of water. The frozen grids were assembled into autogrids to form complete cartridges and were maintained at temperatures below -180°C during the experiment. Cryo-EM observation was performed using a Glacios instrument from ThermoFisher Scientific, operated at 200 kV. Images were recorded with a Ceta-D camera from ThermoFisher Scientific.

*In vitro* transfection of BM cells with LNPs.

BM cells were derived from the femurs of animals, and their recovery was optimized through mechanical crushing after the removal of muscle and connective tissues. The isolated BM cells were suspended in a solution containing 4% FBS (Gibco, 10270-106) and PBS (Gibco, 14287080). RBCs were lysed at room temperature for 10 minutes using ACK lysis buffer (Gibco, A1049201). The cell suspension was subsequently filtered through a 40 μm sterile strainer (Corning, 431750) and rinsed with a 4% FBS PBS solution. After cell resuspension, the cells were enumerated and viability was evaluated using AOPI staining (Nexcelom Bioscience, CS2-0106) on a Countstar Altair cell viability counter. Subsequently, cells were seeded at a concentration of 1.0E + 06 mL^-1^ in Stemspan SFEM (STEMCELL Technologies, 096000) obtained from Stemcell Technologies. The culture medium was supplemented with 50 ng mL^-1^ mSCF (PeproTech, 250-03), 6 ng mL^-1^ mIL-3 (PeproTech, 213-13), and 10 ng mL^-1^ mIL-6 (PeproTech, 216-16), all provided by the same company. LNPs mRNA formulation was added 6 h after cell plating, and after 24 h of transfection, the medium was replaced. Cells were collected 72 h post-transfection, and the editing efficiency was assessed.

### Bioluminescence imaging

The C57/B6J mice (6-8 weeks) were purchased from GemPharmatech (Shanghai, China). C57BL/6J mice were intravenously injected with formulations of LNP-168-Luc, LNP-168-Luc-miR-122T (Supplementary Data Table 9,10). For *in vivo* imaging, each mouse was intraperitoneally injected with luciferin substrate solution 24 h after intravenous injection (150 mg/kg; Luc001, Shanghai Sciencelight Biology Science & Technology). After a 10-minute interval, the mice were euthanized, and the desired tissues were harvested, washed with PBS, and promptly positioned on the imaging platform. The femurs were gently crushed using a spatula to expose the bone marrow for imaging. The bioluminescence images of the mice were acquired using an IVIS Lumina II *in vivo* imaging system (PerkinElmer) with an exposure time of 60 s or longer to ensure the captured signal fell within the optimal detection range. Subsequently, the bioluminescence images were analyzed using Living Image (version 4.3.1).

### Delivery of Cre mRNA by LNPs

B6.Cg-Gt(ROSA)26Sortm14(CAG-tdTomato)Hze/J Ai14 mice (purchased from Jackson Labs, #007914) were bred in house and both males and females were used for experiments. To explore the effect of LNP-168-Cre-miR-122T injection on *in vivo* transfection of BM HSCs and other cell lineages, we administered intravenous injections to Ai14 mice at a dose of 1 mg/kg (Supplementary Data Table 8). The expression of the reporter protein in cells isolated from BM was detected through flow cytometry (Cre-induced tdTomato fluorescence) and compared to mice treated with LNP-MC3-Cre-miR-122T. Four days after a single injection, we monitored the induction of the reporter gene expression (tdTomato^+^) in LSK, HSCs and other cell lineages. For BM isolation, femurs and tibias were collected, washed in a 4% FBS PBS solution, filtered through a 40 μm cell strainer, and then centrifuged (400 g, 10 min). The obtained cell pellets were lysed with ACK buffer for 10 minutes to lyse RBCs, diluted with a 4% FBS PBS solution, filtered, centrifuged, resuspended in a 4% FBS PBS solution, and counted.

For flow cytometry staining, cell suspensions were first washed, then blocked with BSA for 10min on ice prior to adding fluorescent antibodies for cell surface staining. For mouse BM, cells were stained with lineage markers such as CD11b (Biolegend, 101216)、CD3 (Biolegend, 100206)、Gr1 (Biolegend, 108424)、B220 (Biolegend, 103212) and TER-119 (Biolegend, 133307), as well as stem cell markers CD117/c-Kit (Biolegend, 105826), Sca-1 (eBioscience, 12-5981-82), CD150 (Biolegend, 115914), CD48 (Biolegend, 103421), and CD135 (eBioscience, 17-1351-82). Lineage negative single cells were gated in a bivariate plot of c-Kit and Sca-1 to identify double positive LSK cells. Mice HSPC subsets were defined as follows: HSC Long-term Hematopoietic stem cell (Lin^-^LSK^+^CD135^-^CD48^-^CD150^+), MPP multipotent progenitor (Gate^^1, Lin-^LSK^+^CD135^+^CD150^-; Gate^^2, Lin-LSK+^CD135^-^CD48^+^CD150^-; Gate^^3, Lin-^LSK^+^CD135^-^CD48^+^CD150^+; Gate^^4^, Lin^-^LSK^+^CD135^-^CD48^-^CD150^-^)^50,74^. Single color and fluorescence overlayed images were generated to depict the cells containing tdTomato fluorescence. For flow cytometry acquisition, cells were run on a BD flow cytometer (Fortessa or Fusion) and analyzed using FlowJo software (BD).

### Toxicity and cytokine assays

In all mouse studies, serum samples were routinely collected at the following time points: Mock (un-treated), 2 h (Day0, after LNP-168-ABE8e-PCSK9 infusion), 6 h (Day0), 24 h (Day1), 48 h (Day2), Day 7, Day14, Day16 (24 h, after second-dose LNP infusion), Day22, and Day29. To assess hepatic toxicity, serum samples were collected at various time points over a 4-week period and allowed to stand at 4°C until coagulation was complete. Serum was centrifuged at 10000 rpm for 10 minutes at 4°C, and the supernatant was collected. Subsequent experiments were conducted in accordance with the manufacturer’s instructions. The activities of serum ALT and AST were determined using commercially available assay kits (Nanjing Jiancheng, C010-2-1, C009-2-1). Additionally, we quantitatively measured the levels of anti-Cas9 and anti-PEG IgG antibodies in mouse serum using mouse anti-Cas9 ELISA kits (Novatein Biosciences, NB-E1372PR) and mouse anti-PEG IgG ELISA kits (Life Diagnostics, PEGG-1). The cytokine and inflammatory biomarker, IP-10, was assessed utilizing the Mouse IP-10 ELISA Kit (Abcam, ab260067). TNF-α, was assessed utilizing the Mouse TNF-α ELISA Kit (Abcam, ab208348). IL-6, was assessed utilizing the Mouse IL-6 ELISA Kit (BioLegend, 431307). IL1-β, was assessed utilizing the Mouse IL-1β High Sensitivity ELISA Kit (MULTI SCIENCES, EK201BHS).

### Quantification of ABE8e mRNA and sgRNA levels in mice plasma

Mice plasma samples were homogenized using RLT Lysis buffer (Qiagen, 79216). The diluted lysate underwent reverse transcription and PCR using the HiScript III U + One Step qRT-PCR Probe Kit (Vazyme, Q225-01) following the manufacturer’s instructions. A custom primer-probe mix specific for the 3’-UTR of ABE8e mRNA was utilized on a CFX96 Real-Time PCR Detection System. Purified ABE8e mRNA and sgRNA were employed for standardization. The details of the primers and probes used are provided in Supplementary Data Table 14.

### Isolation of protein corona

Mouse plasma was centrifugated at 13000 × g at 4°C to remove protein aggregates before use. LNP-168-Luc were mixed with an equal volume of C57BL/6 mouse plasma to mimic the protein concentration *in vivo* and incubated for 1 hour at 37°C under shaking^18^. The protein corona-coated LNP-168-Luc were isolated by centrifugation at 13000.0 × g for 30 min, followed by washing with cold PBS three times to remove unbound proteins. The same procedure was performed for plasma aliquots without adding the LNP-168-Luc to verify the absence of protein precipitation. The experiments were conducted three times. The amount of protein in protein corona-coated LNP-168-Luc was determined using a BCA assay kit (Thermo Fisher Scientific, 23225). The obtained protein samples were further lyophilized and stored at -20°C for further experiment.

### Reduction, alkylation, and digestion of proteins

For each sample, 100 μg of protein was mixed in a 6:1 (v/v) ratio with sample loading buffer (250 mM Tris-HCl pH 6.8, 10% SDS, 50% Glycerol, 5% β-Mercaptoethanol, and 0.5% bromophenol blue; Sigma) and incubated at 95°C for 5 min. All samples were run in 10% SDS- PAGE gels until the dye front was 1 cm from the bottom for higher reproducibility. The gels were washed with deionized water. Each lane was cut into cubes of ≈ 1.0 mm2. Gel cubes were transferred to 1.5 mL tubes (Axygen) and destained twice using 50% acetonitrile (ACN, Merck) in 25 mM NH_4_HCO_3_ (Aladdin) at room temperature and dehydrated with 100% ACN for additional 10 min with shaking. Dehydrated gel pieces were reduced with 10 mM DTT (Thermo Fisher Scientific, R0861) at 37°C for 60 min and then dehydrated with 100% ACN for additional 10 min, with shaking. Alkylation was conducted by replacing the DTT solution with 25 mM IAA (Sigma-Aldrich, 16125-25G) and incubated at room temperature for 20 min in the dark followed by dehydrating with 100% ACN for 10 min. The gel pieces were digested with 2 ng μL^-1^ of trypsin (Thermo Fisher Scientific, R001100) in 25 mM NH_4_HCO_3_ overnight at 37°C. Peptides were extracted from gel pieces with consecutive incubations: 65% ACN (ACN:H_2_O:formic acid=65:35:5) with 5 min sonication in a water bath and 3 min incubation at 37°C; 100% ACN for 30 min at 37°C. Supernatants were combined in a fresh vial, dried using a vacuum centrifuge at 50°C, and resuspended in 100 μL of 0.1% formic acid. All samples were desalinated using Pierce C18 Tips (Thermo Fisher Scientific).

### Mass spectrometry

The resulting samples were further analyzed by electrospray liquid chromatography–tandem mass spectrometry using an Orbitrap Exploris 480 (Thermo Fisher Scientific) coupled online to an EASY-nLC 1200 (Thermo Fisher Scientific). Samples were run on a Thermo Fisher Scientific PepMap RSLC C18 EASY-Spray Column (3 μm particle size, 75 μm × 15 cm, 100 Å) at a flow rate of 250 nL min^-1^ using the following gradient: 0 to 5 min 2 to 8% B, 5 to 97 min 8 to 22% B, 97 to 110 min 22 to 35% B, 110 to 111 min 35 to 90% B, 111 to 120 min at 80% B. Mobile phase A was 0.1% formic acid (FA) in water. Mobile phase B was 0.1% FA in 80% ACN. The data were analyzed using the Proteome Discoverer 2.1.0.81 software (Thermo Fisher Scientific). The data were run against the Mus musculus Uniprot FASTA file to identify the mouse proteins in the corona.

### Liquid chromatography-tandem mass spectrometry quantification of Lipid-168

Blood samples for the quantitation of Lipid-168 were drawn into K2EDTA tubes, processed to plasma, and snap frozen in liquid nitrogen. All samples were stored at -80℃ until analysis. Added 5 μL of plasma samples to the 96-well plate, then mixed with 200 μL of IS solution (0.1% FA in MeOH (with 600 ng/ml Tolbutamide)) to induce protein precipitation. Then the 96-well plate was vortexed for 15 min on the shaker and centrifugated (10 min at 4000 rpm), after that, the 20 μL of supernatants were transferred to a new 96-well plate, added 300 μL of water and mixed well. A standard curve Lipid-168 was created covering the concentration range of 10-20,000 ng/ml. Samples expected to exceed these concentrations were diluted accordingly. Concentrations were determined by multiple reaction monitoring using a high-performance liquid chromatography (HPLC) triplequadrupole mass spectrometer (LC-40, SHIMADZU; SCIEX QTRAP® 6500+, Sciex, Framingham, MA, USA). Plasma concentrations were determined from the standard curve using Analyst 1.7.3 (Sciex, Framingham, MA, USA).

## ANIMAL ETHICS STATEMENTS

The animal experiments were approved by the ECNU Animal Care and Use Committee and in direct accordance with the Ministry of Science and Technology of the People’s Republic of China on Animal Care guidelines. The protocol (protocol ID: m20210238) was approved by the ECNU Animal Care and Use Committee. All mice were euthanized after the completion of the experiments.

## ETHICS STATEMENT

Healthy donors and β-thalassemia patient CD34^+^ HSPCs were isolated from plerixafor-mobilized or unmobilized peripheral blood following Xiangya Hospital Central South University and the First Affiliated Hospital of Guangxi Medical University with review board (IRB) approval and informed patient consent.

## Supporting information

Synthetic routes and characterization data for ionizing lipids.

## ACKNOWLEDGMENTS

We acknowledge R. Kurita and Y. Nakamura for providing the HUDEP-2 cells. We appreciate Min Qiu for offering valuable advice on LC-MS detection and data analysis. The flow cytometry analysis experiments were performed at the Flow Cytometry Core Facility of School of Life Sciences of ECNU. We thank the staff member Ying Zhang for help with flow cytometry data collection. This work was supported by the National Key R&D Program of China 2019YFA0109900 (Y.W.), 2019YFA0109901 (Y.W.), 2019YFA0802800, and 2019YFA0110803 (Y.W.), grants from the Shanghai Municipal Commission for Science and Technology 19PJ1403500 (Y.W.), the National Science Foundation of China grants 82270125(Y.W.), the Scientific Research of BSKY XJ2020025(D. L.). from Anhui Medical University, and and the Anhui Province Fund for Excellent Young Scholars 2024AH030022 (D. L.).

## AUTHOR CONTRIBUTIONS

Y.L., Z.W. and Y.W. conceived and supervised this study. S.X. and D.L. conducted most of the cell and animal experiments and analyzed data with the help of Y.C., F.Z., H.Z. and G.S. Q.W. and C.F. performed the bioinformatics analysis. W.R. and D.X. conducted the mRNA and LNPs production. L.L. assisted with the proteomics analysis. Z.W. and Y.G. designed and synthesized the library of ionizable lipids. Y.Y., Y.L. and B.F. recruited β-thalassemia patients to harvest CD34^+^ cells. S.X., D.L., Y.L., Z.W. and Y.W. wrote the manuscript. All authors contributed to the manuscript and approved its final version.

## DECLARATION OF INTERESTS

Y.L., Z.W. and Y.W. are scientific co-founders in Yoltech Therapeutics. D.X., W.R., Y.G., H.Z. and C.F. are employees in Yoltech Therapeutics. Yoltech Therapeutics has filed patents (CN114989182A, CN116162071A, PCT/CN2023/100791, PCT/CN2023/100823, and PCT/CN2023/106421) on the technology described in this manuscript. The remaining authors declare no competing interests.

## DATA AND MATERIALS AVAILABILITY

All datasets generated during this study are included in this manuscript.

---

**Extended Data Fig.1.**
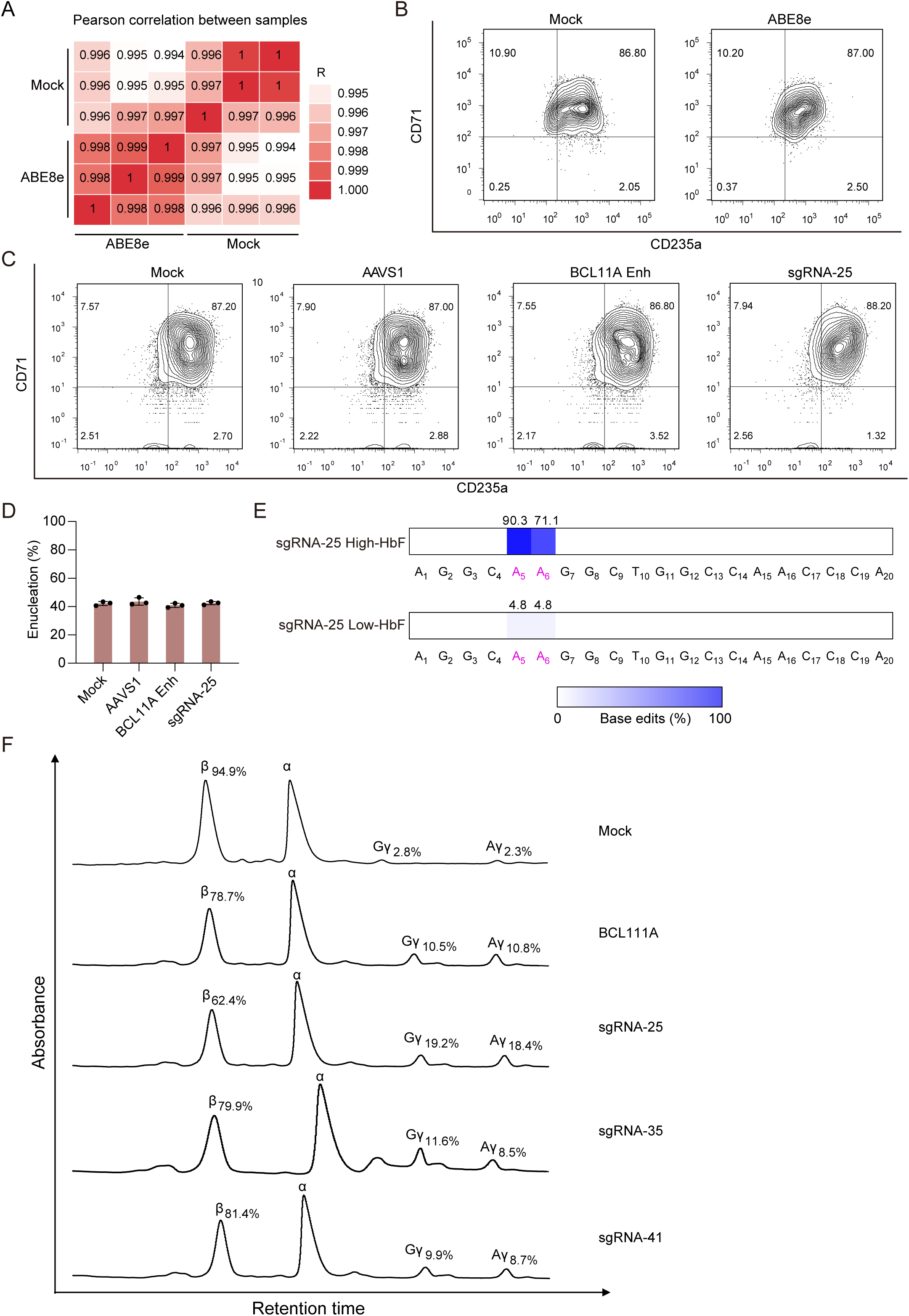
| Highly efficient editing of HBG promoter in CD34^+^ HSPCs. **A.** Heat map showing the pearson correlation of mock and ABE8e edited cells transcriptomes as measured by RNA-seq. The number in each cell is the correlation coefficient. **B.** Flow cytometry plots showing the expression of the erythroid maturation markers CD71 and CD235a *in vitro* differentiation of HUDEP-2 cells. **C.** Flow cytometry plots showing the expression of the RBCs maturation markers CD71 and CD235a after *in vitro* differentiation of ABE8e RNP edited CD34^+^ cells. **D.** Enucleation rates in erythroid cells after *in vitro* differentiation of ABE8e RNP edited and un-edited CD34^+^ cells. **E.** Base-editing rates in bulk base-edited cells in high and low HbF cells after *in vitro* differentiation of ABE8e RNP edited CD34^+^ cells. The T-C conversion rate of sgRNA-25 in CD34^+^ cells was measured by deep sequencing and quantified with *CRISPResso2*. F. Representative RP-HPLC chromatograms of erythroid cells derived from *in vitro* differentiation of edited CD34^+^ HSPCs. Protein levels of β-like globins by genome editing mediated by ABE8e-BCL11A Enh, ABE8e-sgRNA-25, ABE8e-sgRNA-35 and ABE8e-sgRNA-41 were analyzed. Three independent experiments were performed. Data are plotted as the mean ± s.d.

**Extended Data Fig.2.**
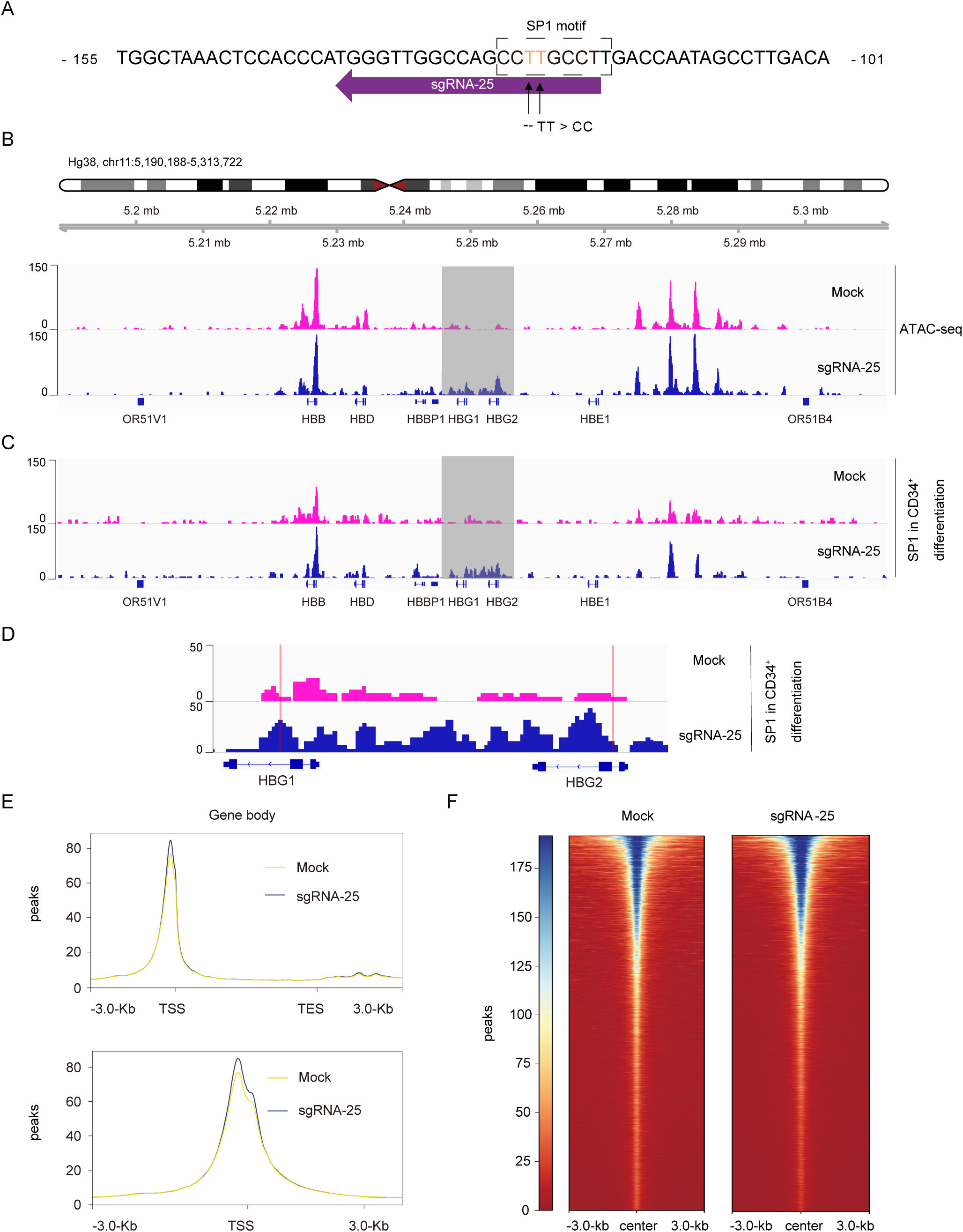
| SP1 rapidly activates γ-globin after base editing of sgRNA-25. **A.** Schematic diagram of the sequence of the γ-globin promoter of CD34^+^ promoter editing cells. The left arrow indicates the sequence position of sgRNA-25, and the two upward arrows indicate the targeted bases. **B.** ATAC-seq of *in vitro* differentiation of ABE8e RNP edited and un-edited CD34^+^ cells. **C.** SP1 CUT& Tag profiles in the β-globin cluster. Antibodies and cell types for each track are shown on the right. The promoters of duplicated γ-globin genes (HBG2 and HBG1) are highlighted in gray. **D.** Zoomed-in view of the HBG2 and HBG1 promoter regions. The sequence of sgRNA-25 is highlighted in orange. Heatmap comparison of signals in primary human CD34^+^-derived erythroid cells with or without sgRNA-25 base editing within binding sites. **E.** The enrichment curve of the CUT&Tag binding signal in the upstream and downstream 3-kb regions of the gene body and TSS region. **F.** Heat map comparison of overlapping peaks CUT&TAG.

**Extended Data Fig.3.**
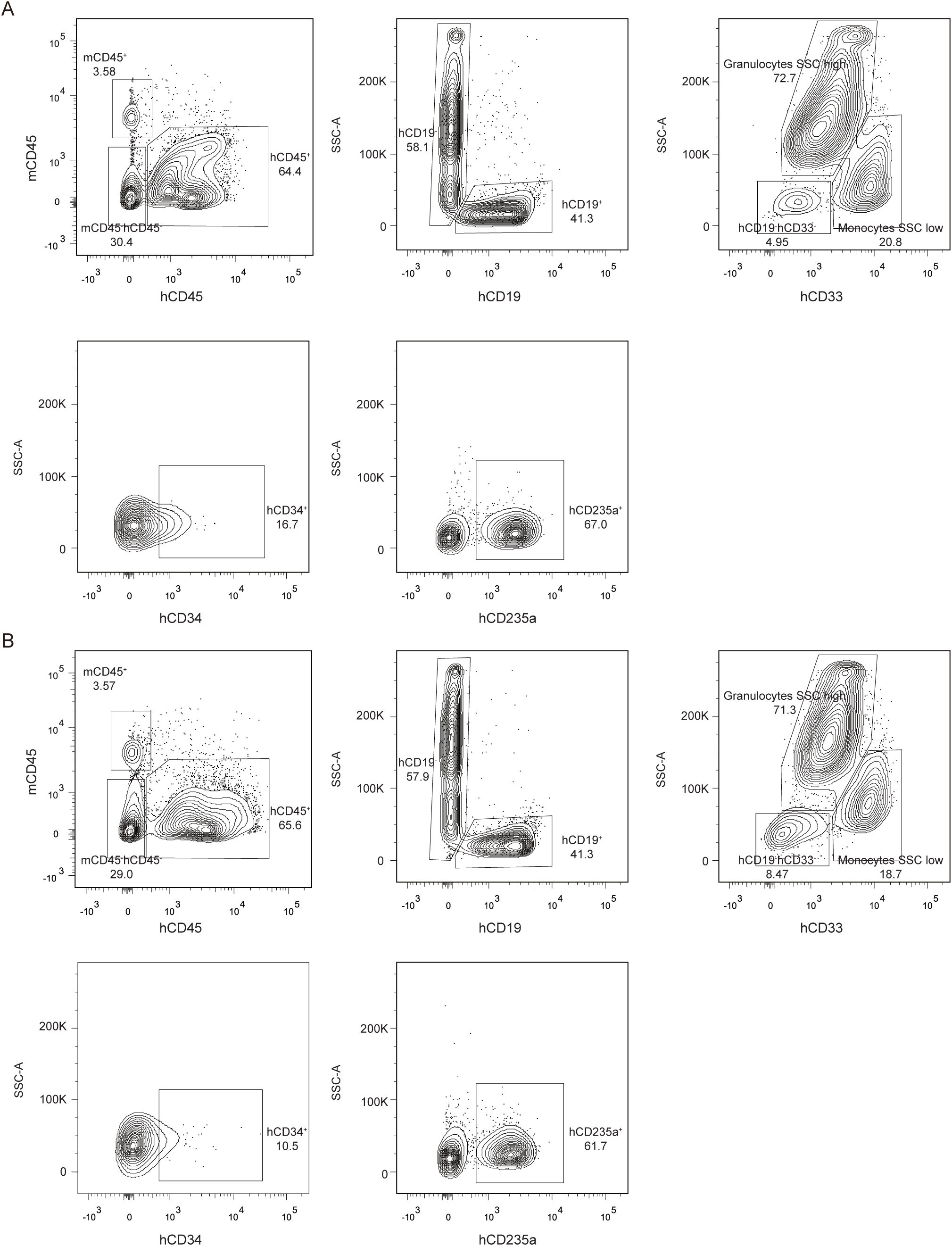
| Representative xenografted BM flow cytometry analysis. **A-B.** Live cells from engrafted mouse BM. Human cells gated from hCD45^+^ population, mouse cells gated from mCD45^+^ population. B cells gated from hCD45^+^CD19^+^ population. Granulocytes gated from hCD45^+^CD19^-^CD33dim with SSC high population. Monocytes gated from hCD45^+^CD19^-^CD33^+^ with SSC low population. CD34^+^ cells gated from hCD45^+^CD19^-^CD33^-^CD34^+^population. Erythroid cells gated from hCD45^-^mCD45^-^hCD235a^+^ population.

**Extended Data Fig.4.**
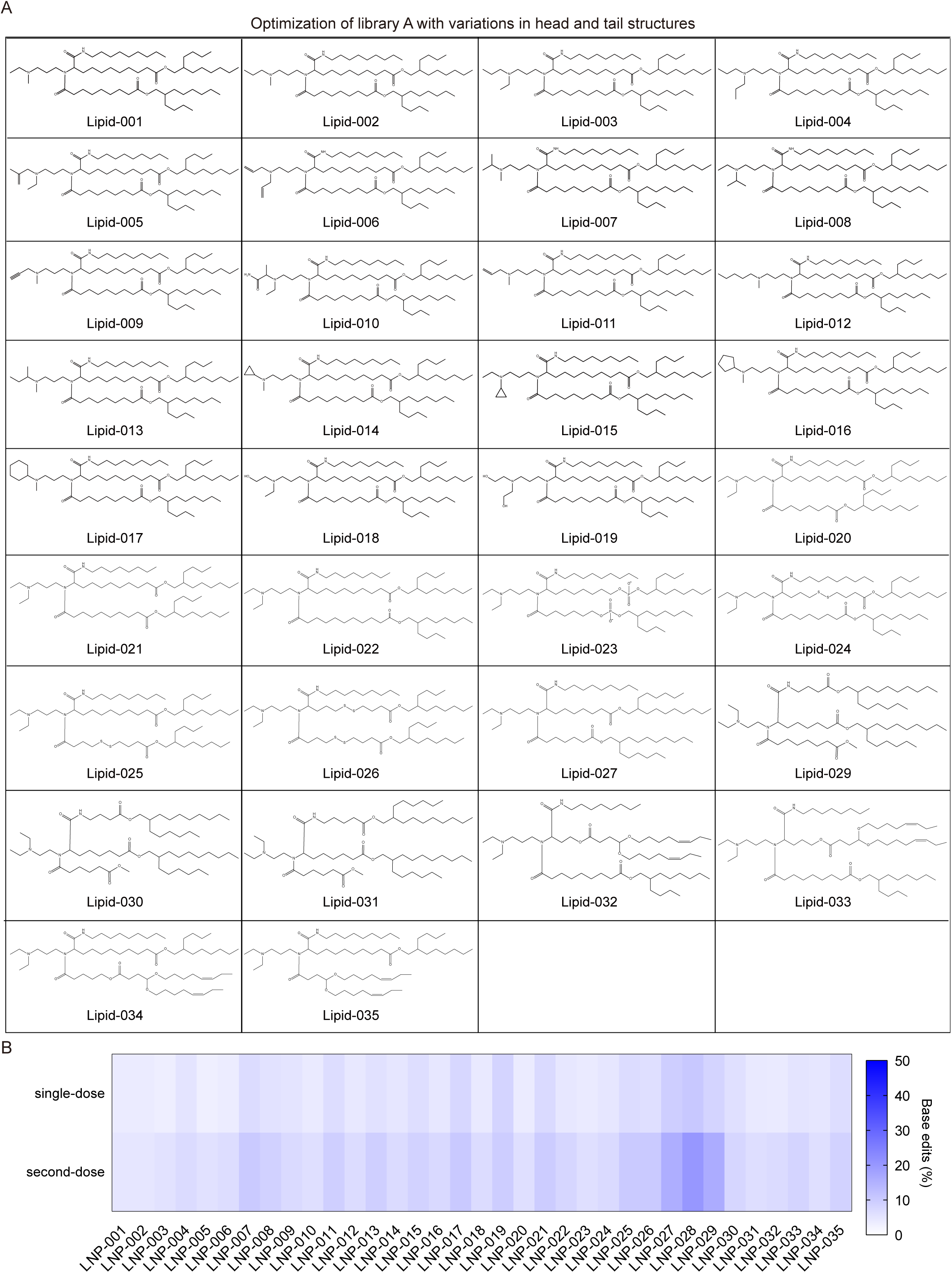
| Structures of ionizable lipids in Library A and the editing efficiency of BM cells in mice treated with different LNPs. **A.** Structures of various headgroups and tails in a combinatorial library A of ionizable lipids. **B.** The top-performing ionizable lipid structures within library A were determined by analyzing the editing efficiency of BM cells in mice treated with 35 different LNP-ABE8e-PCSK9 formulations (intravenous injection, 2 mg/kg of LNP-ABE8e-PCSK9, 3-4 mice per LNP mixture).

**Extended Data Fig.5.**
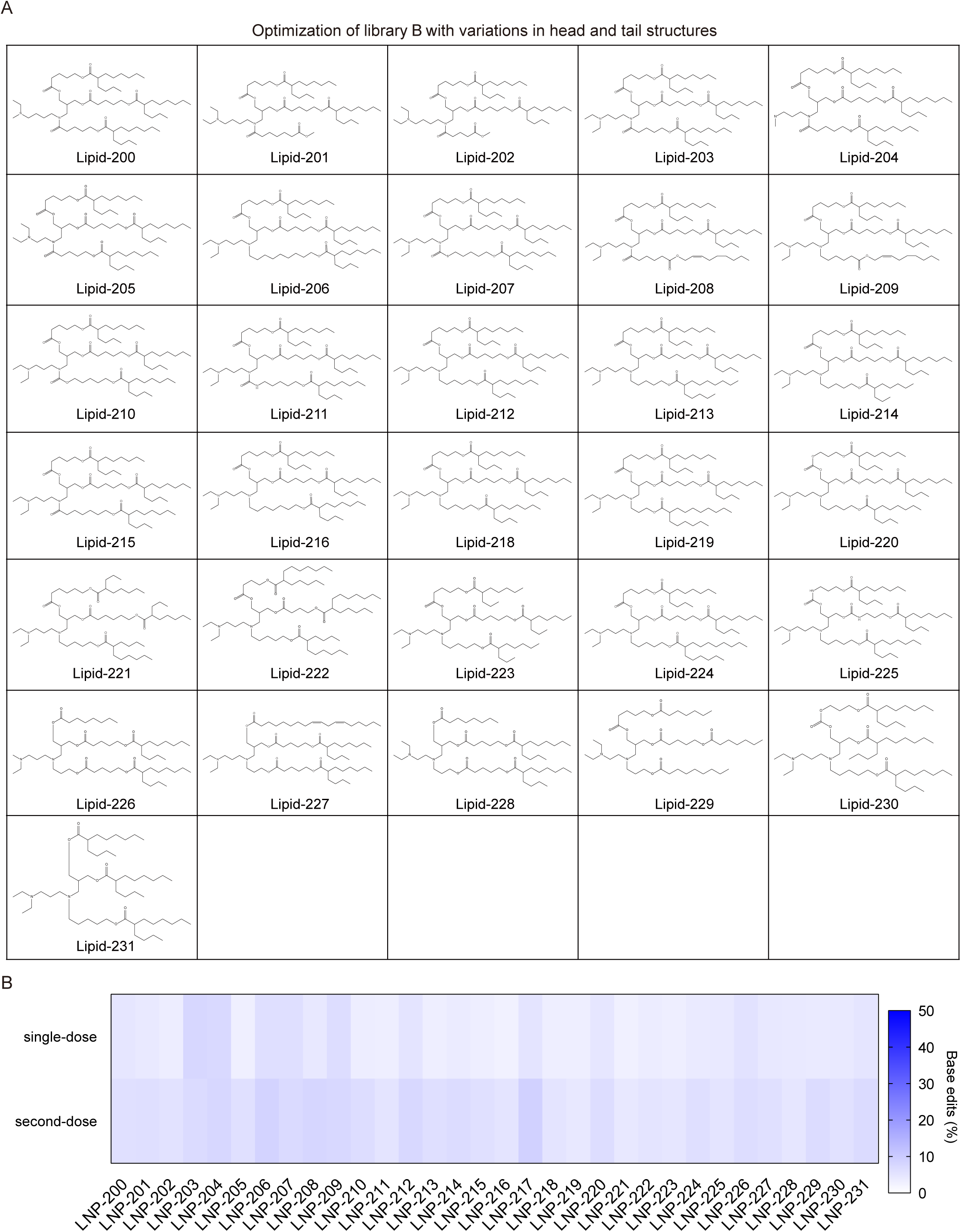
| Structures of ionizable lipids in Library B and the editing efficiency of BM cells in mice treated with different LNPs. **A.** Structures of various headgroups and tails in a combinatorial library B of ionizable lipids. **B.** The top-performing ionizable lipid structures within library B were determined by analyzing the editing efficiency of BM cells in mice treated with 32 different LNP-ABE8e-PCSK9 formulations (intravenous injection, 2 mg/kg of LNP-ABE8e-PCSK9, 3-4 mice per LNP mixture).

**Extended Data Fig.6.**
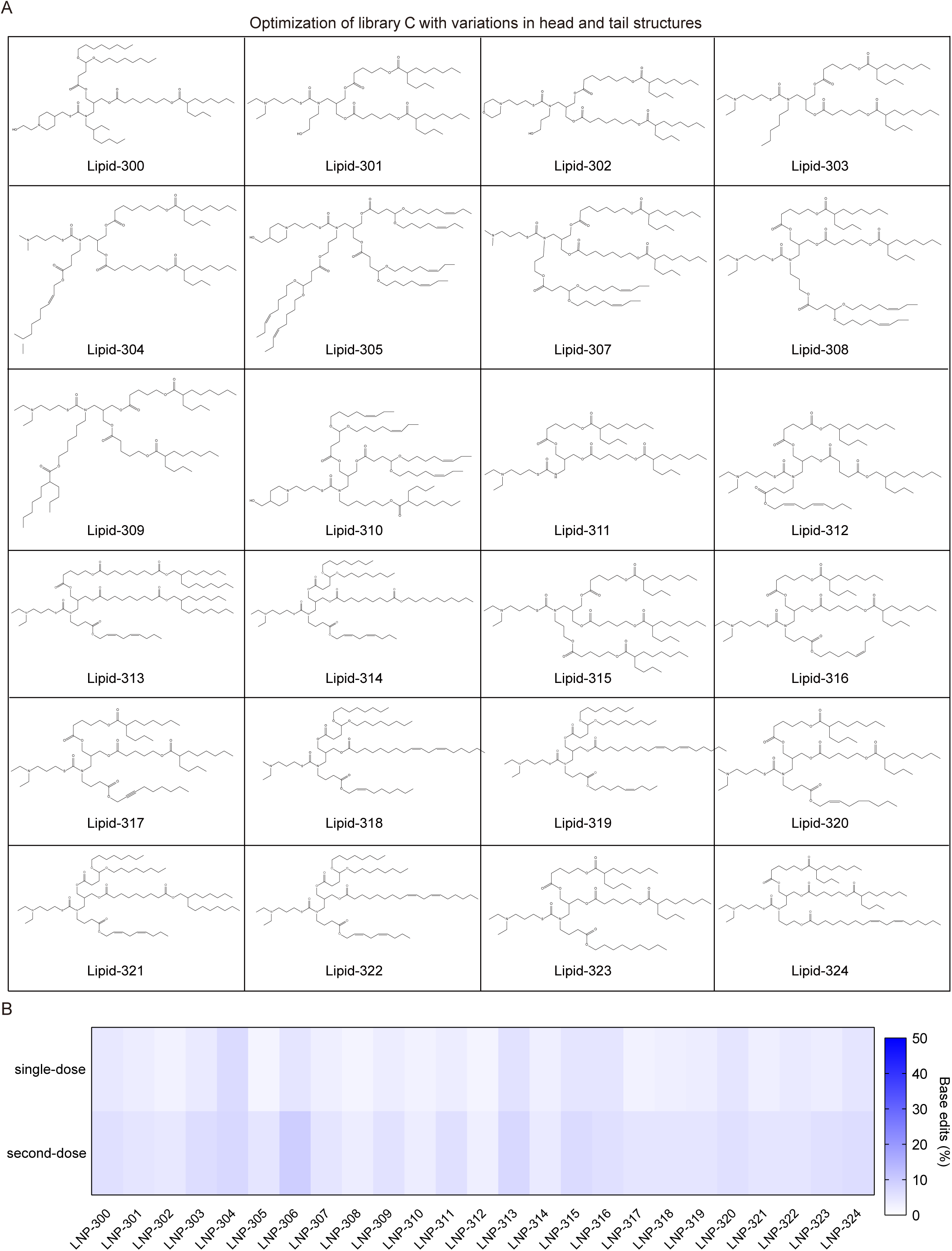
| Structures of ionizable lipids in Library C and the editing efficiency of BM cells in mice treated with different LNPs. **A.** Structures of various headgroups and tails in a combinatorial library C of ionizable lipids. **B.** The top-performing ionizable lipid structures within library C were determined by analyzing the editing efficiency of BM cells in mice treated with 25 different LNP-ABE8e-PCSK9 formulations (intravenous injection, 2 mg/kg of LNP-ABE8e-PCSK9, 3-4 mice per LNP mixture).

**Extended Data Fig.7.**
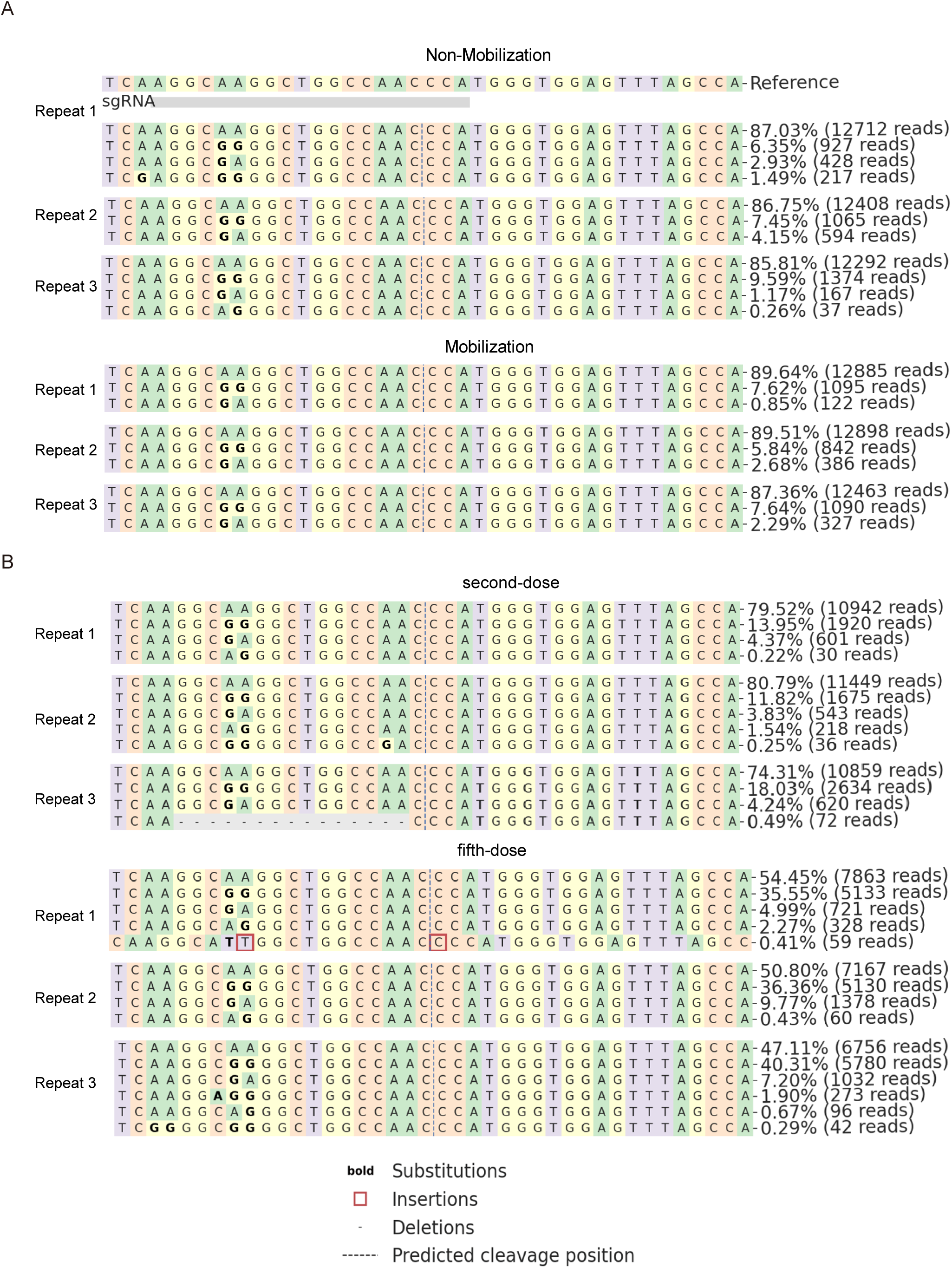
| Base editing tables the HBG promoter in the BM cells after LNP-028-ABE8e-HBG injection. **A.** Next-generation sequencing of BM cells from mice of mobilized and non-mobilized groups after LNP-028-ABE8e-HBG base edit. **B.** Next-generation sequencing of BM cells from mice of second-dose and fifth-doses groups after LNP-028-ABE8e-HBG base edit.

**Extended Data Fig.8.**
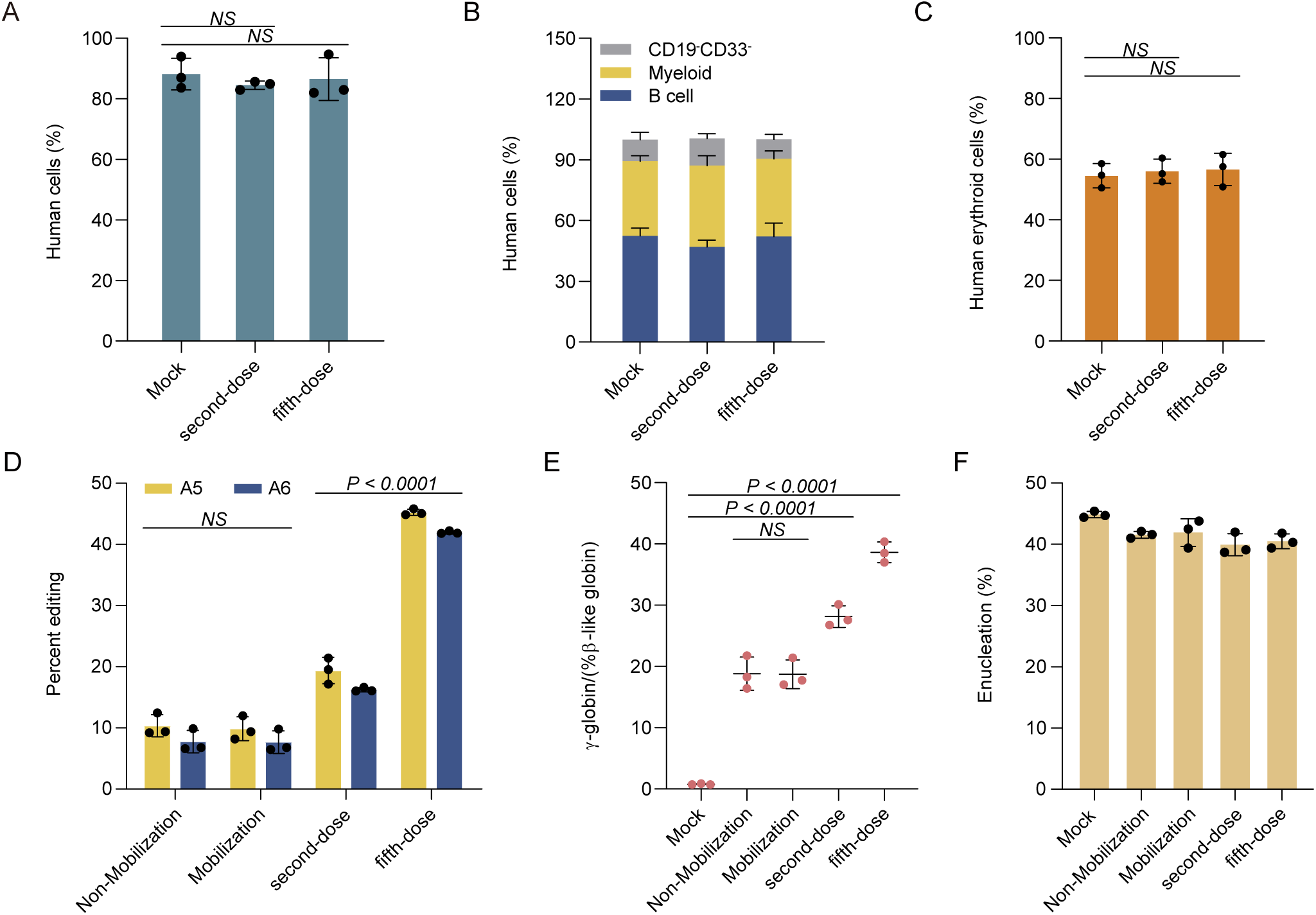
| Long-term multilineage engraftment of LNP-028-ABE8e-HBG edited HSPCs in immunodeficient mice. **A-C.** BM was collected 2 weeks after injection of LNP-028-ABE8e-HBG and analyzed by flow cytometry for human cell chimerism (**A**), and multilineage reconstitution (**B**) or human erythroid cells (**C**) in the BM. **D.** Base editing efficiency was assessed *in vitro* in differentiated erythroid cells derived from BM cells of engrafted mice after LNP-028-ABE8e-HBG injection. **E.** γ-globin expression was analyzed by RT-qPCR in erythroid cells differentiated *in vitro* from LNP-028-ABE8e-HBG edited BM cells. **F.** Enucleation of *in vitro* differentiated erythroid cells from LNP-028-ABE8e-HBG edited BM cells. Each symbol represents a mouse, and there were a total of n=3 mice. All statistical significances in the figures were analyzed using one-way ANOVA with Dunnett’s multiple comparisons test, and data represent mean ± SD, with *p*-values noted where appropriate. *NS*, not significant.

**Extended Data Fig.9.**
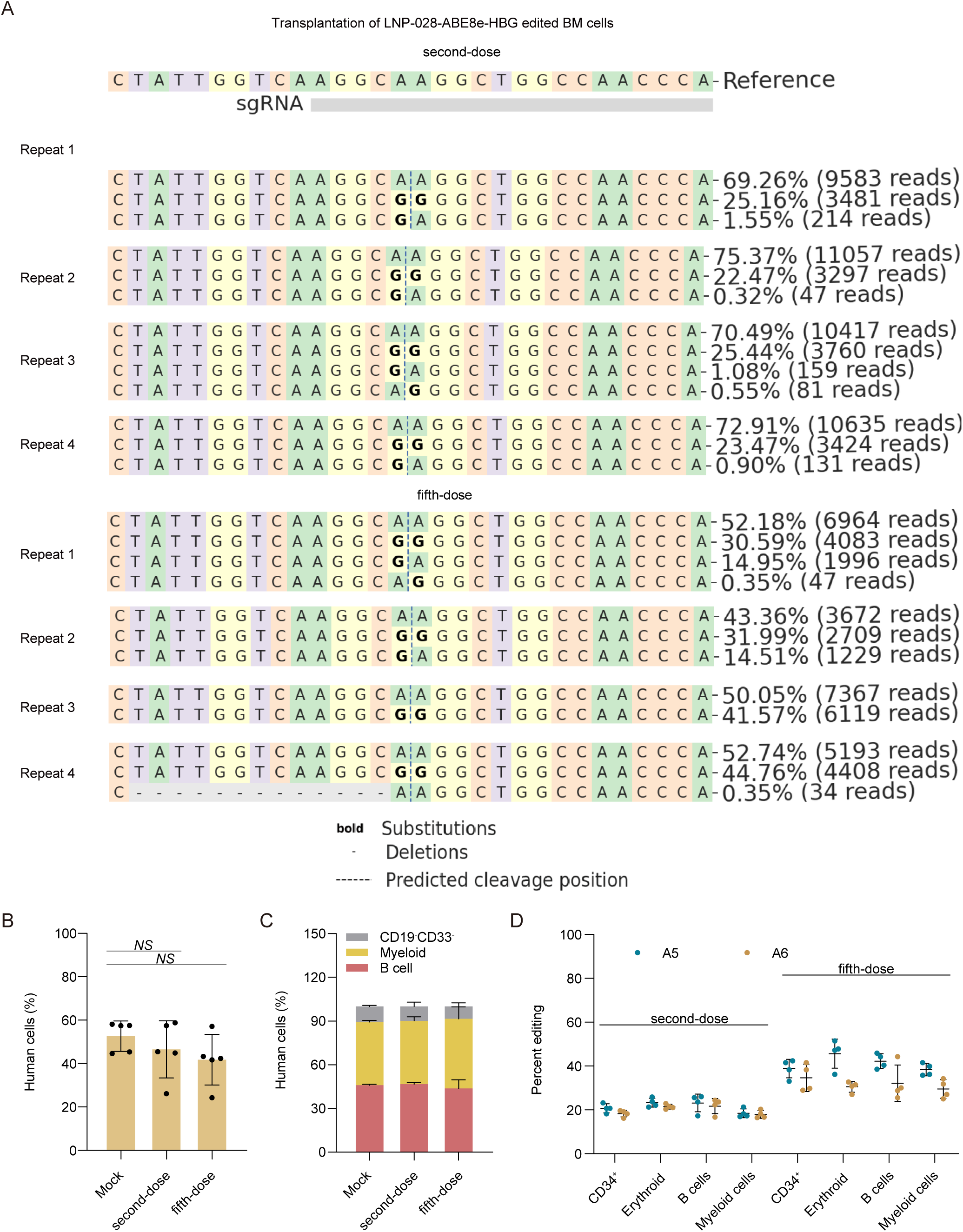
| Efficient HBG promoter base editing in HSCs after transplantation of LNP-028-ABE8e-HBG edited BM cells. **A.** After transplanting BM cells edited with LNP-028-ABE8e into *NCG-X* mice for 16 weeks, the editing efficiency of BM cells was analyzed through deep sequencing. **B.** After transplanting LNP-028-ABE8e-HBG edited BM cells into *NCG-X* mice for 16 weeks, the chimerism level of human cells in BM cells was analyzed by flow cytometry. **C.** Percentage of engrafted human B cells, myeloid cells, and CD19^-^CD33^-^ cells in base-edited BM cells were assessed after 16 weeks of transplantation using flow cytometry. **D.** After 16 weeks of transplantation, deep sequencing analysis was conducted to evaluate the base editing efficiency at positions A5 and A6 within different hematopoietic cell populations in BM cells.

**Extended Data Fig.10.**
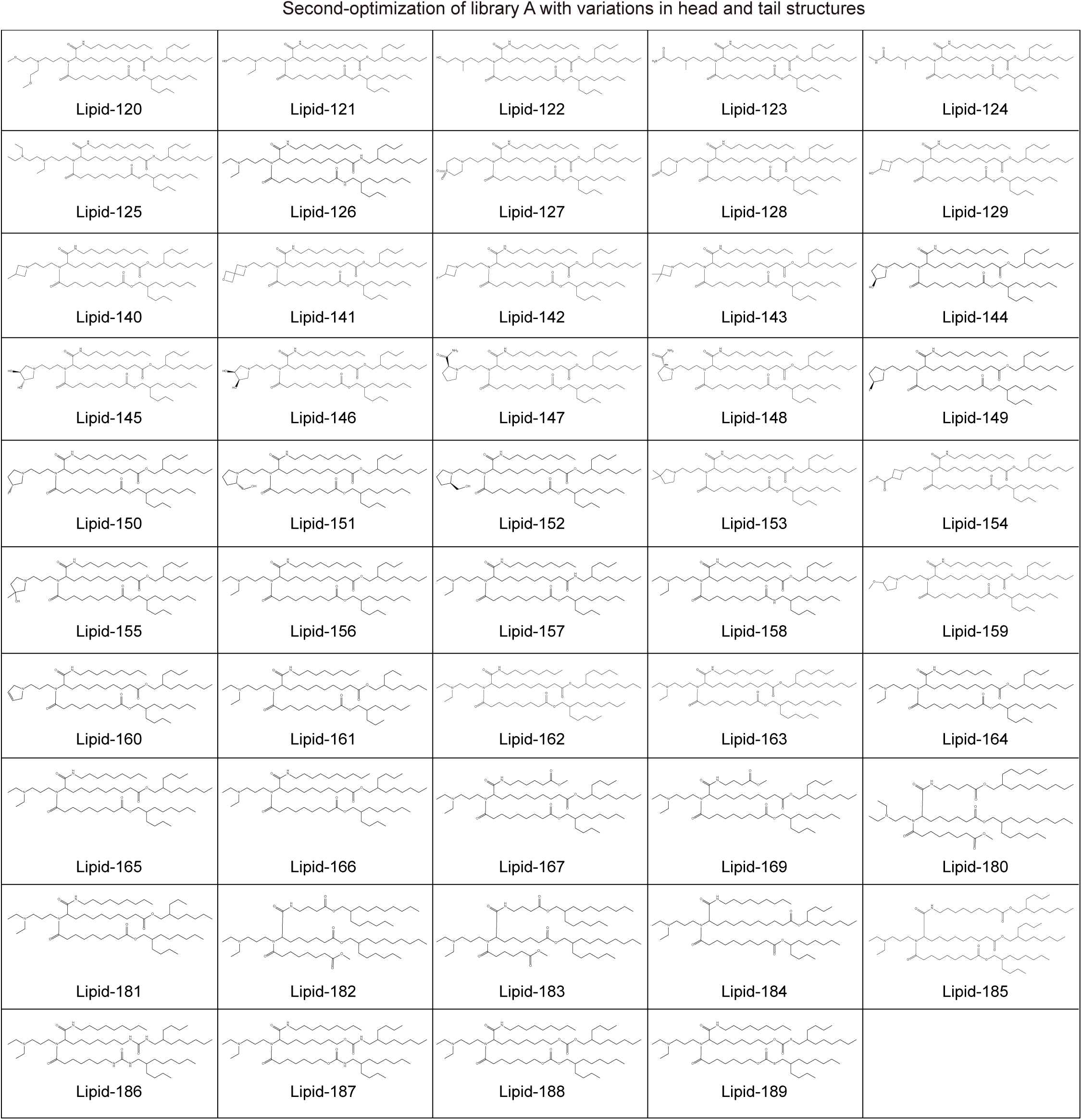
| Structures of ionizable lipids in the second optimization of Library A. Structures of various headgroups and tails in a second combinatorial library A containing 49 different ionizable lipids.

**Extended Data Fig.11.**
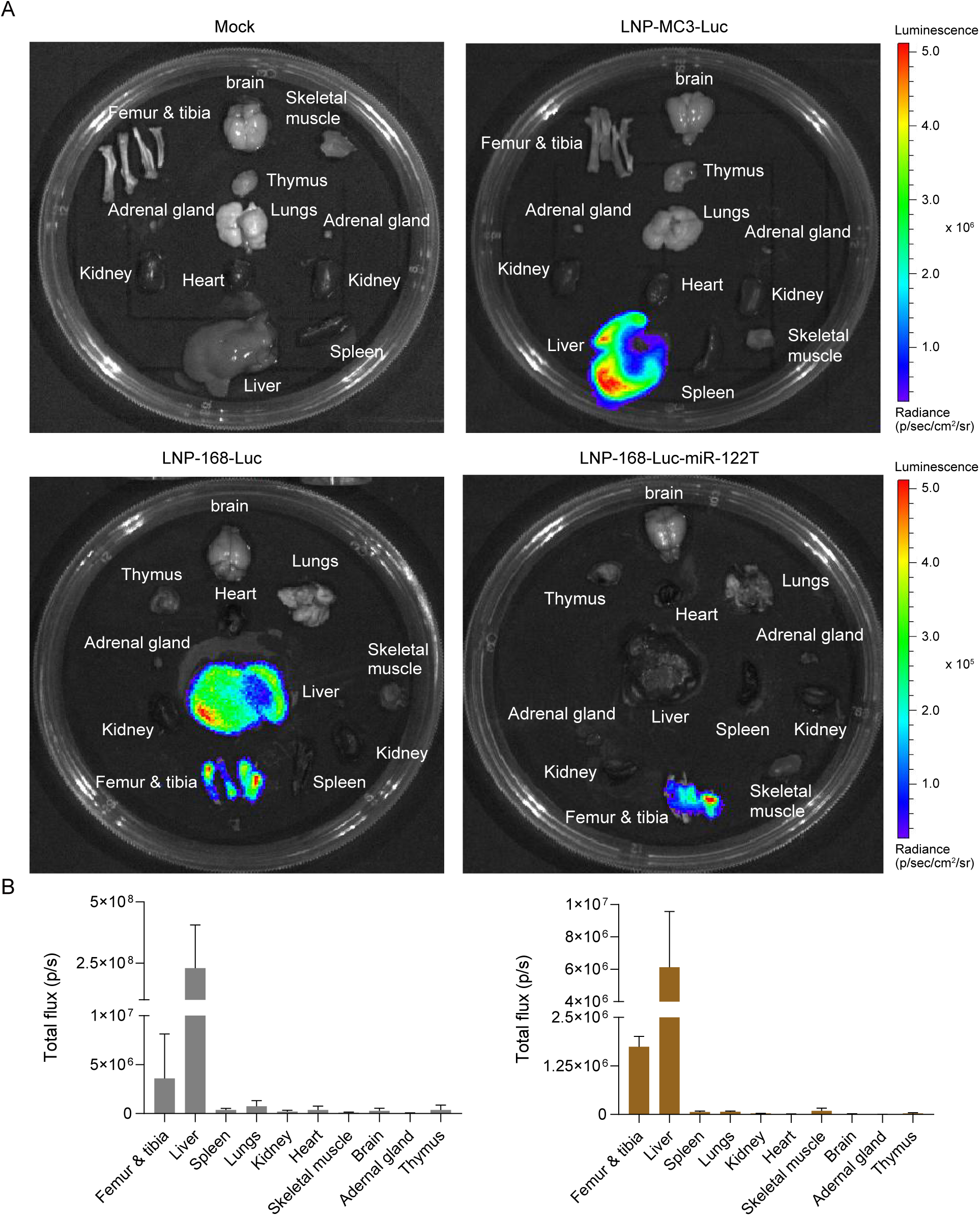
| Biodistribution of i.v. administration of LNP-168-Luc. **A.** Representative IVIS bioluminescence images of all organs after 24 h. LNP-168-Luc were injected at a dose of 1 mg/kg. A representative sample set of various tissues and organs dissected from these mice was analyzed 10 minutes after the administration of D-luciferin. **B.** Quantification of the total flux intensity in the LNP-168-Luc and LNP-MC3-Luc groups (n=3 mice).

**Extended Data Fig.12.**
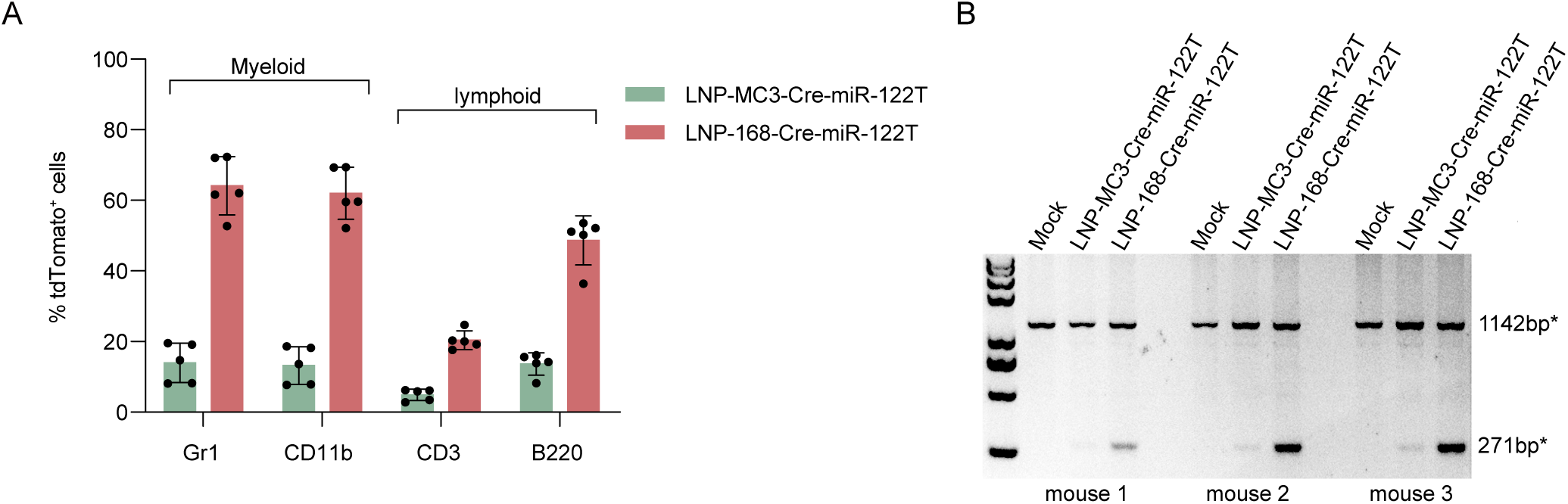
| *In vivo* LNP-168-Cre editing of BM cells in Ai14 mice. **A.** The tdTomato^+^ cells expression in mouse lymphoid, Gr1^+^ or CD11b^+^ myeloid cells in BM cells from Ai14 mice after administration of LNP-Cre. **B.** Semiquantitative PCR of genomic DNA isolated from BM cells of A14 mice after administration of LNP-Cre. * 271bp, Cre-recombinase edited gDNA region and * 1142bp, unedited region is indicated.

**Extended Data Fig.13.**
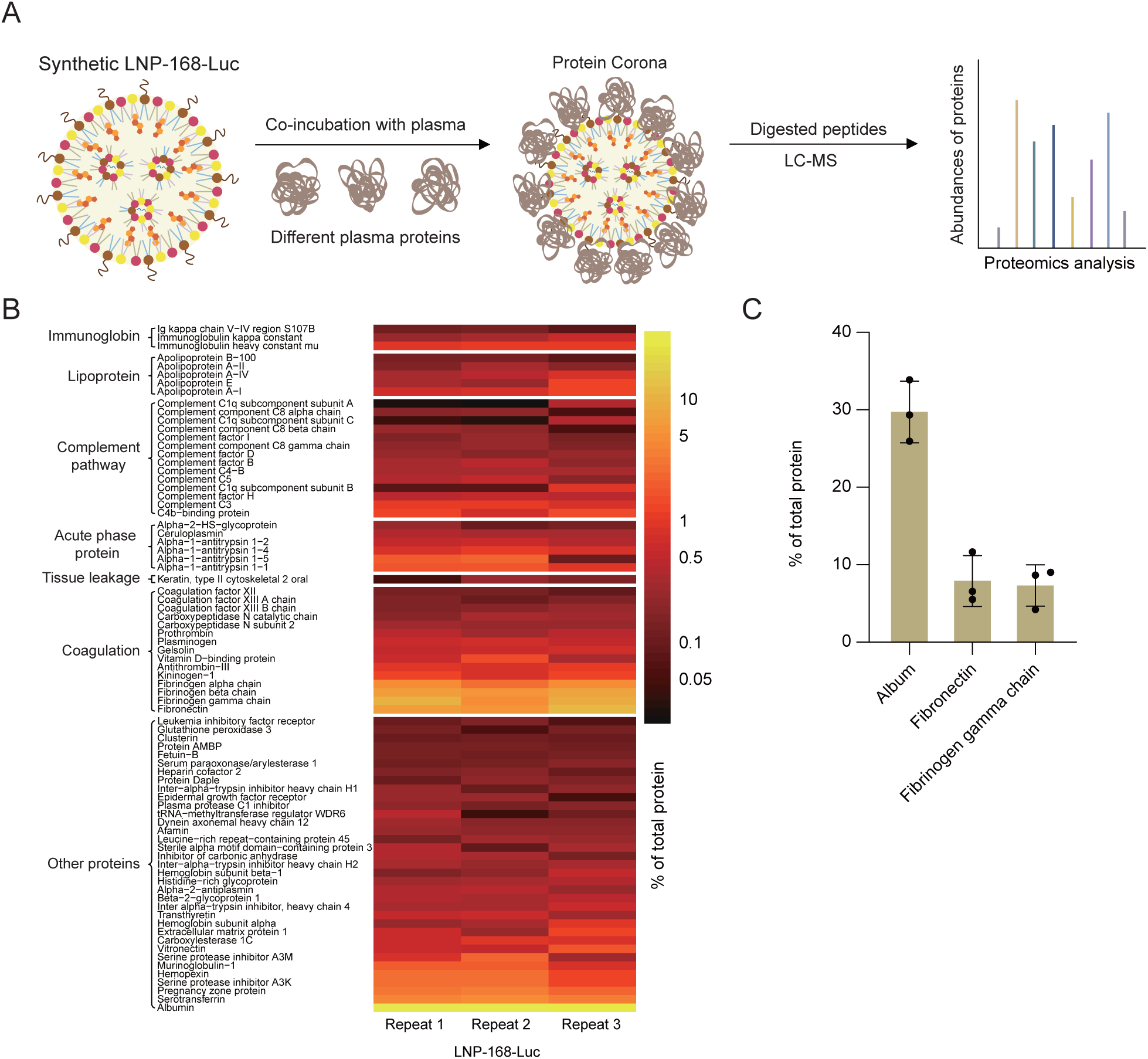
| Proteomics study of protein coronas formed on LNP-168-Luc. **A.** Schematic showing the use of mass spectrometry in the analysis of LNPs mRNA with proteins in the plasma interactions. **B.** Heat map of the average abundance of proteins with distinct biological functions in the protein coronas of LNP-168-Luc. **C.** Quantification of percentage of total proteins of the top three protein components in the protein corona of the LNP-168-Luc is shown (n=3 mice).

**Extended Data Fig.14.**
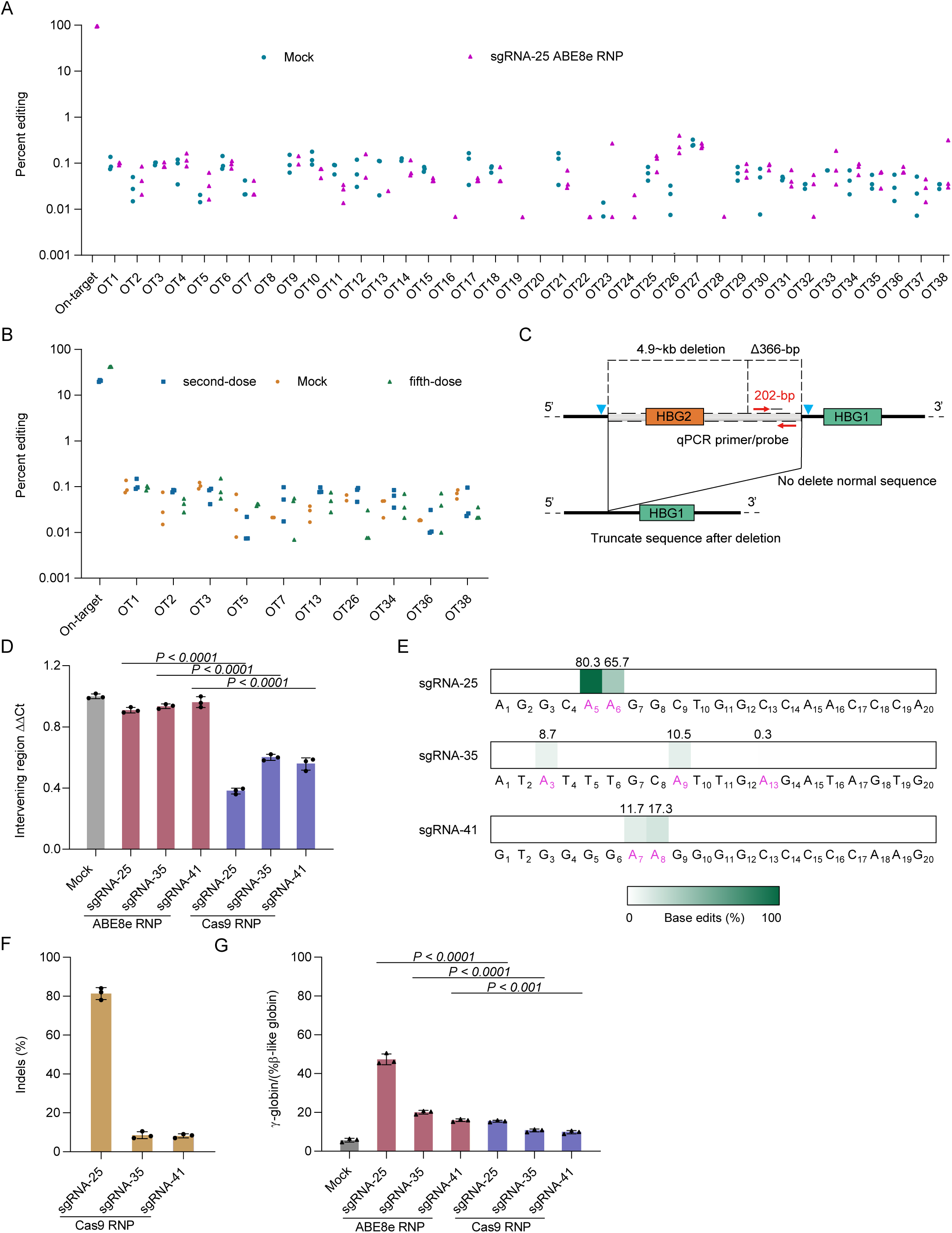
| Assessment of off-target editing *in vitro* and *in vivo*. **A**. Total 38 potential genomic sgRNA-25 off-target sites were evaluated by amplicon deep sequencing and quantified with *CRISPResso2*. Some off-target detection values in the experiment were zero and were not displayed in the graph. **B.** Ten potential genomic sgRNA-25 off-target sites were evaluated by amplicon deep sequencing in BM cells from engrafted mice after LNP-028-ABE8e-HBG injection. Some off-target detection values in the experiment were zero and were not displayed in the graph. **C.** HBG ABE8e cleavage sites are indicated by inverted triangle. Quantitative PCR (qPCR) primers targeting the intergenic sequence between the cleavage sites are indicated (red arrows). The larger deletion presumed to arise from simultaneous ssDNA breaks in HBG2 and HBG1, with the loss of the intervening 4.9-kb, is shown (dotted line). Primers for △366-bp denotes a quantitative qPCR assay that detects deletions ≥366-nt upstream of the cleavage site in HBG1. **D.** Deletion analysis after RNP electroporation sgRNA expression in human CD34^+^ cells. n=[3 replicates from independent electroporations. **E.** Base editing in CD34^+^ HSPC donors by ABE8e RNP after days 5. Base editing was measured by deep sequence analysis. Five days after Cas9 RNP editing, the indel efficiency in donor CD34^+^ cells were examined. **F.** Indel frequency was analyzed by *synthego*. **G.** β-like globin expression by RT-qPCR analysis in erythroid cells differentiated *in vitro* from RNP-edited CD34^+^ HSPCs. All statistical significances in the figures were analyzed using one-way ANOVA with Dunnett’s multiple comparisons test, and data represent mean ± SD, with *p*-values noted where appropriate.

