## Supplementary material for "In vivo genome editing of human hematopoietic stem cells for treatment of blood disorders by mRNA delivery": Synthetic routes and characterization data for ionizing lipids.

### EXAMPLE 1 Synthesis of Lipid-028

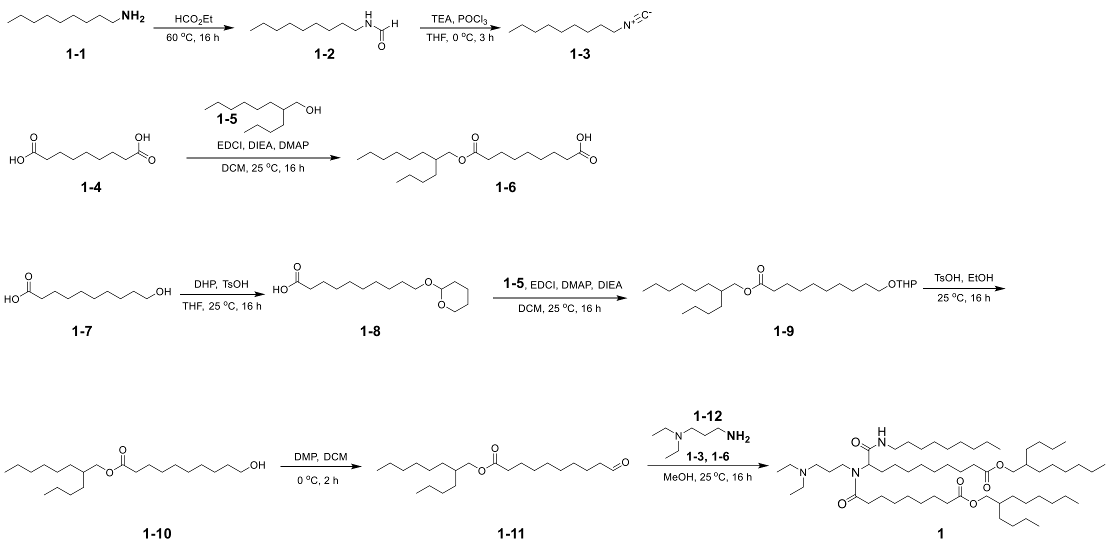

Intermediate **1-2**: *N*-nonylformamide

The reaction solution of nonan-1-amine (10.00 g, 77.38 mmol, 1.0 eq) in ethyl formate (30 mL) was stirred at 60 ºC for 16 hours. The reaction mixture was concentrated under reduced pressure. The desired product was obtained as a colorless oil (11.96 g, 100% yield) which can be used for the next step without further purification.

Intermediate **1-3**: 1-isocyanooctane

Phosphorus oxychloride (18.98 g, 123.81 mmol, 1.6 eq) was added to the solution of **1-2** (11.96 g, 77.38 mmol, 1.0 eq) and TEA (64.53 mL, 464.28 mmol, 6.0 eq) in THF (100 mL) slowly at 0 ºC. The reaction mixture was stirred for 3 hours and quenched by aqueous saturated $NaHCO_3$ (100 mL), then extracted with EtOAc (100 mL x 3). The organic layers were combined and dried over anhydrous $Na_2SO_4$, then concentrated under reduced pressure, the residue was purified by silica column (PE/EtOAc = 1/0 to 40/1), the desired product was obtained as a colorless oil (8.10 g, 68.3% yield).

Intermediate **1-6**: 9-((2-butyloctyl)oxy)-9-oxononanoic acid

The solution of nonanedioic acid (10.10 g, 53.65 mmol, 5.0 eq), DMAP (0.66 g, 5.37 mmol, 0.5 eq), DIEA (13.87 g, 107.30 mmol, 10.0 eq) and EDCI (3.09 g, 16.09 mmol, 1.5 eq) was stirred at 25 ºC for 0.5 hours before 2-butyloctan-1-ol (2.00 g, 10.73 mmol, 1.0 eq) was added. The reaction soltuion was stirred at 25 ºC for 15.5 hours and concentrated under reduced pressure. The residue was diluted with $H_2O$ (100 mL) and extracted with EtOAc (100 mL x 3), the organic layers were combined and washed with aqueous saturated NaCl solution, dried over anhydrous $Na_2SO_4$, then concentrated under reduced pressure. The residue was purified by silica column, 9-((2-butyloctyl)oxy)-9-oxononanoic acid was obtained as a colorless oil (3.00 g, 78.4% yield).

Intermediate **1-8**: 10-((tetrahydro-2*H*-pyran-2-yl)oxy)decanoic acid

The solution of 10-hydroxydecanoic acid (5.00 g, 26.56 mmol, 1.0 eq), DHP (3.35 g, 39.84 mmol, 1.5 eq), TsOH (0.23 g, 1.33 mmol, 0.05 eq) in THF (50 mL) was stirred at 25 °C for 16 hours. The reaction solution was concentrated, the residue was purified by silica column (PE/EtOAc = 10/1 to 1/1), the desired product was obtained as a colorless oil (6.30 g, 87.1% purity).

Intermediate **1-9**: 2-butyloctyl 10-((tetrahydro-2*H*-pyran-2-yl)oxy)decanoate

A solution of **1-8** (6.30 g, 23.13 mmol, 1.0 eq), **1-5** (6.46 g, 34.70 mmol, 1.5 eq), DMAP (1.41 g, 11.56 mmol, 0.5 eq), DIPEA (8.97 g, 69.39 mmol, 3.0 eq), EDCI (6.65 g, 34.70 mmol, 1.5 eq) in DCM (60 mL) was stirred at 25 °C for 16 hours. The reaction solution was concentrated under reduced pressure, the residue was purified by silica column (PE/EtOAc = 40/1 to 10/1), the desired product was obtained as a colorless oil (5.30 g, 52.0% yield).

Intermediate **1-10**: 2-butyloctyl 10-hydroxydecanoate

The solution of **1-9** (5.30 g, 12.03 mmol, 1.0 eq) and TsOH (2.07 g, 12.03 mmol, 1.0 eq) in EtOH (20 mL) was stirred at 25 °C for 16 hours and concentrated under reduced pressure, the residue was purified by silica column (PE/EtOAc = 20/1 to 2/1), the desired product was obtained as a colorless oil (1.60 g, 37.3% yield).

Intermediate **1-11**: 2-butyloctyl 10-oxodecanoate

To a solution of **1-10** (0.80 g, 2.24 mmol, 1.0 eq), $NaHCO_3$ (0.47 g, 5.60 mmol, 2.5 eq) in DCM (10 mL) was slowly added DMP (1.14 g, 2.69 mmol, 1.2 eq) at 0 °C. The reaction solution was stirred at 0 °C for 2 hours and quenched by aqueous saturated $Na_2S_2O_3$ (5 mL), then extracted with DCM (30 mL x 3), the organic layers were combined, dried over anhydrous $Na_2SO_4$ and concentrated unde reduced pressure, the residue was purified by silica column (PE/EtOAc = 1/0 to 4/1), the desired product was obtained as a colorless oil (570.0 mg, 71.6% yield).

Compound **1**: 2-butyloctyl 10-(9-((2-butyloctyl)oxy)-*N*-(3-(diethylamino)propyl)-9-oxononanamido)-11-(nonylamino)-11-oxoundecanoate

A solution of **1-11** (150.0 mg, 0.42 mmol, 1.0 eq), **1-12** (54.70 mg, 0.42 mmol, 1.0 eq) in MeOH (2 mL) was stirred at 25 ºC for 0.5 hour before **1-6** (0.18 g, 0.50 mmol, 1.2 eq) was added, the reaction soluton was stirred for another 0.5 hour at 25 ºC, then **1-3** (64.4 mg, 0.42 mmol, 1.0 eq) was added. After stirring at 25 ºC for 15 hours, the reaction solution was concentrated and diluted with $H_2O$ (30 mL), then extracted with EtOAc (10 mL x 3), the organic layers were combined, diried over ahydrous $Na_2SO_4$ and concentrated under reduced pressure, the residue was purified by silica column (DCM/MeOH = 1/0 to 20/1), the desired product was obtained as a colorless oil (164.1 mg, 40% yield). MS: m/z $[M+H]^+$ = 976.9. $^1H$ NMR (400 MHz, $CDCl_3$) $\delta$ 6.65 (s, 1H), 4.71 (s, 1H), 3.96 (d, $J$ = 4.0 Hz, 4H), 3.23-3.14 (m, 3H), 2.51-2.27 (m, 10H), 1.98-1.33 (m, 17H), 1.30-1.19 (m, 59H), 1.04-0.96 (m, 6H), 0.92-0.82 (m, 15H).

### EXAMPLE 2 Synthesis of Lipid-217

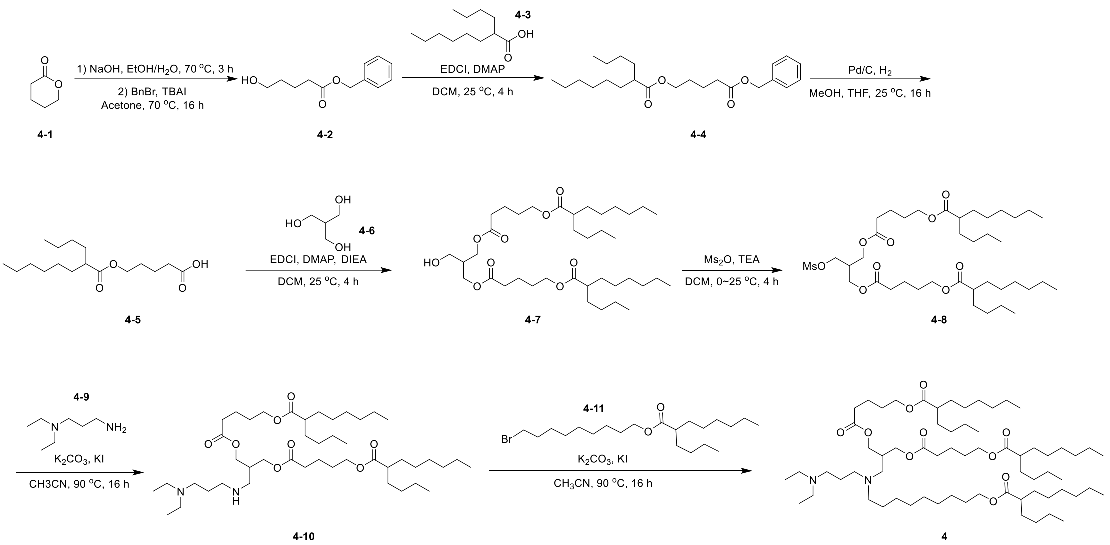


Intermediate **4-2**: benzyl 5-hydroxypentanoate

The solution of tetrahydro-2*H*-pyran-2-one (25.00 g, 249.70 mmol, 1.0 eq) and NaOH (10.99 g, 274.67 mmol, 1.1 eq) in $H_2O$ (20 mL) and ethanol (200 mL) was stirred at 70 °C for 3 hours and concentrated under reduced pressure. Then acetone (200 mL), TBAI (4.61 g, 12.48 mmol, 0.05 eq), BnBr (51.25 g, 299.64 mmol, 1.2 eq) was added, the resulting solution was stirred at 70 °C for 16 hours and diluted with $H_2O$ (500 mL), then extracted with EtOAc (250 mL x 2), the organic layers were combined and dried over anhydrous $Na_2SO_4$, then concentrated under reduced pressure, the residue was purified by silica column (PE/EtOAc = 10/1 to 2/1), the desired product was obtained as a colorless oil (37.00 g, 71.2% yield).

Intermediate **4-4**: 5-(benzyloxy)-5-oxopentyl 2-butyloctanoate

A solution of benzyl 5-hydroxypentanoate (37.00 g, 177.67 mmol, 1.0 eq), 2-butyloctanoic acid (35.59 g, 177.67 mmol, 1.0 eq), DMAP (21.70 g, 177.67 mmol, 1.0 eq) and EDCI (51.09 g, 266.50 mmol, 1.5 eq) in DCM (250 mL) was stirred at 25 °C for 4 hours. The reaction solution was diluted with $H_2O$ (500 mL) and extracted with DCM (200 mL x 2), the organic layers were combined and dried over anhydrous $Na_2SO_4$, then concentrated under reduced pressure, the residue was purified by silica column (PE/EtOAc = 20/1 to 4/1), the desired product was obtained as a colorless oil (64.00 g, 92.2% yield).

Intermediate **4-5**: 5-((2-butyloctanoyl)oxy)pentanoic acid

The solution of 5-(benzyloxy)-5-oxopentyl 2-butyloctanoate (64.00 g, 163.87 mmol, 1.0 eq) and Pd/C (3.49 g, 32.78 mmol, 0.2 eq, 10% purity) in MeOH (75 mL) and THF (75 mL) was stirred at 25 °C for 16 hours with a $H_2$ balloon. The reaction solution was filtered and concentrated under reduced pressure, the desired product was obtained as a colorless oil (45.00 g, 91.4% yield) which can be used for the next step without further purification.

Intermediate **4-7**: ((2-(hydroxymethyl)propane-1,3-diyl)bis(oxy))bis(5-oxopentane-5,1-diyl) bis(2-butyloctanoate)

A solution of 5-((2-butyloctanoyl)oxy)pentanoic acid (10.00 g, 33.29 mmol, 1.0 eq), 2-(hydroxymethyl)propane-1,3-diol (3.53 g, 33.29 mmol, 1.0 eq), DMAP (0.81 g, 6.66 mmol, 0.2 eq), EDCI (9.57 g, 49.94 mmol, 1.5 eq) and (8.60 g, 66.58 mmol, 2.0 eq) in DCM (100 mL) was stirred at 25 °C for 4 hours. The reaction solution was diluted with $H_2O$ (100 mL) and extracted with DCM (100 mL x 2), the organic layers were combined and dried over anhydrous $Na_2SO_4$, then concentrated under reduced pressure, the residue was purified by silica column (PE/EtOAc = 20/1 to 3/1), the desired product was obtained as a colorless oil (7.80 g, 69.9% yield).

Intermediate **4-8**: ((2-(((methylsulfonyl)oxy)methyl)propane-1,3-diyl)bis(oxy))bis(5-oxopentane-5,1-diyl) bis(2-butyloctanoate)

A solution of ((2-(hydroxymethyl)propane-1,3-diyl)bis(oxy))bis(5-oxopentane-5,1-diyl) bis(2-butyloctanoate) (3.90 g, 5.81 mmol, 1.0 eq), TEA (1.76 g, 17.43 mmol, 3.0 eq) in DCM (30 mL) was cooled in an ice bath, $Ms_2O$ (2.02 g, 11.62 mmol, 2.0 eq) was added dropwise. Then reaction solution was warmed and kept at 25 °C for 4 hours. The solution was diluted with $H_2O$ (30 mL) and extracted with DCM (50 mL x 2), the organic layers were combined and dried over anhydrous $Na_2SO_4$, then concentrated under reduced pressure, the residue was purified by silica column (PE/EtOAc = 50/1 to 5/1), the desired product was obtained as a colorless oil (3.85 g, 88.4% yield).

Intermediate **4-10**: ((2-(((3-(diethylamino)propyl)amino)methyl)propane-1,3-diyl)bis(oxy))bis(5-oxopentane-5,1-diyl) bis(2-butyloctanoate)

A solution of **4-8** (1.00 g, 1.34 mmol, 1.0 eq), *N*, *N*-diethylpropane-1,3-diamine (870.0 mg, 6.70 mmol, 5.0 eq), $K_2CO_3$ (560.0 mg, 4.02 mmol, 3.0 eq), KI (220.0 mg,

1.34 mmol, 1.0 eq) in $CH_3CN$ (20 mL) was stirred at 90 °C for 16 hours. The reaction solution was diluted with EtOAc (20 mL x 2), the organic layers were combined and dried over anhydrous $Na_2SO_4$, then concentrated under reduced pressure, the residue was purified by silica column (DCM/MeOH = 50/1 to 10/1), the desired product was obtained as a colorless oil (700.0 mg, 66.9% yield). MS: m/z $[M+H]^+$ = 783.6.

A solution of **4-10** (500.0 mg, 0.64 mmol, 1.0 eq), **4-11** (390.0 mg, 0.96 mmol, 1.5 eq), $K_2CO_3$ (270.0 mg, 1.92 mmol, 3.0 eq) and KI (110.0 mg, 0.64 mmol, 1.0 eq) in $CH_3CN$ (10 mL) was stirred at 90 °C for 16 hours. The reaction solution was diluted with $H_2O$ (20 mL) and extracted with EtOAc (20 mL x 2), the organic layers were combined and dried over anhydrous $Na_2SO_4$, then concentrated under reduced pressure, the residue was purified by silica column (DCM/MeOH = 50/1 to 20/1), the desired product was obtained as a colorless oil (132.8 mg, 18.8% yield). MS: m/z $[M+H]^+$ = 1107.9. $^1H$ NMR (300 MHz, $CDCl_3$) $\delta$ 4.15-4.01 (m, 10 H), 3.44-3.20 (m, 4H), 2.71-2.50 (m, 6H), 2.39-2.23 (m, 12H), 2.02-1.40 (m, 28H), 1.38-1.22 (m, 48H), 0.92-0.75 (m, 18 H).

### EXAMPLE 3 Synthesis of Lipid-306

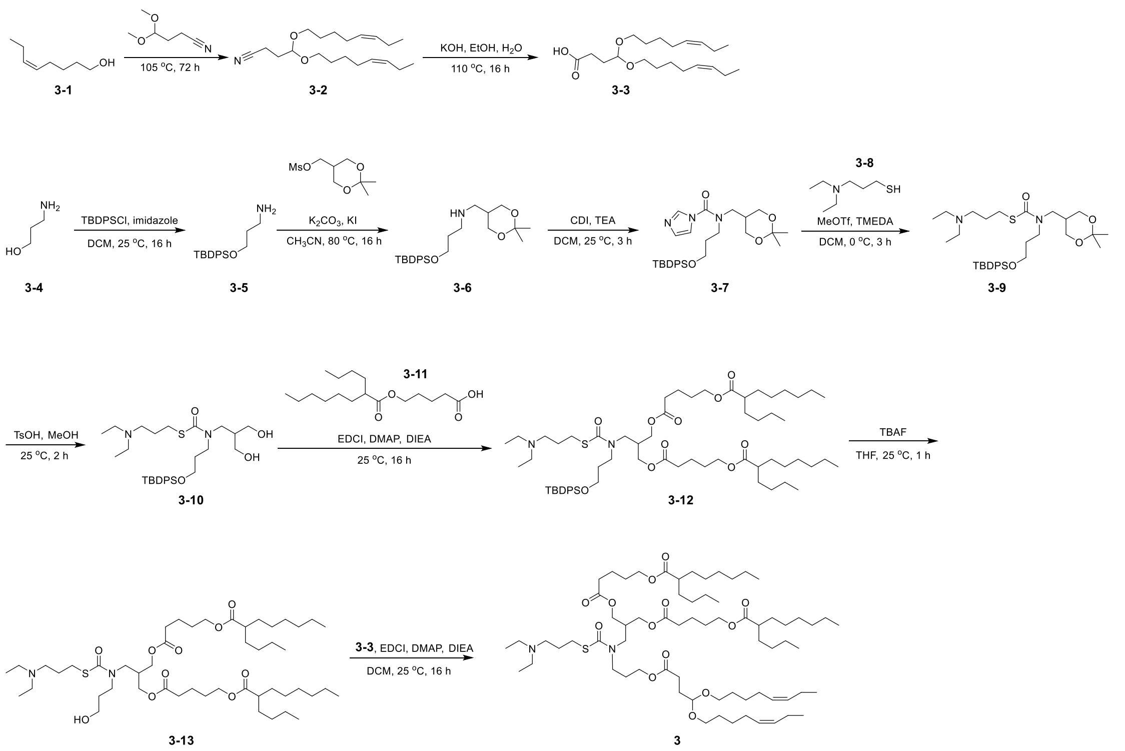


Intermediate **3-2**: 4,4-bis(((*Z*)-oct-5-en-1-yl)oxy)butanenitrile

A mixture of (*Z*)-oct-5-en-1-ol (49.63 g, 387.12 mmol, 2.0 eq), 4,4-dimethoxybutanenitrile (25.00 g, 193.56 mmol, 1.0 eq) and PPTS (2.43 g, 9.68 mmol, 2.0 eq) in sealed tube was stirred at 110 °C for 72 hours. The reaction mixture was diluted with $H_2O$ (200 mL) and extracted with EtOAc (200 mL x 3), the organic layers were combined and washed with aqueous saturated NaCl solution, dired over anhydrous $Na_2SO_4$, concentrated under reduced pressure. The residue was purified by silica column. 4,4-bis(((*Z*)-oct-5-en-1-yl)oxy)butanenitrile was obtained as a colorless oil (26.52 g, 41.8% yield). MS: m/z $[M+H]^+$ = 357.3. $^1$HNMR (400 MHz-$CDCl_3$) $\delta$ 5.42-5.28 (m, 4H), 4.57 (t, $J$ = 5.3 Hz, 1H), 3.61 (dt, $J$ = 9.3, 6.6 Hz, 2H), 3.44 (dt, $J$ = 9.3, 6.6 Hz, 2H), 2.42 (t, $J$ = 7.3 Hz, 2H), 2.12-1.90 (m, 9H), 1.66-1.54 (m, 5H), 0.97 (t, $J$ = 7.5 Hz, 6H).

Intermediate **3-3**: 4,4-bis(((*Z*)-oct-5-en-1-yl)oxy)butanoic acid

The solution of **3-2** (2.00 g, 6.22 mmol, 1.0 eq) and KOH (1.05 g,18.66 mmol, 3.0 eq) in EtOH (6 mL) and $H_2O$ (6 mL) was stirred at 110 °C for 16 hours in a sealed tube. The reaction mixture was concentrated, diluted with $H_2O$ and adjusted to pH = 5 with aqueous HCl (1 N). The resulting solution was extracted with EtOAc (50 mL x 3), the organic layers were combined and washed with aqueous saturated NaCl solution, dried over anhydrous

$Na_2SO_4$, then concentrated under reduced pressure. 4,4-bis(((*Z*)-oct-5-en-1-yl)oxy)butanoic acid was obtained as a yellow oil (1.50 g, 70.8% yield). $^1$H NMR (400 MHz, $CDCl_3$) $\delta$ 5.41-5.27 (m, 4H), 4.45 (t, *J* = 5.6 Hz, 1H), 3.51 (dt, *J* = 9.0, 6.7 Hz, 2H), 3.39 (dt, *J* = 9.0, 6.7 Hz, 2H), 2.43 (t, *J* = 7.2 Hz, 2H), 2.12-1.98 (m, 8H), 1.81 (q, *J* = 7.3 Hz, 2H) 1.65-1.49 (m, 4H), 1.44-1.32 (m, 4H), 0.94 (t, *J* = 7.5 Hz, 6H).

Intermediate **3-5**: 3-((*tert*-butyldiphenylsilyl)oxy)propan-1-amine

To a solution of 3-aminopropan-1-ol (5.00 g, 66.57 mmol, 1.0 eq) and imidazole (11.33 g, 166.42 mmol, 2.5 eq) in DCM (100 mL) was slowly added TBDPSCl (27.45 g, 99.85 mmol, 1.5 eq). The solution was stirred at 25 °C for 16 hours and concentrated under reduced pressure. The residue was purified by silical column, 3-((*tert*-butyldiphenylsilyl)oxy)propan-1-amine was obtained as a colorless oil (19.00 g, 91.0% yield). MS: m/z $[M+H]^+$ = 314.2.

Intermediate **3-6**: 3-((*tert*-butyldiphenylsilyl)oxy)-*N*-((2,2-dimethyl-1,3-dioxan-5-yl)methyl)propan-1-amine

The solution of (2,2-dimethyl-1,3-dioxan-5-yl)methyl methanesulfonate (3.00 g, 13.38 mmol, 1.0 eq), **3-5** (8.39 g, 26.76 mmol, 2.0 eq), $K_2CO_3$ (3.70 g, 26.76 mmol, 2.0 eq) and KI (2.22 g, 13.38 mmol, 1.0 eq) in $CH_3CN$ (50 mL) was stirred at 80 °C for 16 hours. The reaction mixture was concentrated and diluted with $H_2O$ (100 mL), then extracted with EtOAc (50 mL x 3), the organic layers were combined and wahsed with aqueous saturated NaCl solution, dried over anhydrous $Na_2SO_4$, then concentrated under reduced pressure. The residue was purified by silica column, 3-((*tert*-butyldiphenylsilyl)oxy)-*N*-((2,2-dimethyl-1,3-dioxan-5-yl)methyl)propan-1-amine was obtained as a colorless oil (4.70 g, 79.5% yield). MS: m/z $[M+H]^+$ = 442.3.

Intermediate **3-7**: *N*-(3-((*tert*-butyldiphenylsilyl)oxy)propyl)-*N*-((2,2-dimethyl-1,3-dioxan-5-yl)methyl)-1*H*-imidazole-1-carboxamide

To a solution of **3-6** (4.70 g, 10.64 mmol, 1.0 eq) and TEA (5.38 g, 53.20 mmol, 5.0 eq) in DCM (50 mL) was added CDI (5.18 g, 31.92 mmol, 3.0 eq) in batchs. The reaction mixture was stirred at 25 °C for 3 hours and extracted with DCM (50 mL x 3), the organic layers were combined, washed with aqueous saturated NaCl solution, dired over anhydrous $Na_2SO_4$ and concentrated under reduced pressure. The residue was purified by silica column, *N*-(3-((*tert*-butyldiphenylsilyl)oxy)propyl)-*N*-((2,2-dimethyl-1,3-dioxan-5-yl)methyl)-1H-imidazole-1-

carboxamide was obtained as a colorless oil (4.40 g, 77.2% yield). MS: m/z $[M+H]^+$ = 536.3.

Intermediate **3-9**: *S*-(3-(diethylamino)propyl) (3-((*tert*-butyldiphenylsilyl)oxy)propyl)((2,2-dimethyl-1,3-dioxan-5-yl)methyl)carbamothioate

MeOTf (0.61 g, 3.73 mmol, 1.0 eq) was slowly added to a solution of **3-7**(2.0 g, 3.73 mmol, 1.0 eq) in DCM (20 mL) at 0 ºC. The reaction solution was stirred at 0 ºC for 1 hour before TMEDA (1.30 g, 11.19 mmol, 3.0 eq) was added. Then **3-8** (0.82 g, 5.59 mmol, 1.5 eq) was added, the reaction solution was stirred at 0 ºC for another 2 hours and concentrated under reduced pressure. The residue was purified by silica column, *S*-(3-(diethylamino)propyl) (3-((*tert*-butyldiphenylsilyl)oxy)propyl)((2,2-dimethyl-1,3-dioxan-5-yl)methyl)carbamothioate (1.80 g, 78.5% yield). MS: m/z $[M+H]^+$ = 615.4.

Intermediate **3-10**: *S*-(3-(diethylamino)propyl) (3-((*tert*-butyldiphenylsilyl)oxy)propyl)(3-hydroxy-2-(hydroxymethyl)propyl)carbamothioate

The solution of **3-9** (0.90 g, 1.46 mmol, 1.0 eq) and PTSA (0.50 g, 2.92 mmol, 2.0 eq) in MeOH (10 mL) was stirred at 25 ºC for 2 hours. The reaction solution was concentrated under reduced pressure, *S*-(3-(diethylamino)propyl) (3-((*tert*-butyldiphenylsilyl)oxy)propyl)(3-hydroxy-2-(hydroxymethyl)propyl)carbamothioate was obtained as a yellow oil (0.84 g, 100% yield, crude) which can be used for the next step without further purification. MS: m/z $[M+H]^+$ = 575.3.

Intermediate **3-12**: 2-(10-butyl-3,9-dioxo-2,8-dioxahexadec-1-yl)-4-(2,2-dimethyl-3,3-diphenyl-4-oxa-3-silahept-7-yl)-10-ethyl-5-oxo-4,10-diaza-6-thiadodec-1-yl 5-[(2-butyl-1-oxooctyl)oxy]pentanoate

A mixture of **3-11**(1.32 g, 4.38 mmol, 3.0 eq), DIPEA (0.57 g, 4.38 mmol, 3.0 eq), DMAP (0.36 g, 2.92 mmol, 2.0 eq) and EDCI (0.84 g, 4.38 mmol, 3.0 eq) was stirred at 25 ºC for 0.5 hours before **3-10** (0.84 g, 1.46 mmol, 1.0 eq) was added. The reaction mixture was stirred at 25 ºC for 15.5 hours and extracted with DCM (25 mL x 3), the organic layers were combined and washed with aqueous saturated NaCl solution, dried over anhydrous $Na_2SO_4$, then concentrated under reduced pressure, the residue was purified by silica column, 2-(10-butyl-3,9-dioxo-2,8-dioxahexadec-1-yl)-4-(2,2-dimethyl-3,3-diphenyl-4-oxa-3-silahept-7-yl)-10-ethyl-5-oxo-4,10-diaza-6-thiadodec-1-yl 5-[(2-butyl-1-oxooctyl)oxy]pentanoate (1.40 g, 84.1% yield) was obtained as a coloerless oil. MS: m/z

$[M+H]^+ = 1139.8$.

Intermediate **3-13**: 2-(10-butyl-3,9-dioxo-2,8-dioxahexadec-1-yl)-10-ethyl-4-(3-hydroxypropyl)-5-oxo-4,10-diaza-6-thiadodec-1-yl 5-[(2-butyl-1-oxooctyl)oxy]pentanoate

TBAF (0.78 mL, 0.78 mmol, 1.0 M, 3.0 eq) was added to a solution of **3-12** (300.0 mg, 0.26 mmol, 1.0 eq) in THF (15 mL). The reaction solution was stirred at 25 °C for 1 hour and concentrated under reduced pressure. The residue was diluted with $H_2O$ (50 mL) and extracted with petrol ether (30 mL x 3), the organic layers were combined and washed with aqueous saturated NaCl solution, dried over anhydrous $Na_2SO_4$, then concentrated under reduced pressure. The residue was purified by silica column, 2-(10-butyl-3,9-dioxo-2,8-dioxahexadec-1-yl)-10-ethyl-4-(3-hydroxypropyl)-5-oxo-4,10-diaza-6-thiadodec-1-yl 5-[(2-butyl-1-oxooctyl)oxy]pentanoate was obtained as a colorless oil (110.0 mg, 46.4% yield). MS: m/z $[M+H]^+ = 901.6$. $^1H$ NMR (300 MHz, $CDCl_3$) $\delta$ 4.25-4.00 (m, 8H), 3.75-3.30 (m, 6H), 2.98-2.78 (m, 2H), 2.73-2.25 (m, 12H), 1.80-1.50 (m, 16H), 1.50-1.12 (m, 30H), 1.10-0.98 (m, 6H), 0.97-0.98 (m, 6H), 0.97-0.75 (m, 12H).

Compoud **3**: (24*Z*)-8-(10-butyl-3,9-dioxo-2,8-dioxahexadec-1-yl)-10-(6-ethyl-1-oxo-6-aza-2-thiaoct-1-yl)-18-{[(5*Z*)-oct-5-enyl]oxy}-5,15-dioxo-10-aza-6,14,19-trioxaheptacos-24-en-1-yl 2-butyloctanoate

The solution of **3-3** (0.14 g, 0.42 mmol, 1.5 eq), DIPEA (72.0 mg, 0.56 mmol, 2.0 eq), DMAP (51.0 mg, 0.42 mmol, 1.5 eq) and EDCI (81.0 mg, 0.42 mmol, 1.5 eq) was stirred at 25 °C for 0.5 hours before **3-13** (0.25 g, 0.28 mmol, 1.0 eq) was added. The resulting solution was stirred at 25 °C for 15.5 hours and extracted with DCM, washed with aqueous saturated NaCl, dried over anhydrous $Na_2SO_4$, then concentrated under reduced pressure. The residue was purified by silica column, (24*Z*)-8-(10-butyl-3,9-dioxo-2,8-dioxahexadec-1-yl)-10-(6-ethyl-1-oxo-6-aza-2-thiaoct-1-yl)-18-{[(5*Z*)-oct-5-enyl]oxy}-5,15-dioxo-10-aza-6,14,19-trioxaheptacos-24-en-1-yl 2-butyloctanoate was obtained as a colorless oil (130.0 mg, 38.3% yield). MS: m/z $[M+H]^+ = 1123.9$. $^1H$ NMR (300 MHz, $CDCl_3$) $\delta$ 5.47-5.25 (m, 4H), 4.55-4.48 (m, 1H), 4.20-4.01 (m, 8 H), 3.71-3.52 (m, 2H), 3.51-3.28 (m, 5H), 2.98-2.81 (m, 2H), 2.65-2.26 (m, 14H), 2.12-1.80 (m, 10H), 1.81-1.50 (m, 20H), 1.48-1.13 (m, 38H), 1.11-0.78 (m, 22H).

**EXAMPLE 4 Synthesis of Lipid-168**

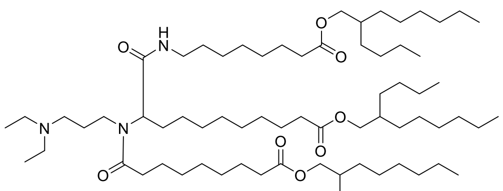

Intermediate **2-6**: 2-butyloctyl 8-isocyanooctanoate

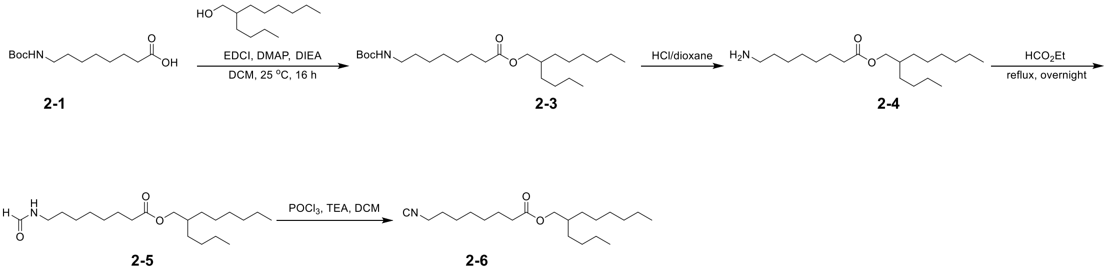


Intermediate **2-3**: 2-butyloctyl 8-((*tert*-butoxycarbonyl)amino)octanoate

The solution of **2-1** (8.02 g, 30.93 mmol, 1.5 eq), 2-butyloctan-1-ol (5.00 g, 20.62 mmol, 1.0 eq), DMAP (2.52 g, 20.62 mmol, 1.0 eq), DIEA (7.99 g, 61.86 mmol, 3.0 eq) and EDCI (5.93 g, 30.93 mmol, 1.5 eq) in DCM (60 mL) was stirred at 25 °C for 16 hours. The reaction mixture was concentrated under reduced pressure, the residue was purified by silica column. 2-butyloctyl 8-((*tert*-butoxycarbonyl)amino)octanoate was obtained as a colorless oil (7.20 g, 81.6% yield). MS: m/z $[M+H]^+$ = 428.4.

Intermediate **2-4**: 2-butyloctyl 8-aminooctanoate

The solution of **2-3** (7.20 g, 16.84 mmol, 1.0 eq) in HCl/dioxane (4 M, 40 mL) was stirred at 25 °C for 2 hours and concentrated under reduced pressure. The residue was diluted with aqueous saturated $NaHCO_3$ solution (100 mL) and extracted with DCM (100 mL x 3), the organic layers were combined and concentrated under reduced pressure, 2-butyloctyl 8-aminooctanoate was obtained as a colorless oil (5.00 g, 90.6% yield). MS: m/z $[M+H]^+$ = 328.3.

Intermediate **2-5**: 2-butyloctyl 8-formamidooctanoate

The solution of **2-4** (5.00 g, 15.27 mmol, 1.0 eq) in ethyl formate (50 mL) was reflux overnight and concentrated under reduced pressure. The residue was purified by silica column, 2-butyloctyl 8-formamidooctanoate was obtained as a colorless oil (2.63 g, 30.8% yield). MS: m/z $[M+H]^+$ = 356.3.

Intermediate **2-6**: 2-butyloctyl 8-isocyanooctanoate

Phosphorus oxychloride (1.71 g, 11.82 mmol, 1.6 eq) was added to the solution of **2-5** (2.63 g, 7.39 mmol, 1.0 eq) and (6.15 mL, 44.34 mmol, 6.0 eq) in THF (20 mL) slowly at 0 ºC. The reaction mixture was stirred for 3 hours and quenched by saturated aqueous $NaHCO_3$ (100 mL), then extracted with EtOAc (30 mL x 3). The organic layers were combined and dried over anhydrous $Na_2SO_4$, then concentrated under reduced pressure. The residue was purified by silica column, 2-butyloctyl 8-isocyanooctanoate was obtained as a colorless oil (1.45 g, 57.9% yield).

Compound **2** was prepared according to a similar Ugi procedure of compound **1** from intermediates **1-6**, **2-6**, **1-11** and *N*, *N*-diethylpropane-1,3-diamine. 39.8% yield. MS: m/z $[M+H]^+$ = 1161.0. $^1H$ NMR (400 MHz, $CDCl_3$) $\delta$ 6.65 (s, 1H), 4.69 (s, 1H), 3.96 (d, $J$ = 8.0 Hz, 5H), 3.38-3.12 (m, 4H), 2.53-2.26 (m, 11H), 1.99-1.89 (m, 1H), 1.77-1.54 (m, 16H), 1.47-1.39 (m, 3H), 1.33-1.17 (m, 71H), 1.02-0.98 (m, 6H), 0.93-0.84 (m, 18H).

| | Parameter | Value |
|---|---|---|
| 1 | **Sample name** | **Lipid-028** |
| 2 | Comment | Probe name: 4NUC<br>Field map 17Dec2019<br>rms diff/ rms err = 2.2192<br>Cycle number 8 out of 8<br><br>Estimated shim power dissipation = 5.8 W |
| 3 | Site | |
| 4 | Spectrometer | vnmrs |
| 5 | Solvent | cdcl3 |
| 6 | Temperature | 60.0 |
| 7 | Pulse Sequence | s2pul |
| 8 | Experiment | 1D |
| 9 | Spectrometer Frequency | 399.75 |
| 10 | Spectral Width | 8012.8 |
| 11 | Lowest Frequency | -801.0 |
| 12 | Nucleus | 1H |
| 13 | Acquired Size | 16384 |
| 14 | Spectral Size | 32768 |

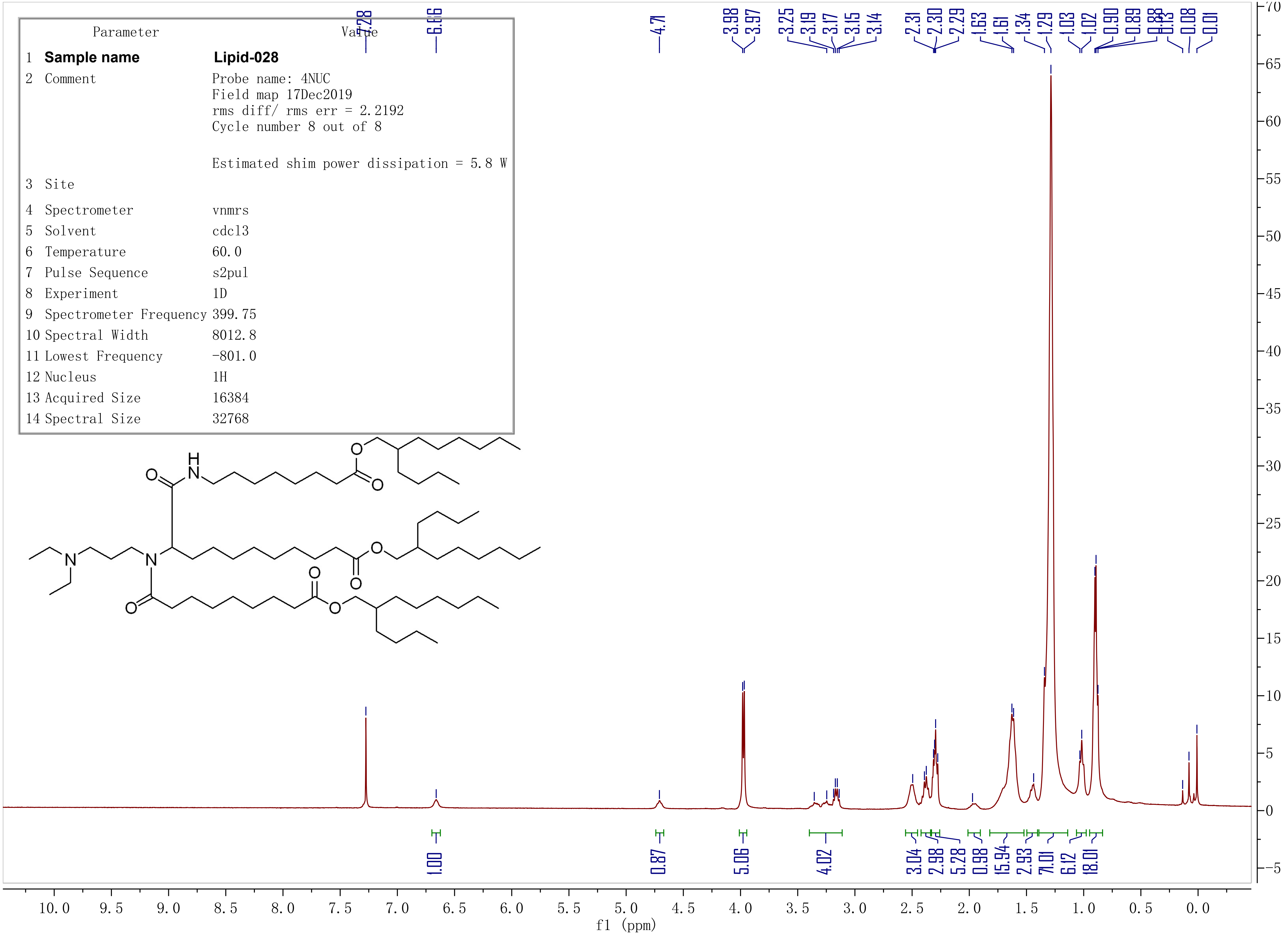

**Sample name: Lipid-306**
Probe name: 4NUC
Field map 17Dec2019
rms diff/rms err = 2.2192
Cycle number 8 out of 8

Estimated shim power dissipation = 5.8 W

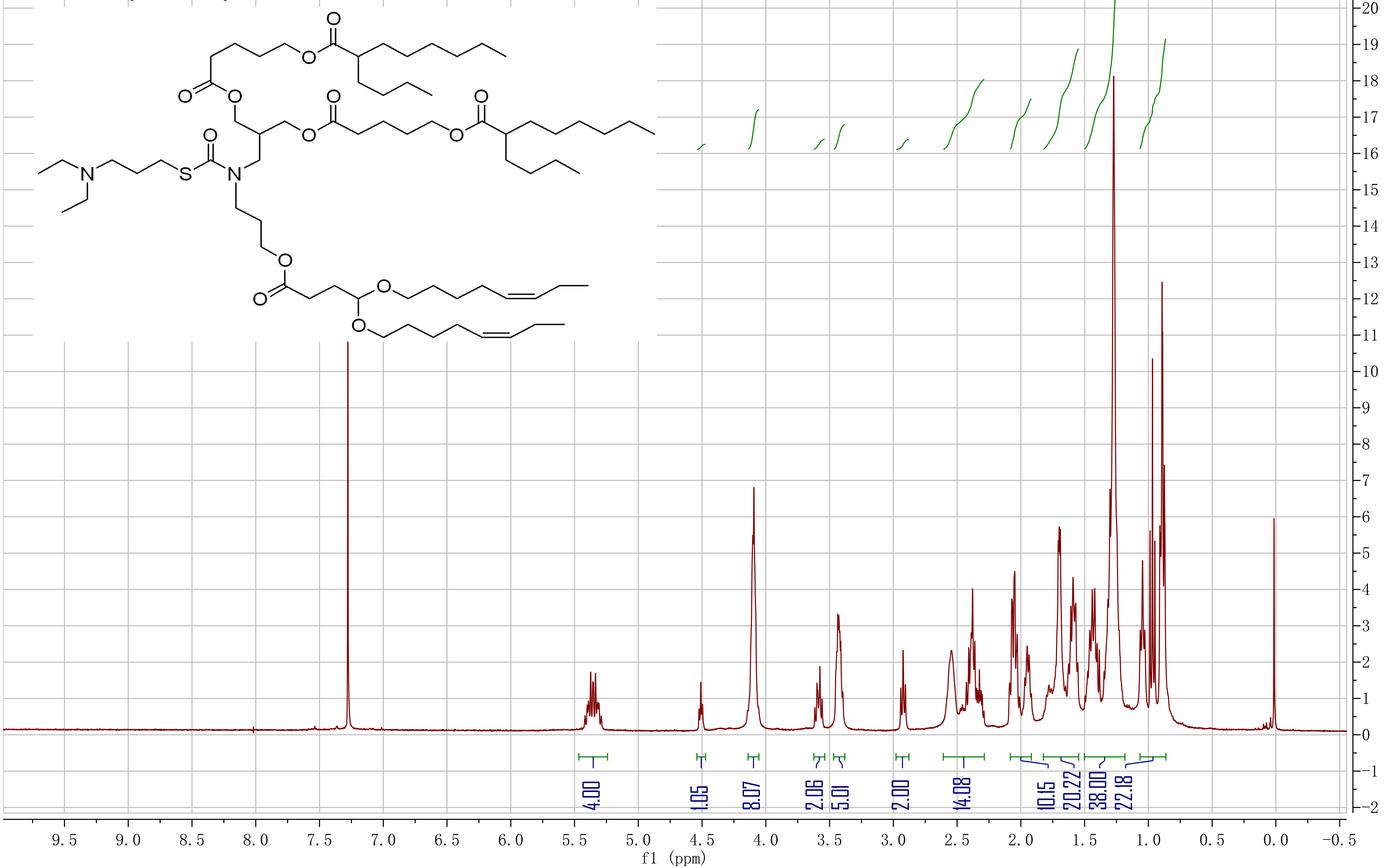

| | Parameter | Value |
|---|---|---|
| 1 | **Sample name** | **Lipid-217** |
| 2 | Comment | |
| 3 | Owner | admin |
| 4 | Site | |
| 5 | Spectrometer | QUANTUM-I |
| 6 | Author | |
| 7 | Solvent | CDCl3 |
| 8 | Temperature | 295.0 |
| 9 | Pulse Sequence | s1pul30 |
| 10 | Experiment | 1D |
| 11 | Spectrometer Frequency | 399.95 |
| 12 | Spectral Width | 8012.0 |
| 13 | Lowest Frequency | -1606.3 |
| 14 | Nucleus | 1H |
| 15 | Acquired Size | 32050 |
| 16 | Spectral Size | 65536 |

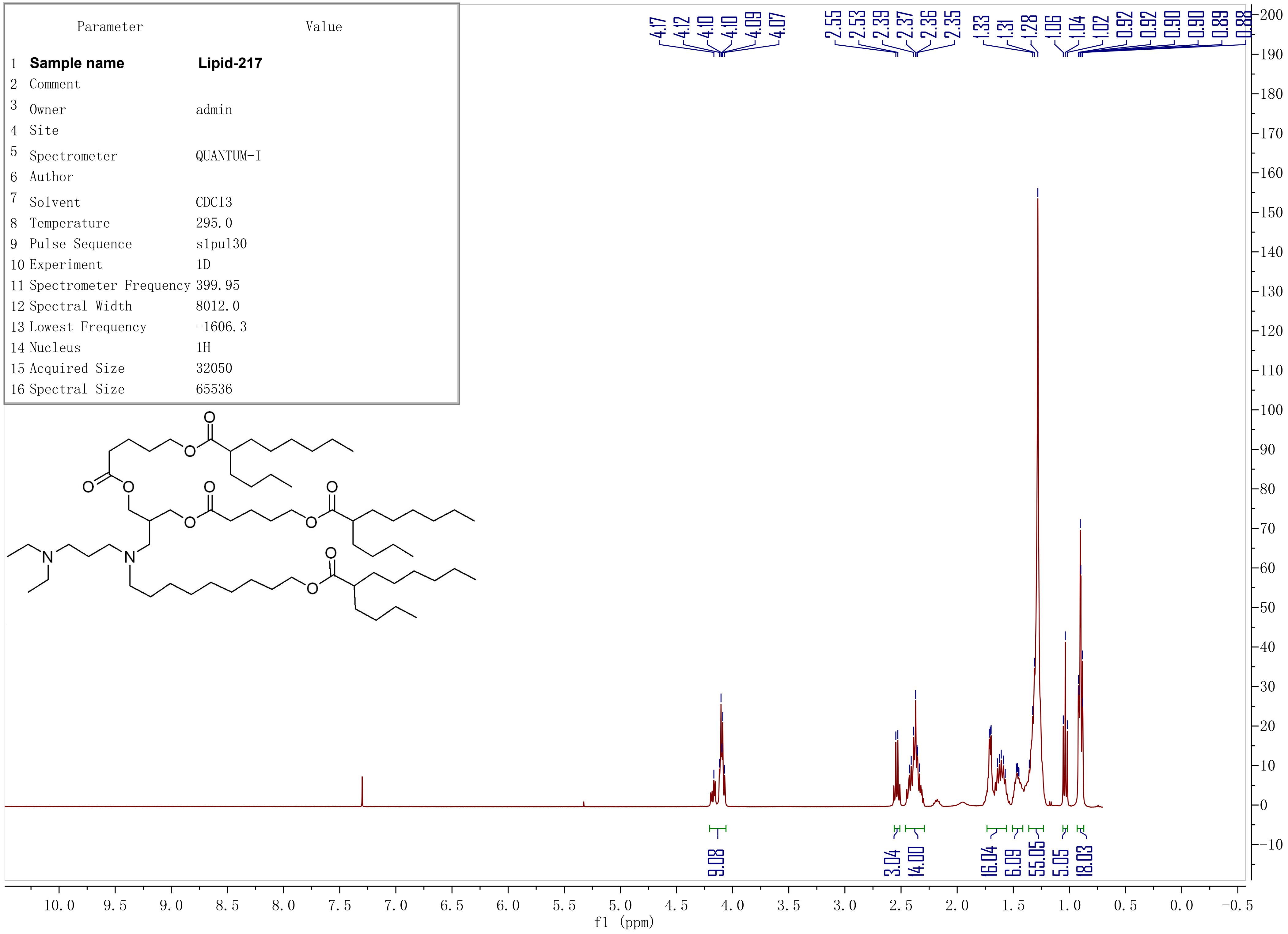

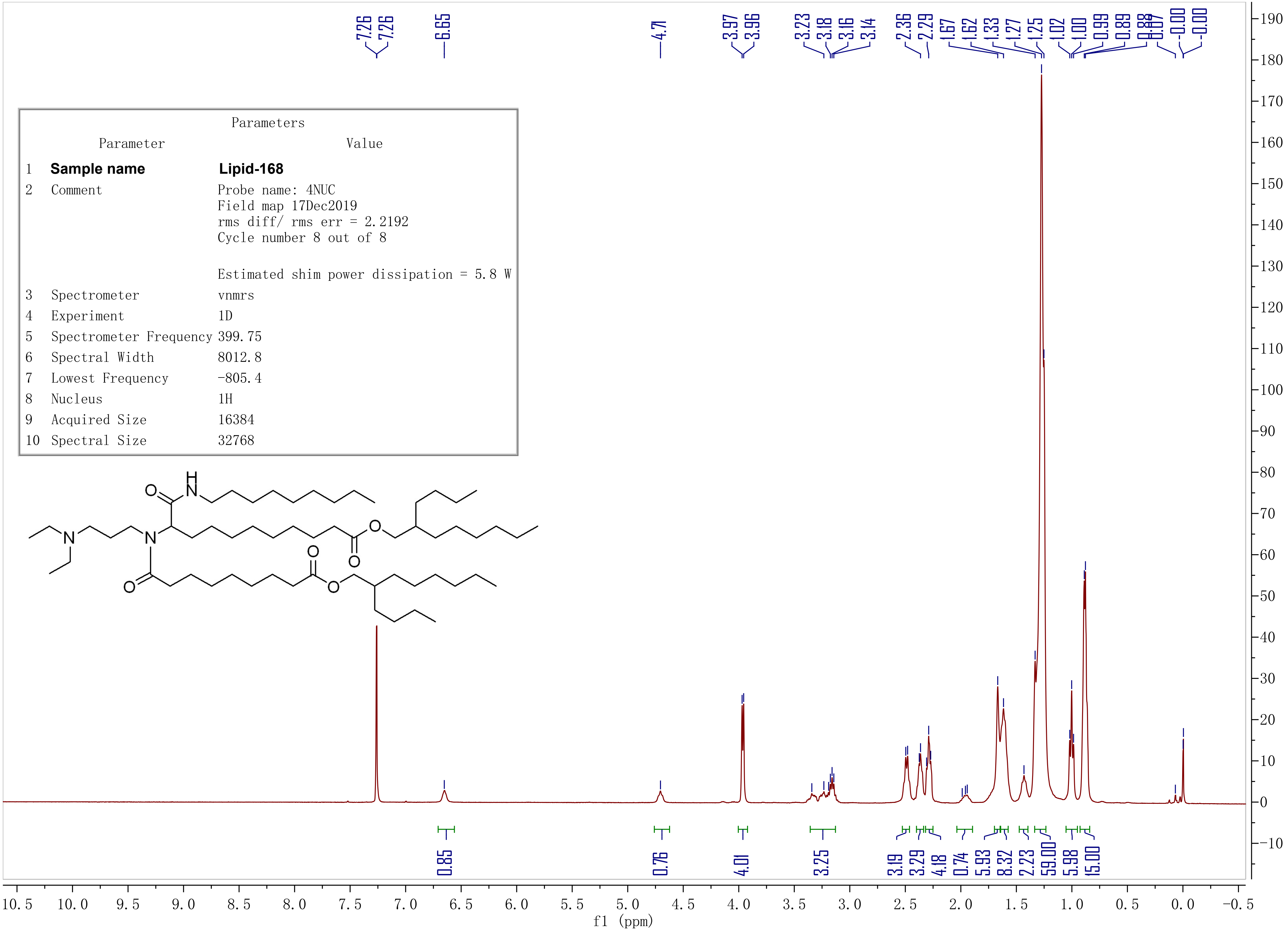

Parameters
Parameter Value
1 Sample name Lipid-168
2 Comment Probe name: 4NUC
Field map 17Dec2019
rms diff/ rms err = 2.2192
Cycle number 8 out of 8
Estimated shim power dissipation = 5.8 W
3 Spectrometer vnmrs
4 Experiment 1D
5 Spectrometer Frequency 399.75
6 Spectral Width 8012.8
7 Lowest Frequency -805.4
8 Nucleus 1H
9 Acquired Size 16384
10 Spectral Size 32768
7.26 7.26 6.65 4.71 3.97 3.96 3.23 3.18 3.16 3.14 2.36 2.29 1.67 1.62 1.33 1.27 1.25 1.02 1.00 0.99 0.89 0.88 0.87 -0.00 -0.00
0.85 0.76 4.01 3.25 3.19 3.29 4.18 0.74 5.93 8.32 2.23 59.00 5.98 15.00
f1 (ppm)
